# Autophagy initiation triggers p150^Glued^-AP-2β interaction on the lysosomes to facilitate their transport

**DOI:** 10.1101/2022.06.19.496704

**Authors:** Aleksandra Tempes, Karolina Bogusz, Agnieszka Brzozowska, Jan Weslawski, Matylda Macias, Oliver Tkaczyk, Aleksandra Lew, Malgorzata Calka-Kresa, Tytus Bernas, Andrzej A. Szczepankiewicz, Katarzyna Orzoł, Magdalena Mlostek, Shiwani Kumari, Magdalena Bakun, Tymon Rubel, Anna R. Malik, Jacek Jaworski

## Abstract

The endocytic adaptor protein 2 (AP-2) complex binds dynactin as part of its noncanonical function, which is necessary for dynein-driven autophagosome transport along microtubules in neuronal axons. The absence of this AP-2-dependent transport causes neuronal morphology simplification and neurodegeneration. The mechanisms that lead to formation of the AP-2–dynactin complex have not been studied to date. However, the inhibition of mammalian/mechanistic target of rapamycin complex 1 (mTORC1) enhances the transport of newly formed autophagosomes by influencing the biogenesis and protein interactions of Rab-interacting lysosomal protein (RILP), another dynein cargo adaptor. We tested effects of mTORC1 inhibition on interactions between the AP-2 and dynactin complexes, with a focus on their two essential subunits, AP-2β and p150^Glued^. We found that the mTORC1 inhibitor rapamycin enhanced p150^Glued^–AP-2β complex formation in both neurons and non-neuronal cells. Additional analysis revealed that the p150^Glued^–AP-2β interaction was indirect and required integrity of the dynactin complex. In non-neuronal cells rapamycin-driven enhancement of the p150^Glued^–AP-2β interaction also required the presence of cytoplasmic linker protein 170 (CLIP-170), the activation of autophagy, and an undisturbed endolysosomal system. The rapamycin-dependent p150^Glued^–AP-2β interaction occurred on lysosomal-associated membrane protein 1 (Lamp-1)-positive organelles but without the need for autolysosome formation. Rapamycin treatment also increased the acidification and number of acidic organelles and increased speed of the long-distance retrograde movement of Lamp-1-positive organelles. Altogether, our results indicate that autophagy regulates the p150^Glued^–AP-2β interaction, possibly to coordinate sufficient motor-adaptor complex availability for effective lysosome transport.

## Introduction

The effective cooperation of endomembrane components and the cytoskeleton is necessary for efficient intracellular communication and cell contacts with the extracellular environment. Microtubules are cytoskeleton elements that are essential for both the integrity of membrane compartments and their long-distance movement [1, 2]. Microtubules are dynamic and polarized, meaning that their ends (referred to as plus and minus) can undergo dynamic changes and are not identical [3]. This polarization determines the rules of directed cargo transport along microtubules by molecular motors, e.g., kinesins and dynein [4, 5]. The latter transports cellular cargo from the plus end to the minus end of microtubules [5–7]. Dynein does not act alone; it requires additional protein complexes to efficiently hold cargo and move along microtubules. One of these complexes is dynactin, a large multiprotein complex that initiates dynein movement, increases its processivity, and supports cargo attachment [5, 8, 9]. Dynactin consists of two major parts: sidearm and actin-related protein 1 (Arp-1) rod [5, 7, 10, 11]. The sidearm binds microtubules and dynein [5, 7, 10, 11]. The Arp-1 rod, in cooperation with dynein activators and adaptors, is responsible for cargo binding [5, 7, 10–15]. p150^Glued^ is part of the sidearm, the largest dynactin subunit, and a member of the microtubule plus-end tracking protein (+TIP) family [16]. Its binding to microtubule plus ends and its plus-end tracking behavior require the presence of cytoplasmic linker protein 170 (CLIP-170) [17, 18]. In some model systems (e.g., neuronal axons), this is essential for the initiation of dynein-dynactin-bound cargo transport along tyrosinated microtubules [8, 19].

The adaptor protein 2 (AP-2) complex consists of two large subunits (α, β), one medium subunit (µ), and one small subunit (σ) [20]. All four subunits contribute to the trunk of the AP-2 complex, but α and β2 C-termini project outside the trunk as α and β2 appendages (i.e., ears), respectively [20]. Canonically, AP-2 serves as a cargo adaptor complex in clathrin-mediated endocytosis [21]. However, evidence supports AP-2 functions outside the initiation of clathrin-mediated endocytosis, particularly in macroautophagy (hereafter called autophagy), lysosome tubulation, and microtubular transport [22–27]. The latter function was first discovered in neurons, in which AP-2 was found to be central to the retrograde transport of neuronal amphisomes that are produced by autophagosome-late endosome fusion and in axons act as signaling organelles that carry activated receptors for neurotrophins, such as tropomyosin receptor kinase B (TrkB) [24, 28] to the cell soma. The lack of this AP-2-dependent transport in axons resulted in disturbances in the morphology of dendrites and neurodegeneration [22, 24]. For dynein cargo adaptor function, AP-2 binds microtubule-associated protein 1A/1B-light chain 3 (LC3) on the amphisome surface via its AP-2µ subunit, whereas the AP-2β ear was shown to co-immunoprecipitate with p150^Glued^ [24].

Autophagy is a cellular process during which cells trap proteins or organelles (e.g., mitochondria) that are designated for degradation in double-membrane structures, called autophagosomes, and deliver them to lysosomes [29–32]. Autophagosome formation is a multistep process. Mammalian/mechanistic target of rapamycin complex 1 (mTORC1) is among its best-known regulators [33, 34]. Low mTORC1 activity allows autophagy initiation, but also autophagosome maturation, microtubular transport, and fusion with the lysosome [35–40]. Notably, in the case of neurons, the role of mTOR inhibition in autophagy initiation, particularly in axons, is still debated because of conflicting findings on whether rapamycin potentiates this process [41–44].

The effective termination of autophagy requires the fusion of autophagosomes or amphisomes with lysosomes, which contain degradative enzymes that are needed for autophagosome cargo destruction. It heavily relies on autophagosome and lysosome transport along microtubules. Dynein-dynactin transports autophagosomes retrogradely for fusion with lysosomes [45–47]. To meet autophagosomes, lysosomes may use both kinesins and dynein-dynactin [48, 49]. Lysosomes are dispersed through the cytoplasm with two distinguishable pools: perinuclear and peripheral [49]. The peripheral pool serves additional purposes (e.g., exocytosis [50]); when the demand for lysosomes greatly increases, however, such as during nutrient starvation that initiates autophagy, they move via dynein-dynactin transport toward autophagosomes that are already positioned in the cell center. To date, only two adaptors (ALG2 and JIP4) have been shown to recruit dynein-dynactin to lysosomes on demand upon nutrient starvation [51, 52].

Although AP-2–dynactin was shown to transport TrkB-positive amphisomes in neurons, unclear is whether the AP-2–dynactin complex also forms naturally in non-neuronal cells, in which amphisomes are considered very transient. Further details of the AP-2–dynactin interaction and its potential regulation are lacking. Our unpublished preliminary mass spectrometry data suggested a potential role for mTOR in the regulation of the p150^Glued^–AP-2β interaction. This is particularly intriguing when considering the important role of kinases in the regulation of microtubular transport [39, 53] and a recent finding that dynein can be recruited to autophagosomes by LC3 and Rab-interacting lysosomal protein (RILP) when mTORC1 activity is low [39]. Therefore, the present study investigated whether mTORC1 controls the p150^Glued^–AP-2 interaction and, if so, how and for what purpose. We found that mTORC1 inhibition enhanced the p150^Glued^–AP-2β interaction in both neurons and non-neuronal cells. We also found that p150^Glued^–AP-2β complex formation, boosted by mTORC1 inhibition, in non-neuronal cells required an intact dynactin complex and the undisturbed initiation of autophagy and endolysosomal pathway. We also found that the autophagy-induced p150^Glued^–AP-2β interaction occurred on lysosomes, which accelerated their retrograde motility. Thus, we revealed a novel mechanism whereby functions of essential components of cellular transport machinery are regulated at the level of autophagy initiation.

## Materials and Methods

### Plasmids and siRNA

The following plasmids were commercially available or described previously: pEGFPC1 (Clontech), β-actin-GFP and β-actin-tdTomato [54], HA-BirA [55], pAvi-tag-thrombin-HA (also known as Bio-Thrombin-HA; [56], pEGFPC1-Ap2b1 [57], pEGFPC2-Avi-tag-p150^Glued^ [24], pEGFPC2-Avi-tag-βGal (also known as bioβ-Gal) [58], EB3-GFP [59], pEGFP-CLIP-170 and pEGFPC1-CLIP-170-Δhead [60] (gift from Anna Akhmanova), pEGFPC1-p50 [61] (gift from Casper Hoogenraad), pEGFP-N1-Lamp1-GFP [62] (gift from Juan Bonifacino), pET-28-His_6_-AP-2β appendage domain [24] (gift from Volker Haucke), pGEX-4T1 (Merck, catalog no. GE28-9545-49), pGEX-4T1-GST-Eps15 (aa 541-790; gift from Mark McNiven) [63], and pGFP-LC3 [64] (gift from Iwona Ciechomska). Additional plasmids generated for this study are described in Supplementary Materials and Maethods in Supplementary Information. The following siRNAs were purchased from Invitrogen: Select Negative Control No. 1 siRNA (catalog no. 4390843; siCtrl), Silencer Select siRNA rat Clip1#1 (catalog no. 4390771, ID: s134775; siCLIP-70), Silencer Select siRNA human Clip1#1 (catalog no. 4392420, ID: s12372; siCLIP-170), Silencer Select siRNA rat Atg5 (catalog no. 4390771, ID: s172246), and Silencer Select siRNA rat Snap29 (catalog no. 4390771, ID: s138000).

### Antibodies

Commercially available primary antibodies that were used for this study are listed in Table S1. Rabbit anti-pan CLIP antibody (clone 2221; Western blot, 1:500) that recognized both cytoplasmic linker protein 115 (CLIP-115) and CLIP-170 was a gift from Casper Hoogenraad [65]. Alexa Fluor 488-, 568-, 594-, and 647-conjugated secondary antibodies (anti-mouse, anti-goat, and anti-rabbit) were obtained from Thermo Fisher. Horseradish peroxidase (HRP)-conjugated secondary antibodies were obtained from Jackson ImmunoResearch. Anti-mouse/anti-rabbit IRDye 680RD and IRDye 800CW were purchased from LI-COR Biosciences.

### Cell line and primary neuronal cultures and transfection

Rat2, HEK293, and primary hippocampal neurons cultures and transfections were performed using standard previously published protocols. For details, please refer to Supplementary Information.

### Drugs and drug treatment

All drugs, unless indicated otherwise, were dissolved in dimethylsulfoxide (DMSO), the final concentrations of which in the culture medium did not exceed 0.1%. For mTOR inhibition, Rat2 and HEK293T cells were treated with rapamycin (100 nM, Calbiochem, catalog no. 553210) or AZD-8055 (100 nM, Cayman Chemical, catalog no. 16978-5) for 2 h before the experiment (see figure legends for detailed descriptions). For translation inhibition, cells were treated for 2 h with 35 µM cycloheximide (Calbiochem, catalog no. 239763). For mTOR inhibition-dependent autophagy arrest at the initiation step, 25 µM SBI-0206965 (Merck, catalog no. SML1540) was used for 2.5 h (see figure legends for detailed descriptions). When combined with rapamycin, SBI-0206965 was added 30 min before the addition of rapamycin. To inhibit autophagic flux, cells were treated for 2 h with 60 µM 1-adamantyl(5- bromo-2-methoxybenzyl)amine (ABMA; Medchemexpress, catalog no. HY-124801) or 50 µM chloroquine (dissolved in water; Sigma-Aldrich, catalog no. C6628). For lysosomal vacuolar (H+)-adenosine triphosphatase (vATPase) inhibition, cells were treated for 2 h with bafilomycin A1 (Baf A1; 100 nM; Bioaustralis, catalog no. 88899-55-2). When cells were treated with an inhibitor and rapamycin, these compounds were administered at the same time and incubated for 2 h. For the alkalization of the cellular environment, 20 mM NH_4_Cl (dissolved in water; Sigma-Aldrich, catalog no. 213330) was added for 3 h. When cells were treated with both NH_4_Cl and rapamycin, NH_4_Cl was added to the cells 1 h prior to rapamycin, and the cells were then incubated with both drugs for an additional 2 h. To induce autophagy independently from mTORC1 inhibition, 100 μM L-690330 (Tocris, catalog no. 0681) was added to the cells for 3 h. Nocodazole (100 nM; Sigma-Aldrich, catalog no. M1404) was used to inhibit microtubule dynamics. For live-cell imaging experiments, the drug was added to Rat2 cells 1 h before imaging. For the PLA experiments, in which nocodazole was added to Rat2 cells alone or combined with rapamycin, it was added 2 h 15 min or 1 h before fixation, depending on whether it was used before or after rapamycin treatment. Before live imaging or fixation, neurons were treated with either vehicle (0.1% DMSO, 2 h) or rapamycin (100 nM, 2 h).

### Animals and rapamycin treatment

Rapamycin treatment and brain protein lysate isolation were performed according to a protocol that was approved by the 1st Ethical Committee in Warsaw (Poland; decision no. 843/2008 and 288/2012). Mature (3-month-old) male Wistar rats were used for the experiments. Rapamycin was initially dissolved in 100% ethanol at a 0.1 mg/ml concentration and stored at -20°C. Immediately before the injection, rapamycin was diluted in a vehicle solution that contained 5% Tween 80 and 5% PEG 400 (low-molecular-weight grade of polyethylene glycol; Sigma) and injected intraperitoneally (i.p.; 10 mg/kg) three times per week for 1 week. A control group of rats was injected with a vehicle solution that contained 5% Tween 80, 5% PEG 400, and 4% ethanol. Protein extraction from adult rat brains was performed as described previously [24] with a slight modification that involved the addition of 100 nM rapamycin to the lysis buffer for brains of rapamycin-treated animals.

### Proximity ligation assay and PLA-EM

Standard PLA procedures were performed as described previously [58] and detailed procedure is described in Supplementary Information. For PLA-EM, Rat2 cells were grown for 24 h and fixed for 15 min in 4% PFA with the addition of 0.1% glutaraldehyde in PBS. The cells were then washed three times with PBS. Afterward, cells in PBS were permeabilized by three cycles of freezing in liquid nitrogen, thawed, incubated for 20 min with 1% sodium borohydride in PBS, washed three times with PBS, incubated for 20 min at room temperature with 3% hydrogen peroxide in PBS/ethanol (1:1), and washed again three times in PBS. Next, fixed and permeabilized cells were incubated for 1 h in 5% bovine serum albumin (BSA) in PBS at room temperature. Afterward, the cells were incubated for 48 h at 4°C with primary mouse anti-p150^Glued^ and rabbit anti-AP-2β antibodies that were diluted in 0.1% donkey serum/PBS and washed three times in PBS at room temperature. The cells were then incubated for 60 min at 37°C with PLA probes (Sigma-Aldrich, catalog no. DUO92002 and DUO92004) and washed twice for 5 min with buffer A (Sigma-Aldrich, catalog no. DUO82046). Ligation and amplification were performed according to the manufacturer’s protocol using DuoLink In Situ Detection Reagents Brightfield (Sigma-Aldrich, catalog no. DUO92012). After the PLA reaction, the cells were additionally fixed with 2.5% glutaraldehyde in PBS for 2 h at 4°C and washed twice with PBS and once with deionized water. Next, the cells were incubated with 3% hexamethylenetetramine, 5% silver nitrate, and 2.5% disodium tetraborate for 10 min at 60°C, washed three times with water, once with 0.05% tetrachloroauric acid, once with 2.5% sodium thiosulfate, and finally three times with water (all at room temperature). As the last step, the cells were postfixed with 1% osmium tetroxide for 1 h at room temperature, washed with water, incubated in 1% aqueous uranyl acetate for 1 h, dehydrated with increasing dilutions of ethanol, and infiltrated with epoxy resin (Sigma-Aldrich, catalog no. 45-359-1EA-F). After resin polymerization at 60°C, fragments of coverslips with embedded cells were cut out with scissors and glued to the resin blocks. The blocks were then trimmed and cut with a Leica ultramicrotome (Ultracut R) to obtain ultrathin sections (70 nm thick) and collected on 100 mesh copper grids (Agar Scientific, catalog no. AGS138-1). Specimen grids were examined with a Tecnai T12 BioTwin transmission electron microscope (FEI) that was equipped with a 16 megapixel TemCam-F416 (R) camera (Tietz Video and Imaging Processing Systems).

### Immunofluorescence and fixed cell image acquisition and analysis

Procedures used for immunofluorescence and fixed cell image acquisition and analysis are described in detail in Supplementary Information.

### Live imaging of microtubule dynamics

For microtubule dynamics analysis, Rat2 cells were electroporated with pEGFP-EB3 or pEGFP-CLIP-170 plasmids. Forty-eight hours later, time-lapse movies of EGFP-EB3 or EGFP-CLIP-170 comets were taken using an Andor Revolutions XD spinning disc microscope with a 63× objective and 1.6 x tube lens (optovar) at 1004 × 1002 pixel resolution. Images were taken with an exposure of 200 ms and interval of 0.3 s, collecting a total of 600 frames over 3 min. During imaging, cells were kept in a Chamlide magnetic chamber (Quorum Technologies) at 37°C with 5% CO_2_ in the incubator that was part of the microscope system. Only EGFP-EB3 or EGFP-CLIP-170 comets whose movements lasted at least four consecutive frames and had a displacement length of at least 10 pixels (0.68 µm) were analyzed using the ImageJ “TrackMate” plugin [66]. The reported values are the number of tracks (i.e., the quantification of objects that were detected as comets), the total run length of comets before catastrophe (*Track Displacement*), comet lifetime (*Track Duration*), and *Track Mean Speed*. For the analysis, values were calculated as means for each cell.

### Live imaging of Lamp-1-GFP objects

For the lysosomal-associated membrane protein 1 (Lamp-1) object motility analysis, Rat2 cells were electroporated with Lamp-1-GFP plasmid and imaged approximately 22 h later with an Andor Revolutions XD spinning disc microscope, with the same setup and settings as described above. Time-lapse movies were collected over 3 min at 0.3 s intervals, resulting in 600 frames. Movies were analyzed using the ImageJ “TrackMate” plugin [66]. Only objects that were visible for more than 4 consecutive frames were considered. Objects with movement lengths shorter than 6.8 µm were considered not motile. The number of motile and non-motile objects divided by cell area and their ratio were measured. Other calculated values that were used for the analysis included the following: length of the Lamp-1 object run (*Track Displacement*), time during which the objects were visible (*Track Duration*), and speed with which the object moved (*Track Mean Speed*). For the analysis of Lamp-1-GFP objects in the cell center and in the periphery, Lamp-1-GFP objects were assigned manually to these compartments based on maximum projections.

### Live imaging of neurons

An Andor Revolutions XD confocal spinning disc microscope and a Chamlide magnetic chamber were used for the *in vivo* imaging of cells. Cell recording was performed at 37°C with 5% CO_2_ in a thermostat-controlled incubator. Series of images were acquired at 502 x 501 pixel resolution, with 1 s interval between frames. The total imaging time was 3 min. The images were collected using the 63x objective and 1x optovar. Neuronal processes were manually tracked with ImageJ. Based on these tracks, kymographs were created in ImageJ with the “Kymograph Clear” plugin [67]. By tracing lines on the kymographs, the number of particles that moved at a given time and distance was determined, and their velocity was calculated by determining the difference between the height of the starting point and end point of a given particle.

### Biochemical procedures

All standard biochemical procedures including protein production, pull-down assays, immunoprecipitation, Western blot, kinase assays and RNA isolation and Quantitative Real-Time PCR are described in details in Supplementary Information.

### Statistical analysis

The exact numbers of cells (*n*) that were examined for the respective experiments and number of repetitions of each experiment (*N*) are provided in the figure legends. The statistical analyses were performed using GraphPad Prism 9 software or Rstudio. The Shapiro-Wilk test was used to assess whether the data distribution met the assumptions of a normal distribution. For comparisons between two groups, the *t*-test (in the case of a normal distribution) or Mann-Whitney test (in the case of a non-normal distribution) was used to verify statistical significance. For comparisons between more than two groups, the data were analyzed using one-way analysis of variance (ANOVA), followed by the Bonferroni multiple-comparison *post hoc* test (in the case of a normal distribution) or Kruskal-Wallis test and Dunn’s multiple-comparison *post hoc* test (in the case of a non-normal distribution). For comparisons between two factors, the data were analyzed using two-way ANOVA, followed by Tukey’s multiple-comparison *post hoc* test. The statistical significance of qRT-PCR data was assessed with the one-sample Student’s *t*-test.

## Results

### mTORC1 inhibition increases p150^Glued^–AP-2β interaction in neurons and non-neuronal cells

Recent work shows that mTOR inhibition increases the biosynthesis of RILP and enhances its recruitment to autophagosomes, potentiating their transport in different cell types, including neurons [39]. This finding prompted us to investigate whether rapamycin, an inhibitor of mTORC1, also affects the AP-2–dynactin interaction that is responsible for the axonal transport of amphisomes [24]. We performed the IP of endogenous AP-2β from brain lysates from control rats and rats that were treated with rapamycin for 8 days. Rapamycin treatment effectively decreased the phosphorylation of ribosomal protein S6 (P-S6) at Ser235/236, confirming efficient mTORC1 inhibition, and increased the co-IP of p150^Glued^ with AP-2β. We observed no noticeable difference in overall levels of p150^Glued^ or AP-2β in corresponding input fractions (**Fig. 1A**).

**Fig. 1.**
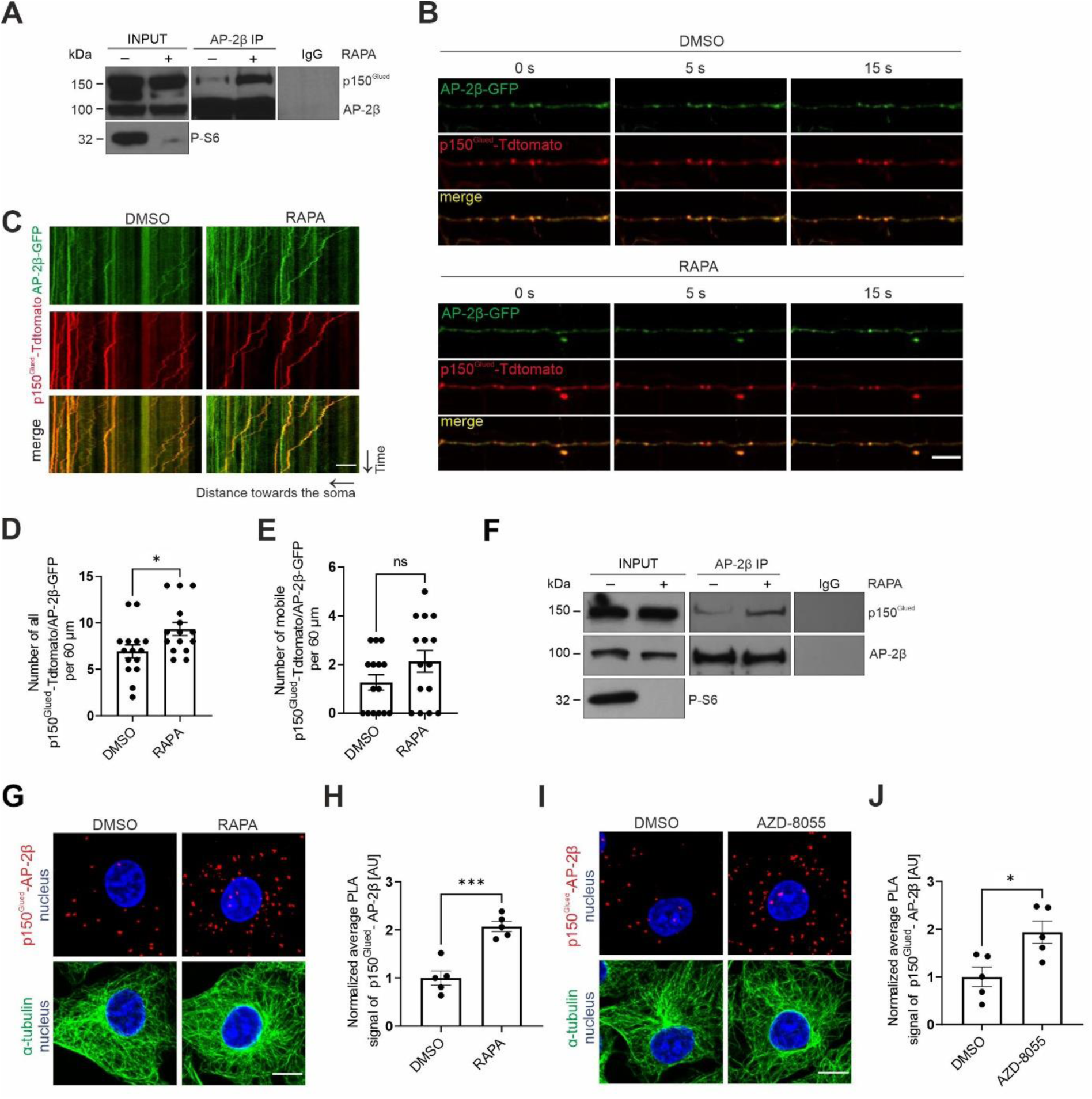
mTORC1 inhibition increases p150^Glued^–AP-2β interaction. (**A**) Western blot analysis of levels of endogenous p150^Glued^, AP-2β, and P-S6 (Ser235/236) and co-immunoprecipitation of endogenous AP-2β with p150^Glued^ from brain lysates from rats that were treated with rapamycin (RAPA+) or vehicle (RAPA-). INPUT, 10% of lysate used for immunoprecipitation. Shown is a representative example from *N =* 3 independent experiments. (**B-E**) Dynamics of p150^Glued^-Tdtomato and AP-2β-GFP co-transport in axons of neurons that were treated for 2 h with 0.1% DMSO or 100 nM rapamycin (RAPA). (**B**) Representative snapshots of 60 μm segment of axon and (**C**) corresponding kymographs of p150^Glued^-Tdtomato- and AP-2β-GFP-positive objects. See also Supplementary Fig. S1 and Movies 1-2. Scale bar = 10 μm. (**D**) Number of all p150^Glued^-tdTomato/AP-2β-GFP objects in axons per 60 μm. The data are expressed as the mean ± SEM. *N* = 4 independent experiments. *n* = 15 cells per variant. *\*p* < 0.05 (Mann-Whitney test). (**E**) Number of mobile p150^Glued^-Tdtomato/AP-2β-GFP-positive objects in axons per 60 μm. The data are expressed as the mean *±* SEM. *N =* 4 independent experiments. *n* = 15 cells per variant. *ns*, nonsignificant (Mann-Whitney test). (**F**) Western blot analysis of levels of endogenous p150^Glued^, AP-2β, and P-S6 (Ser235/236) and co-immunoprecipitation of endogenous p150^Glued^ with AP-2β from HEK293T cells that were treated for 2 h with 0.1% DMSO (RAPA-) or 100 nM rapamycin (RAPA+). Input, 10% of lysate used for immunoprecipitation. Shown is a representative example from *N* = 5 independent experiments. (**G**) Representative images of Rat2 fibroblasts that were treated for 2 h with 0.1% DMSO or 100 nM rapamycin (RAPA) with p150^Glued^–AP-2β PLA signals (red), immunofluorescently labeled tubulin (green), and DAPI-stained nuclei (blue). Scale bar = 10 μm. (**H**) Quantification of the number of p150^Glued^–AP-2β PLA puncta in cells that were treated as in G. The data are expressed as the mean number of PLA puncta per cell, normalized to the control variant (DMSO) *±* SEM. *N* = 5 independent experiments. *n =* 201 cells for each experimental variant*. ***p* < 0.001 (Student’s *t*-test). (**I**) Representative images of Rat2 fibroblasts that were treated for 2 h with 0.1% DMSO or 100 nM AZD-8055 with p150^Glued^– AP-2β PLA signals (red), immunofluorescently labeled tubulin (green), and DAPI-stained nuclei (blue). Scale bar = 10 μm. (**J**) Quantification of the number of p150^Glued^–AP-2β PLA puncta in cells that were treated as in I. The data are expressed as the mean number of PLA puncta per cell, normalized to the control variant (DMSO) ± SEM. *N* = 5 independent experiments. *n* = 211 cells (DMSO), 188 cells (AZD-8055). *\*p* < 0.05 (Student’s *t*-test).

mTORC1 inhibition potentiates the AP-2–dynactin interaction in the brain. Therefore, we next tested effects of rapamycin on the AP-2β and p150^Glued^ interaction in axons of hippocampal neurons, in which its functional significance was demonstrated [24]. We transfected neurons that were cultured *in vitro* (DIV5) with plasmids that encoded tdTomato-tagged p150^Glued^ and green fluorescent protein (GFP)-tagged AP-2β. Two days later, we imaged the behavior of fluorescently tagged proteins in axons in DMSO-treated (control) cells and cells that were treated with 100 nM rapamycin for 2 h (**Fig. 1B-E**, **Movie 1-4**, **Fig. S1A [Supplementary Figures in Supplementary Information]**). Similar to brain tissues, rapamycin decreased P-S6 levels (**Fig. S1B, C**). We also observed a significant increase in the number of p150^Glued^–AP-2β-positive objects (**Fig. 1D**), suggesting that mTORC1 inhibition boosted the p150^Glued^–AP-2β interaction in neurons, but these objects were largely immobile. The difference between fractions of mobile AP-2β/p150^Glued^-positive objects was also statistically nonsignificant under the tested conditions (**Fig. 1E**). Thus, we concluded that mTORC1 inhibition in neurons potentiated the p150^Glued^–AP-2β interaction in axons.

We next investigated whether mTORC1 inhibition has a similar effect on the co-occurrence of p150^Glued^ and AP-2β in non-neuronal cells using HEK293T and Rat2 cell lines. Under basal culture conditions, there was some evidence of an AP-2–dynactin interaction, demonstrated by IP (HEK293T), immunofluorescence co-localization (Rat2), and the PLA (Rat2; **Fig. 1F-J; Fig. S2**). The treatment of HEK293T cells with 100 nM rapamycin for 2 h decreased P-S6 levels and increased AP-2β–p150^Glued^ (**Fig. 1F**). In Rat2 cells, rapamycin decreased S6 phosphorylation (**Fig. S2A**) and enhanced the p150^Glued^–AP-2β interaction, measured by immunofluorescence signal co-localization analysis and PLA using antibodies against endogenous proteins (**Fig. 1G, H**, **Fig. S2B-G**). Co-localization analysis showed relatively low co-localization under basal conditions and an increase in rapamycin-treated cells (**Fig. S2B-F**). The PLA results additionally confirmed the biochemical and immunofluorescence evidence that is described above (**Fig. 1G, H, Fig. S2G**). AZD-8055, an ATP-competitive inhibitor of mTOR (2 h, 100 nM), also effectively decreased P-S6 immunofluorescence (**Fig. S2A**) and increased the p150^Glued^–AP-2β PLA signal in Rat2 cells (**Fig. 1I, J**), supporting our hypothesis that mTOR inhibition enhances the studied interaction.

The best-known function of mTORC1 is the positive regulation of protein synthesis. Therefore, we treated Rat2 cells for 2 h with cycloheximide (CHX) (35 µM), a widely used protein synthesis inhibitor, to test whether the increase in the PLA signal was attributable to a decrease in translation. However, we did not detect a significant difference in the PLA signal in CHX-treated cells compared with control cells (**Fig. S3**). Overall, our results suggest that mTORC1 inhibition enhances the interaction between p150^Glued^ and AP-2β in different cell types but not because of the canonical function of mTORC1 as a translational enhancer.

Excluding the possibility that protein synthesis inhibition was a main driver of an increase in the p150^Glued^–AP-2β interaction prompted us to test whether p150^Glued^ or AP-2β are substrates of mTOR. *In vitro* kinase assays, using GFP-AP-2β or GFP-p150^Glued^ and the mTOR active fragment, excluded such a possibility (**Fig. S4A**). Furthermore, inspection of the available datasets of mTOR-dependent phosphoproteomes ([e.g., 68, 69]) did not support the mTOR-dependent phosphorylation of other dynactin or AP-2 subunits. mTOR inhibition in HEK293T cells also did not affect the connection between the dynactin sidearm and its Arp1 rod, indicated by the lack of differences in the co-IP of p150^Glued^ with p62 and Arp1 between analyzed conditions (**Fig. S4B**). Thus, we concluded that the observed effects of mTORC1 inhibition on the p150^Glued^–AP-2β interaction were not driven by direct actions of mTOR on the dynactin complex or AP-2β.

### p150^Glued^ interaction with AP-2β is indirect and requires dynactin integrity

Our previous study [24] and the data described above show that AP-2 and p150^Glued^ can form a complex, but further characterization is needed. Therefore, in the following experiments, we first focused on the biochemical characterization of this interaction. Using an Avi-tag pull-down assay, we previously demonstrated that full-length p150^Glued^ that is produced in HEK293T cells can effectively bind the *E. coli*-produced β2 ear of AP-2β [24]. Therefore, we used this system to characterize the p150^Glued^–β2-ear interaction further and clarified which p150^Glued^ domains are required. We first compared the ability of the N-terminal (1-490 aa; N) and C-terminal (490-end; C) parts of p150^Glued^ and the full-length protein (**Fig. 2A**) that is produced in HEK293T cells to bind the His-tagged β2 ear. The C-terminal part of p150^Glued^ was as effective as the full-length protein, whereas its N-terminus did not bind the AP-2 fragment, exactly like β-galactosidase that served as the negative control (**Fig. 2B**). Moreover, a shorter fragment of the C-terminus (1049-end; C2; **Fig. 2A**), which is known for its contribution to the dynactin interaction with cargo adaptors [13, 70, 71], also bound the β2 ear (**Fig. 2C**). Notably, however, the newest structural and biochemical data raise the issue of whether the C-terminus of p150^Glued^ binds cargo adaptors directly [10, 11]. Indeed, when both C-terminal fragments of p150^Glued^ were produced in *E. coli,* no interaction with the AP-2β fragment was observed (**Fig. 2D, E**). But, the β2 ear interacted with Eps15 protein (541-790 aa fragment fused to GST; [63, 72]), which was used as a positive control. The most C-terminal part of p150^Glued^ is known for its role in connecting the dynactin sidearm with the Arp-1 rod that binds cargo [73]. Indeed, Western blot indicated that during the p150^Glued^ Avi-tag pull-down, regardless of the harsh washing conditions, the dynactin Arp1-rod proteins (Arp1 and p62) also co-purified from HEK293T cells (**Fig. 2F**). This suggests that intact dynactin might be involved in formation of the multi-protein complex that contains p150^Glued^ and AP-2β. Indeed, overexpression of the p50 subunit of dynactin, which is routinely used to disrupt dynactin complex integrity by dissociation of the sidearm and Arp1 rod [74, 75]; **Fig. 2G**), completely blocked the rapamycin-driven increase in the p150^Glued^-AP-2β PLA signal in Rat2 cells (**Fig. 2H, I**). Thus, although p150^Glued^ and AP-2β are unlikely to bind each other directly, their interaction (evident upon mTORC1 inhibition) requires the C-terminus of p150^Glued^ and an undisturbed interaction between the dynactin sidearm and Arp-1 rod.

**Fig. 2.**
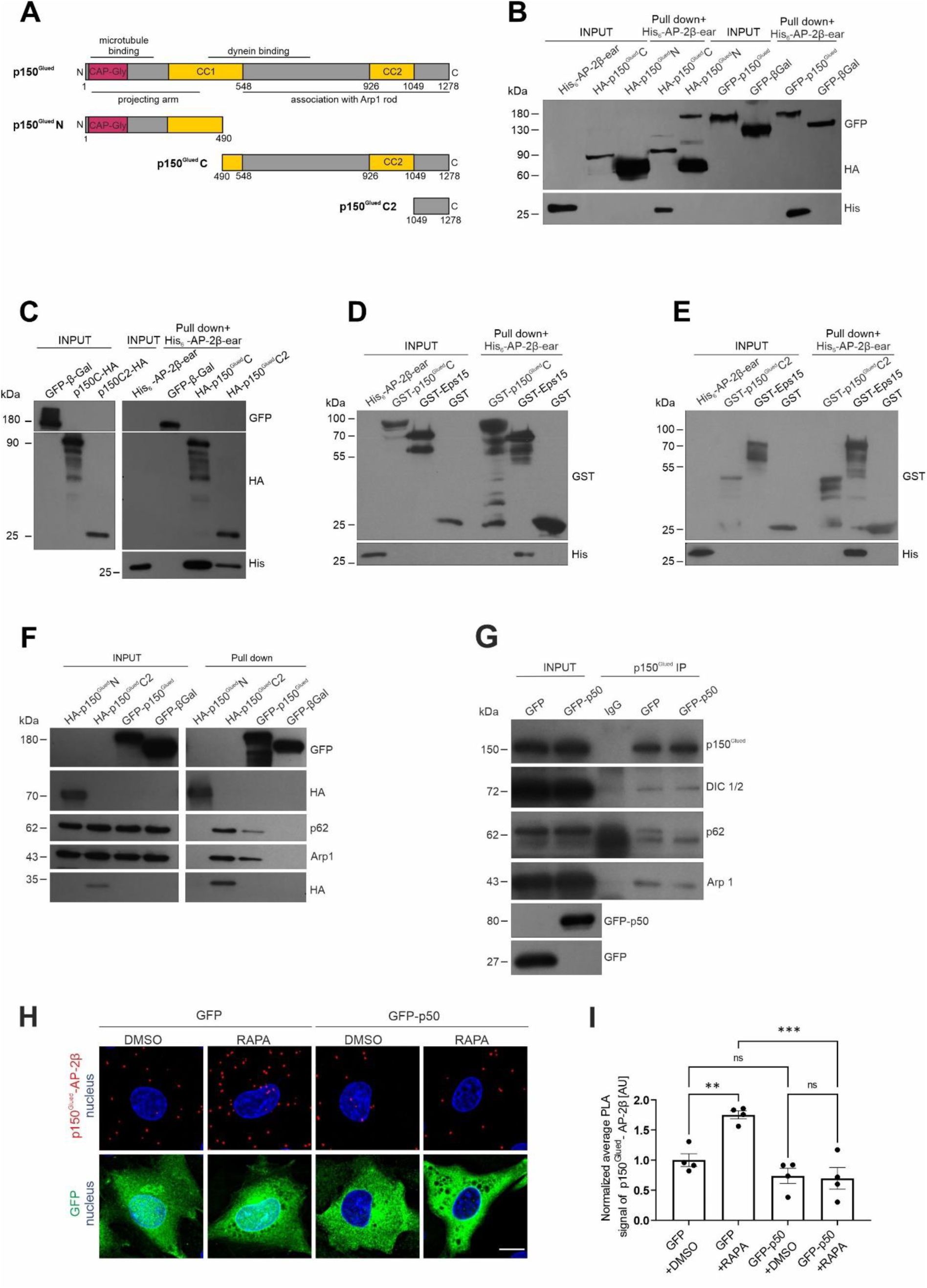
AP-2β interaction with p150^Glued^ is indirect and requires intact dynactin-dynein complex. (**A**) Diagram of the full p150^Glued^ and N, C, and C2 fragments that were used in the study. (**B**) Western blot analysis of *E. coli*-produced His-AP-2β-ear binding to *in vivo* biotinylated AviHA-tagged p150^Glued^ N or C fragments or AviGFP-tagged p150^Glued^ or AviGFP-tagged β-galactosidase that was pulled down from HEK293T cells using Avi-tag pull down. Input, 10% of lysate added to the assay. Shown is a representative example from *N* = 3 independent experiments. (**C**) Western blot analysis of *E. coli*-produced His-AP-2β-ear binding to *in vivo* biotinylated AviHA-tagged p150^Glued^ C or C2 fragments or AviGFP-tagged p150^Glued^ or AviGFP-tagged β-galactosidase that was pulled down from HEK293T cells using Avi-tag pull down. Input, 10% of lysate added to the assay. Shown is a representative example from *N* = 2 independent experiments. (**D, E**) Western blot analysis of pull down of *E. coli*-purified recombinant GST-p150^Glued^ C, GST-p150^Glued^ C2, GST-Eps15, or GST (negative control) with recombinant His-AP-2β ear. Shown is a representative example from *N* = 2 independent experiments. (**F**) Western blot analysis of endogenous p62 and Arp1 binding to *in vivo* biotinylated AviHA-tagged p150^Glued^ N or C2 fragments or full AviGFP-tagged p150^Glued^ or AviGFP-tagged β-galactosidase that was pulled down from HEK293T cells using Avi-tag pull down. Shown is a representative example from *N* = 3 independent experiments. (**G**) Western blot analysis of the co-immunoprecipitation of endogenous p150^Glued^ with other subunits of dynactin-dynein complex (DIC1/2, p62, and Arp1) from HEK293T cells that were transfected with pEGFPC1 or pEGFPC1-p50 plasmids. Shown is a representative example from *N* = 3 independent experiments. (**H**) Representative images of Rat2 fibroblasts that were transfected with pEGFPC1 or pEGFPC1-p50 (green) and treated for 2 h with 0.1% DMSO or 100 nM rapamycin (RAPA) with p150^Glued^–AP-2β PLA signals (red) and DAPI-stained nuclei (blue). Scale bar = 10 μm. (**I**) Quantification of the number of p150^Glued^–AP-2β PLA puncta in cells that were treated as in H. The data are expressed as the mean number of PLA puncta per cell, normalized to the control variant (GFP + DMSO) ± SEM. *N* = 4 independent experiments. *n* = 96 cells (GFP + DMSO), 81 cells (GFP + RAPA), 91 cells (GFP-p50 + DMSO), 92 cells (GFP-p50 + RAPA). *\*\*p* < 0.01, *\*\*\*p* < 0.001, *ns*, nonsignificant (two-way ANOVA followed by Tukey’s multiple-comparison *post hoc* test).

### CLIP-170 is needed for p150^Glued^–AP-2β interaction

p150^Glued^ is a +TIP. Therefore, we tested whether the p150^Glued^–AP-2β interaction requires its ability to target dynamic microtubules. p150^Glued^ microtubule plus-end targeting requires the presence of CLIP-170, which is also an mTOR substrate [17, 18, 58]. Thus, we tested whether CLIP-170 knockdown impacts the p150^Glued^–AP-2β interaction. We knocked down CLIP-170 in Rat2 and HEK293T cells using rat- and human-specific siRNAs, respectively, and performed the PLA and IP 72 h later. Both siRNAs against CLIP-170 effectively reduced CLIP-170 levels compared with cells that were transfected with control siRNAs (**Fig. 3A-D)**. The loss of CLIP-170 also potently prevented the rapamycin-induced p150^Glued^–AP-2β interaction (**Fig. 3A, E, F**). Notably, CLIP-170 knockdown did not affect the overall distribution of AP-2β in either control or rapamycin-treated cells (**Fig. S5A)**.

**Fig. 3.**
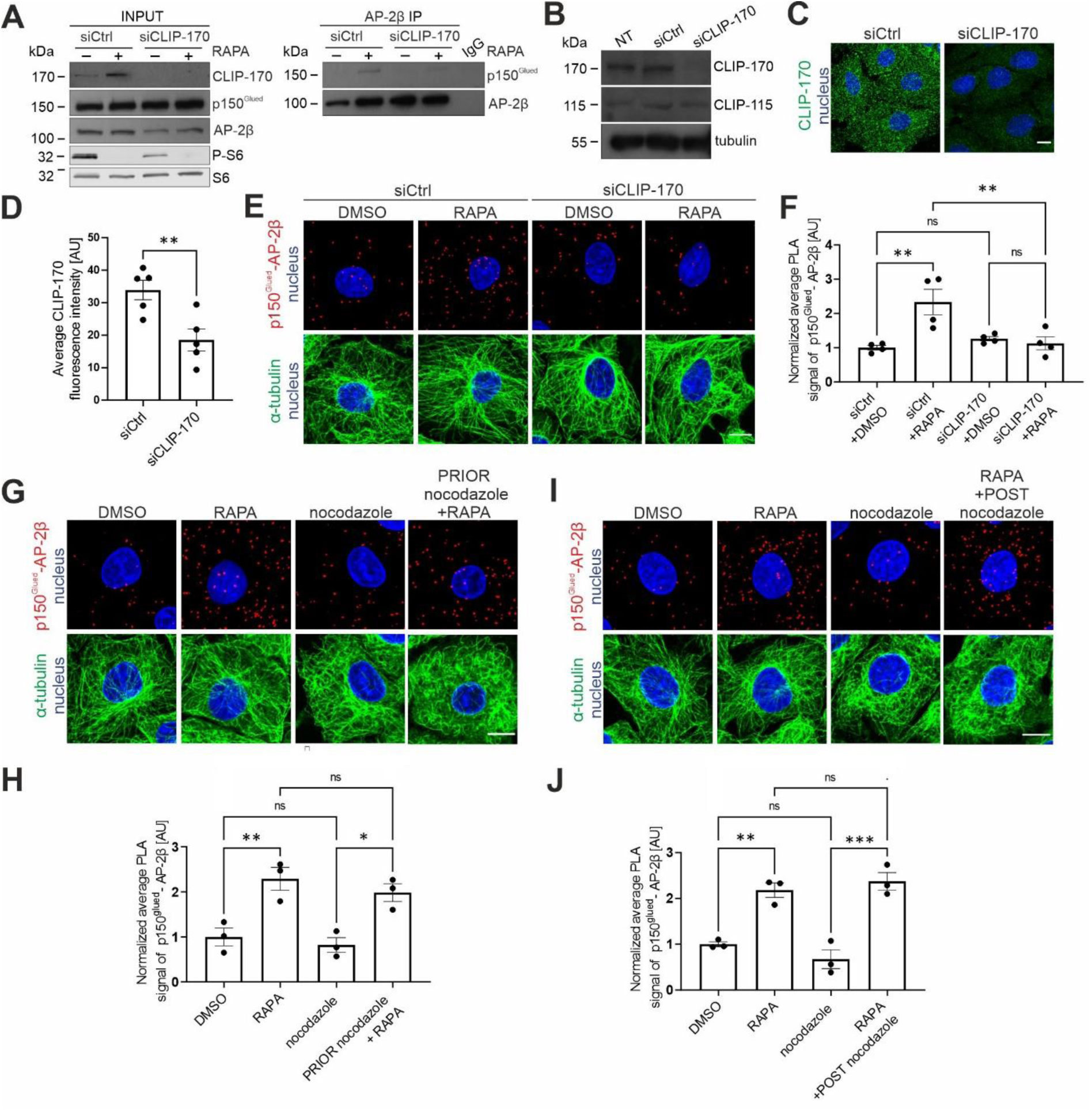
CLIP-170 but not microtubule dynamics is needed for p150^Glued^–AP-2β interaction. (**A**) Western blot analysis of endogenous CLIP-170, p150^Glued^, AP-2β, S6, and P-S6 (Ser235/236) levels and co-immunoprecipitation of endogenous AP-2β with p150^Glued^ in HEK293T cells that were transfected with control siRNA (siCtrl) or siRNA against human CLIP-170 (siCLIP-170) and treated for 2 h with 0.1% DMSO or 100 nM rapamycin (RAPA). Input, 10% of lysate added for immunoprecipitation. Shown is a representative example from *N* = 3 independent experiments. (**B**) Western blot analysis of endogenous CLIP-170 and CLIP-115 levels in non-transfected (NT) Rat2 cells or Rat2 cells that were transfected with siCtrl or rat siCLIP-170. (**C**) Representative images of Rat2 cells that were transfected with siCtrl or rat siCLIP-170, with immunofluorescently labeled endogenous CLIP-170 (green) and nucleus stained with Hoechst 33258 (blue). Scale bar = 10 µm. (**D**) Quantitative analysis of CLIP-170 immunofluorescence in Rat2 cells that were treated as in C. The results are presented as the average intensity of CLIP-170 immunofluorescence in the cell ± SEM. *N* = 5 independent experiments. \*\**p* < 0.01 (Student’s *t*-test). **(E)** Representative images of Rat2 fibroblasts that were transfected with siCtrl or rat siCLIP-170, and treated for 2 h with 0.1% DMSO or 100 nM rapamycin (RAPA), with p150^Glued^–AP-2β PLA signals (red), immunofluorescently labeled tubulin (green), and DAPI-stained nuclei (blue). Scale bar = 10 μm. (**F**) Quantification of the number of p150^Glued^–AP-2β PLA puncta in cells that were treated as in E. The data are expressed as the mean number of PLA puncta per cell, normalized to the control variant (siCtrl + DMSO) ± SEM. *N* = 4 independent experiments. *n* = 149 cells (siCtrl + DMSO), 130 cells (siCtrl + RAPA), 161 cells (siCLIP-170 + DMSO), 142 cells (siCLIP-170 + RAPA). *\*\*p* < 0.01, *ns*, nonsignificant (two-way ANOVA followed by Tukey’s multiple-comparison *post hoc* test). (**G**) Representative images of Rat2 fibroblasts with p150^Glued^–AP-2β PLA signals (red), immunofluorescently labeled tubulin (green), and DAPI-stained nuclei (blue). Cells were treated for 2 h with 0.1% DMSO or 100 nM rapamycin (RAPA) or treated for 2 h 15 min with 100 nM nocodazole alone or in combination with 100 nM rapamycin that was added 15 min after nocodazole (PRIOR nocodazole + RAPA). Scale bar = 10 μm. (**H**) Quantification of the number of p150^Glued^–AP-2β PLA puncta in cells that were treated as in G. The data are expressed as the mean number of PLA puncta per cell, normalized to the control variant (DMSO) ± SEM. *N* = 3 independent experiments. *n* = 126 cells (DMSO), 125 cells (RAPA), 133 cells (nocodazole), 139 cells (PRIOR nocodazole + RAPA). *\*p* < 0.05, *\*\*p* < 0.01, *ns*, nonsignificant (one-way ANOVA followed by Bonferroni multiple-comparison *post hoc* test). (**I**) Representative images of Rat2 fibroblasts with p150^Glued^–AP-2β PLA signals (red), immunofluorescently labeled tubulin (green), and DAPI-stained nuclei (blue). Cells were treated for 2 h with 0.1% DMSO or 100 nM rapamycin (RAPA) or treated for 1 h with 100 nM nocodazole alone or added in the middle of 2 h of 100 nM rapamycin incubation (RAPA + POST nocodazole). Scale bar = 10 μm. (**J**) Quantification of the number of p150^Glued^–AP-2β PLA puncta in cells that were treated as in I. The data are expressed as the mean number of PLA puncta per cell, normalized to the control variant (DMSO) ± SEM. *N* = 3 independent experiments. *n* = 128 cells (DMSO), 118 cells (RAPA), 136 cells (nocodazole), 121 cells (RAPA + POST nocodazole). *\*\*p* < 0.01, *\*\*\*p* < 0.001, *ns*, nonsignificant (one-way ANOVA followed by Bonferroni multiple-comparison *post hoc* test).

CLIP-170 knockdown may result, in addition to p150^Glued^ displacement, in substantial changes in microtubule dynamics, especially in cells that do not express CLIP-115 [17]. Although Rat2 cells express both CLIPs and our siRNAs did not target CLIP-115 (**Fig. 3B**), we directly tested effects of CLIP-170 knockdown on microtubule dynamics and the role of microtubule dynamics in the AP-2–dynactin interaction. Indeed, CLIP-170 knockdown had no effect on EB3-GFP mobility that highlights dynamic plus ends of microtubules (**Fig. S5B-F, Movies 5-6**). Next, we performed a p150^Glued^–AP-2β PLA in Rat2 cells that were treated with 100 nM nocodazole that was added either 15 min before or 1 h after rapamycin treatment. At such low concentrations, nocodazole blocks plus-end microtubule dynamics instead of depolymerizing microtubules [59]. Indeed, 1 h of the nocodazole treatment of Rat2 cells resulted in the loss of EB3-GFP and CLIP-170 comets, confirming the inhibition of microtubule dynamics (**Fig. S6**, **Movies 7-10**). Such treatment did not affect the p150^Glued^–AP-2β PLA signal under basal conditions or in response to rapamycin treatment (**Fig. 3G-J**). In contrast to nocodazole, rapamycin treatment did not affect EB3-GFP microtubule plus-end tracking behavior (**Fig. S7A-D**, **Movie 11, 12**). Notably, rapamycin did not change the plus-end tracking behavior of CLIP-170–GFP that was overexpressed in Rat2 cells (**Fig. S7E-H**, **Movie 13, 14**). This observation is consistent with our previous data on the lack of effect of rapamycin on endogenous CLIP-170 microtubule binding in neurons and HeLa cells [58].

Overall, the data suggest that microtubule dynamics, at least in the short term, is not needed for the p150^Glued^–AP-2β interaction. Low-dose nocodazole treatment should also result in p150^Glued^ displacement from dynamic microtubule plus ends, like CLIP-170, but we did not observe any impact of nocodazole on p150^Glued^–AP-2β complex formation. Thus, p150^Glued^ displacement from microtubules unlikely explains the effects of CLIP-170 knockdown on the p150^Glued^–AP-2β PLA interaction. We confirmed this hypothesis using a dominant-negative CLIP-170 mutant that lacked the N-terminal part of the protein and was previously shown to severely affect microtubule dynamics and displace p150^Glued^ from microtubule plus ends [60]. The 48 h overexpression of this protein in Rat2 cells did not prevent p150^Glued^–AP-2β interaction measured with PLA in the rapamycin-treated cells (**Fig. S8**). In summary, CLIP-170 is needed for the p150^Glued^–AP-2β interaction, but two of its canonical functions (i.e., the regulation of microtubule dynamics and targeting p150^Glued^ to microtubule plus ends) do not appear to be directly or immediately involved.

### mTORC1-dependent autophagy triggers p150^Glued^–AP-2β interaction

Like mTOR, AP-2 was postulated to be essential for autophagy initiation [25]. Thus, we investigated whether mTORC1-controlled autophagy under the conditions that were used in the present study is induced and required for the rapamycin-driven p150^Glued^–AP-2β interaction. Treatment with rapamycin decreased P-S6 level, demonstrating mTORC1 inhibition and resulting in an increase in beclin-1 phosphorylation at Ser30, which is an Ulk-1 target and considered an early marker of autophagy (**Fig. S9A**). Furthermore, after 2 h, rapamycin also increased the ratio of the lipidated form of LC3 (LC3B II) to non-lipidated LC3B I, which is routinely used to assess autophagy (**Fig. S9A, B**). Furthermore, rapamycin also increased the formation of large LC3B foci, further confirming autophagy induction (**Fig. S9C**). To verify whether autophagy induction is required for the mTORC1 inhibition-driven AP-2–dynactin interaction, we investigated whether pretreatment with the autophagy initiation inhibitor SBI-0206965 (25 µM; 30 min before rapamycin administration) counteracts the effects of rapamycin. As expected, pretreatment with SBI-0206965 was sufficient to decrease rapamycin-induced beclin-1 phosphorylation at Ser30, LC3 lipidation, and the formation of endogenous large LC3 foci (**Fig. S9A-C**). Notably, blocking autophagy initiation completely abolished the rapamycin-induced increase in the AP-2β-p150^Glued^ interaction, measured by IP and the PLA in HEK293T and Rat2 cells, respectively (**Fig. 4A-C**). To further confirm that early steps of autophagy are required for the rapamycin-induced interaction of AP-2β and p150^Glued^, we tested whether the knockdown of Atg5, a key protein for this process, exerts an identical effect as SBI-0206965. Rat2 cells were transfected with siAtg5 and siCtrl for 72 h. Western blot showed that siAtg5 effectively reduced Atg5 levels in transfected cells compared with the control and simultaneously inhibited the autophagy process, indicated by a decrease in the LC3B II/LC3B I ratio (**Fig. S9D-F**). The silencing of Atg5 also effectively counteracted the rapamycin-induced increase in the p150^Glued^–AP-2β PLA signal (**Fig. 4D, E**). Thus, the induction of autophagy is needed for p150^Glued^–AP-2β protein complex formation upon mTORC1 inhibition.

**Fig. 4.**
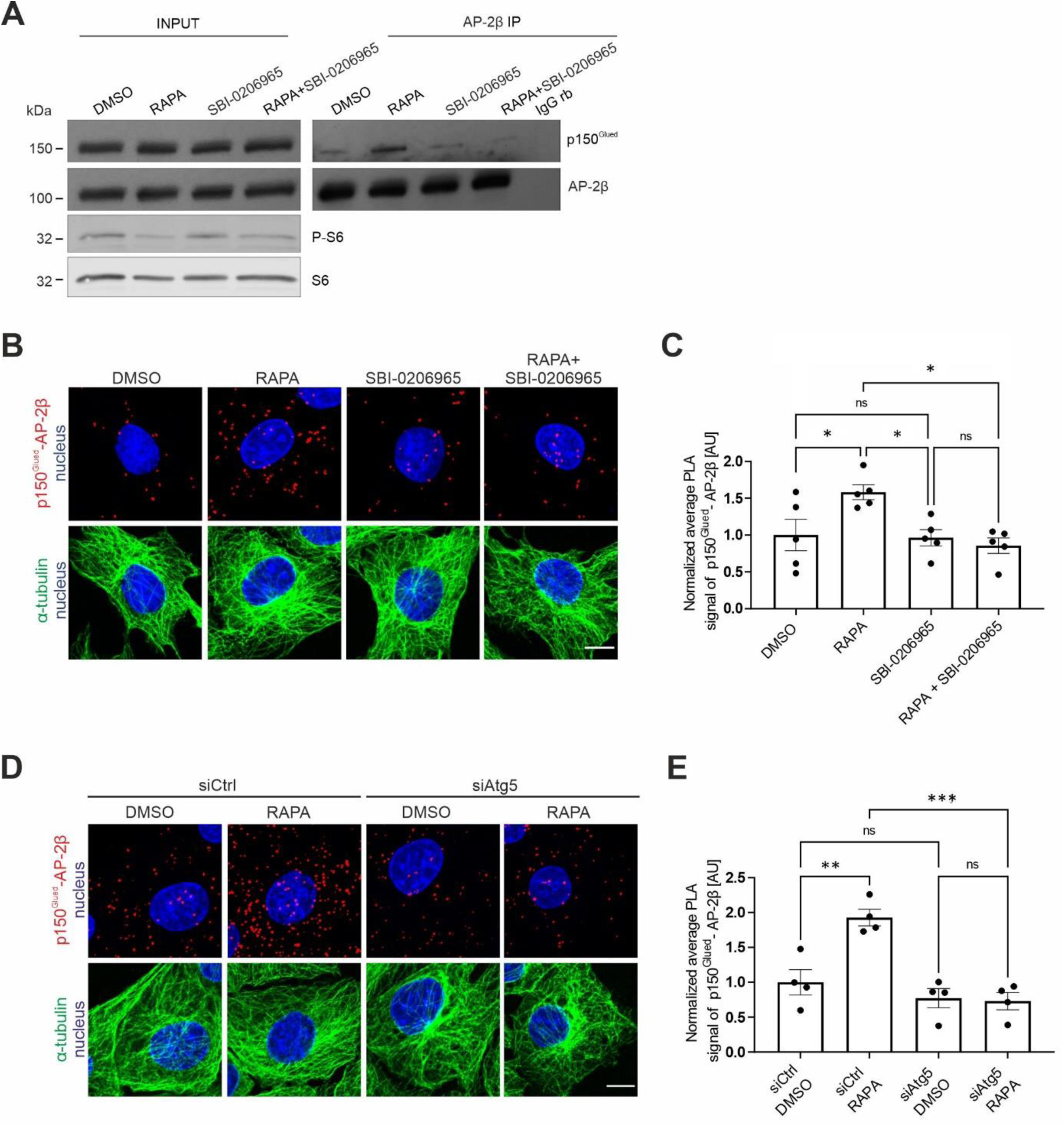
Autophagy induction upon mTORC1 inhibition is needed for p150^Glued^–AP-2β interaction. (**A**) Western blot analysis of endogenous p150^Glued^, AP-2β, S6, and P-S6 (Ser235/236) levels and co-immunoprecipitation of endogenous AP-2β with p150^Glued^ in HEK293T cells that were treated with 0.1% DMSO for 2 h, 100 nM rapamycin (RAPA) for 2 h, 25 μM SBI-0206965 for 2 h 30 min, or 25 μM SBI-0206965 for 30 min and 100 nM rapamycin for 2 h (RAPA + SBI-0206965). Input, 10% of lysate used for immunoprecipitation. Shown is a representative example from *N =* 2 independent experiments. (**B**) Representative images of Rat2 fibroblasts with p150^Glued^–AP-2β PLA signals (red), immunofluorescently labeled tubulin (green), and DAPI-stained nuclei (blue). Cells were treated with 0.1% DMSO for 2 h, 100 nM rapamycin (RAPA) for 2 h, 25 μM SBI-0206965 for 2 h 30 min, or 25 μM SBI-0206965 for 30 min and 100 nM rapamycin for 2 h (RAPA + SBI-0206965). Scale bar = 10 μm. (**C**) Quantification of the number of p150^Glued^-AP-2β PLA puncta in cells that were treated as in B. The data are expressed as the mean number of PLA puncta per cell, normalized to the control variant (DMSO) ± SEM. *N* = 5 independent experiments. *n* = 184 cells (DMSO), 189 cells (RAPA), 169 cells (SBI-0206965), 174 cells (RAPA + SBI-0206965). *\*p* < 0.05, *ns*, nonsignificant (one-way ANOVA followed by Bonferroni multiple-comparison *post hoc* test). (**D**) Representative images of Rat2 fibroblasts that were transfected with siCtrl or rat siAtg5 for 72 h and then treated for 2 h with 0.1% DMSO or 100 nM rapamycin (RAPA), with PLA p150^Glued^–AP-2β signals (red), immunofluorescently labeled tubulin (green), and DAPI-stained nuclei (blue). Scale bar = 10 μm. (**E**) Quantification of the number of p150^Glued^–AP-2β PLA puncta in cells that were treated as in D. The data are expressed as the mean number of PLA puncta per cell, normalized to the control variant (siCtrl + DMSO) ± SEM. *N* = 4 independent experiments. *n* = 199 cells (siCtrl + DMSO), 178 cells (siCtrl + RAPA), 195 cells (siAtg5 + DMSO), 211 cells (siAtg5 + RAPA). \*\*\**p* < 0.001, \*\**p* < 0.01, *ns*, nonsignificant (two-way ANOVA followed by Tukey’s multiple-comparison post hoc test).

### Autophagy initiation is sufficient for p150^Glued^–AP-2β interaction

Our results above show that the initiation of autophagy is essential for effects of mTORC1 inhibition on p150^Glued^–AP-2β complex formation. However, a key issue is whether autophagy initiation, even when mTORC1 is active, is sufficient to induce a similar effect. To investigate this possibility, we treated Rat2 cells with L-690330, an inhibitor of inositol monophosphatase and mTOR-independent activator of autophagy [76], and performed an p150^Glued^–AP-2β PLA. After 3 h of L-690330 (100 µM) treatment, autophagy levels increased, indicated by the LC3B II/LC3B I ratio and formation of large LC3B foci, whereas the level of phosphorylated S6 at Ser235/236 did not decrease as expected (**Fig. 5A-C**). This treatment also increased the p150^Glued^–AP-2β PLA signal similarly to rapamycin treatment (**Fig. 5D, E**). Altogether, our results show that autophagy induction alone is sufficient to induce AP-2– dynactin complex formation. Additionally, as in the case of rapamycin treatment, CLIP-170 knockdown blocked the L-690330-induced increase in the p150^Glued^–AP-2β interaction (**Fig. 5F-H**). This observation suggests potential novel autophagy-related activities of CLIP-170. Thus, we tested whether CLIP-170 knockdown affects autophagy that is induced by rapamycin or L-690330. The loss of CLIP-170 prevented the increase in the LC3B II/LC3B I ratio and LC3 foci formation that were caused by both autophagy inducers (**Fig. 5I-K**). These findings suggest that the dynactin interaction with the AP-2 adaptor complex requires autophagy initiation, which depends on the presence of CLIP-170.

**Fig. 5.**
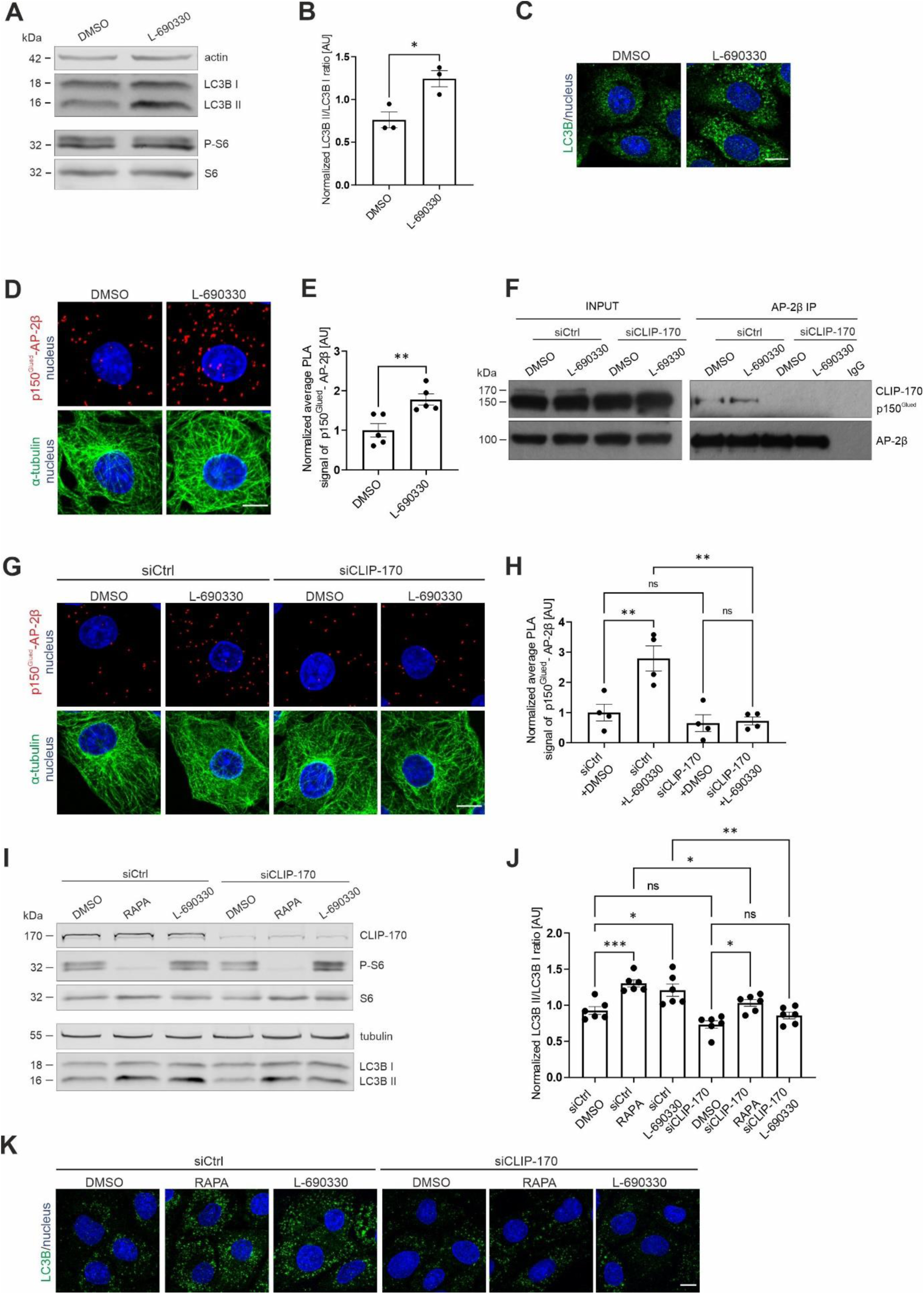
p150^Glued^–AP-2β interaction is induced by autophagy even when mTOR activity is preserved. (**A**) Western blot analysis of endogenous actin, LC3B I, LC3B II, P-S6 (Ser235/236), and S6 levels in Rat2 fibroblasts that were treated for 3 h with 0.1% DMSO or 100 μM L-690330. Shown is a representative example from *N =* 3 independent experiments. (**B**) Densitometry analysis of normalized LC3B II/LCB I ratio in Rat2 cells that were treated as in A. The data are presented as mean of the normalized ratio of LC3B II to LC3B I levels ± SEM. *N* = 3 independent experiments. *\*p ≤* 0.05 (one-tailed Mann-Whitney test). (**C**) Representative images of Rat2 cells that were treated for 3 h with 0.1% DMSO or 100 μM L-690330 with immunofluorescently labeled endogenous LC3B (green) and nuclei stained with Hoechst 33258 (blue). Scale bar = 10 µm. (**D**) Representative images of Rat2 fibroblasts that were treated for 3 h with 0.1% DMSO or 100 μM L-690330 with p150^Glued^–AP-2β PLA signals (red), immunofluorescently labeled tubulin (green), and DAPI-stained nuclei (blue). Scale bar = 10 μm. (**E**) Quantification of the number of p150^Glued^–AP-2β PLA puncta in cells that were treated as in D. The data are expressed as the mean number of PLA puncta per cell, normalized to the control variant (DMSO) ± SEM. *N* = 5 independent experiments. *n* = 200 cells (DMSO), 181 cells (L-690330). *\*\*p* < 0.01 (Student’s *t*-test). (**F**) Western blot analysis of endogenous CLIP-170, p150^Glued^, and AP-2β levels and co-immunoprecipitation of endogenous AP-2β with p150^Glued^ in HEK293T cells that were transfected with control siRNA (siCtrl) or siRNA against human CLIP-170 (siCLIP-170) and treated for 3 h with 0.1% DMSO or 100 μM L-69330. Input, 10% of lysate used for immunoprecipitation. Shown is a representative example from *N =* 2 independent experiments. (**G**) Representative images of Rat2 fibroblasts that were transfected with siCtrl or rat siCLIP-170 and treated for 3 h with 0.1% DMSO or 100 μM L-690330 with PLA p150^Glued^–AP-2β signals (red), immunofluorescently labeled tubulin (green), and DAPI-stained nuclei (blue). Scale bar = 10 μm. (**H**) Quantification of the number of p150^Glued^–AP-2β PLA puncta in cells that were treated as in G. The data are expressed as the mean number of PLA puncta per cell, normalized to the control variant (siCtrl +DMSO) ± SEM. *N* = 4 independent experiments. *n* = 167 cells (siCtrl + DMSO), 163 cells (siCtrl + L-690330), 146 cells (siCLIP-170 + DMSO), 164 cells (siCLIP-170 + L-690330). *\*\*p* < 0.01, *ns*, nonsignificant (two-way ANOVA followed by Tukey’s multiple-comparison *post hoc* test). (**I**) Western blot analysis of endogenous CLIP-170, P-S6 (Ser235/236), S6, tubulin, LC3B I, and LC3B II levels in Rat2 fibroblasts that were transfected with siCtrl or rat siCLIP-170 and treated for 2 h with 0.1% DMSO or 100 nM rapamycin (RAPA) or treated for 3 h with 100 μM L-690330. Shown is a representative example from *N* = 6 independent experiments. (**J**) Densitometry analysis of normalized LC3B II/LCB I ratio in Rat2 cells that were treated as in I. The data are presented as mean of the normalized ratio of LC3B II to LC3B I levels ± SEM. *N* = 6 independent experiments. *\*p* < 0.05, *\*\*p* < 0.01, *\*\*\*p* < 0.001, *ns*, nonsignificant (two-way ANOVA followed by Tukey’s multiple-comparison *post hoc* test). (**K**) Representative images of Rat2 cells that were transfected with siCtrl or rat siCLIP-170 and treated for 2 h with 0.1% DMSO or 100 nM rapamycin (RAPA) or treated for 3 h with 100 μM L-690330 with immunofluorescently labeled endogenous LC3B (green) and nuclei stained with Hoechst 33258 (blue). Scale bar = 10 µm.

### ABMA and chloroquine prevent rapamycin-induced p150^Glued^–AP-2β interaction

Based on the data that were obtained, the initiation of autophagy appears to play a key role in the p150^Glued^–AP-2β-protein interaction, but unknown is whether undisturbed autophagic flux is also required. Therefore, we treated cells with rapamycin in the presence of chloroquine (CQ) and ABMA, two compounds that affect this process via different mechanisms [77, 78]. ABMA stimulates the formation of amphisomes, which however are unable to fuse with lysosomes to finish the autophagy [79]. Chloroquine appears to have pleiotropic effects that include direct blockade of the fusion of autophagosomes with lysosomes by preventing the recruitment of SNAP29 to the fusion site, slowing the acidification of lysosomes or disturbing endosomal flow [78, 80]. Rat2 cells were treated with rapamycin in the presence of ABMA (60 µM) or CQ (50 µM) for 2 h. Cells that were treated with DMSO, ABMA, or CQ alone served as controls. Although fluorescence analysis did not show a profound enhancement of LC3B protein cluster formation after treatment with ABMA (**Fig. S10A**), Western blot confirmed that it increased the LC3B II/LC3B I ratio as expected (**Fig. S10B, C**) [77]. Treatment with CQ also resulted in autophagic flux inhibition as described previously (**Fig. S10D-F)** [78]. Both ABMA and CQ blocked the increase in the p150^Glued^–AP-2β PLA signal that was caused by rapamycin (**Fig. 6A-D**). ABMA and CQ prevent the fusion of autolysosome-preceding compartments with lysosomes but may also have additional effects on the endomembrane system. Therefore, we tested whether the knockdown of SNAP29, which plays a key role in autophagosome-lysosome fusion [81], affects the rapamycin-induced p150^Glued^–AP-2β interaction. Rat2 cells were transfected with control siRNA or siRNA against Snap29 (siSnap29) for 72 h. The qRT-PCR analysis of RNA that were isolated from siRNA-transfected cells showed a significant decrease in Snap29 mRNA levels compared with controls (**Fig. S10G**). Additionally, immunofluorescent staining and Western blot analysis showed an increase in p62/SQSTM1 protein (**Fig. S10H-J**), which is expected to accumulate in the cell in the absence of Snap29 and with autophagosome-lysosome fusion inhibition. However, Snap29 knockdown did not prevent the enhancement of the p150^Glued^–AP-2β PLA signal by rapamycin (**Fig. 6E, F**). Thus, we concluded that both ABMA and CQ effectively prevented the rapamycin-induced p150^Glued^–AP-2β interaction, but SNAP29-dependent fusion unexpectedly did not appear to be required.

**Fig. 6.**
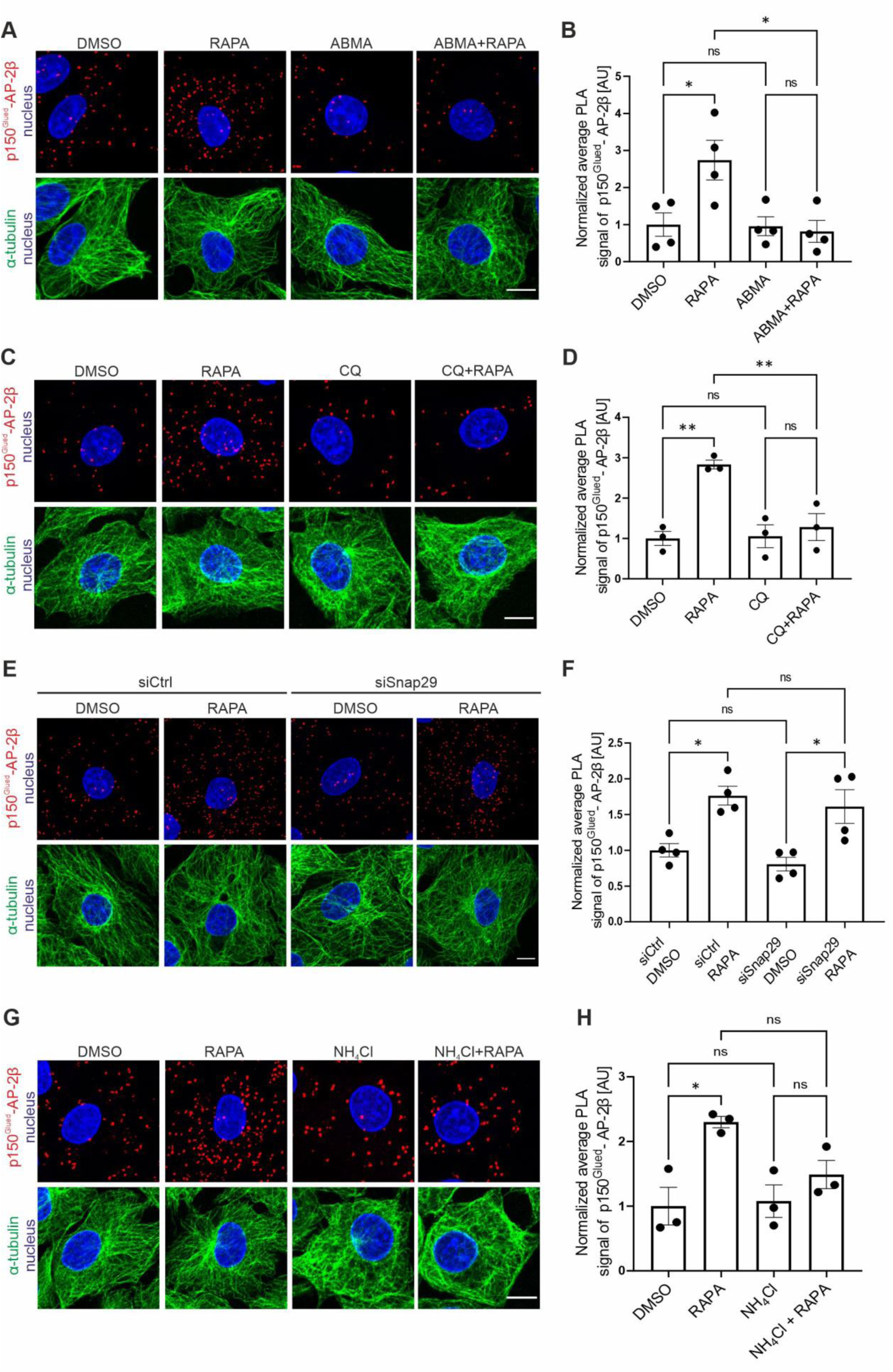
ABMA and chloroquine prevent rapamycin-induced p150^Glued^–AP-2β interaction. (**A**) Representative images of Rat2 fibroblasts that were treated for 2 h with 0.1% DMSO, 100 nM rapamycin (RAPA), 60 μM ABMA (ABMA), or 60 μM ABMA and 100 nM rapamycin (ABMA + RAPA) with p150^Glued^–AP-2β PLA signals (red), immunofluorescently labeled tubulin (green), and DAPI-stained nuclei (blue Scale bar = 10 μm. (**B**) Quantification of the number of p150^Glued^–AP-2β PLA puncta in cells that were treated as in A. The data are expressed as the mean number of PLA puncta per cell, normalized to the control variant (DMSO) ± SEM. *N* = 4 independent experiments. *n* = 140 cells (DMSO), 144 cells (RAPA), 148 cells (ABMA), 134 cells (ABMA + RAPA). *\*p* < 0.05, *ns* – nonsignificant (one-way ANOVA followed by Bonferroni multiple-comparison *post hoc* test). (**C**) Representative images of Rat2 fibroblasts that were treated for 2 h with 0.1% DMSO, 100 nM rapamycin (RAPA), 50 μM chloroquine (CQ), or 50 μM chloroquine and 100 nM rapamycin for 2 h (CQ + RAPA) with PLA p150^Glued^–AP-2β signals (red), immunofluorescently labeled tubulin (green), and DAPI-stained nuclei (blue). Scale bar = 10 μm. (**D**) Quantification of the number of p150^Glued^–AP-2β PLA puncta in cells that were treated as in C. The data are expressed as the mean number of PLA puncta per cell, normalized to the control variant (DMSO) ± SEM. *N* = 3 independent experiments. *n* = 142 cells (DMSO), 143 cells (RAPA), 135 cells (CQ), 154 cells (CQ + RAPA). *\*\*p* < 0.01, *ns* – nonsignificant (one-way ANOVA followed by Bonferroni multiple-comparison *post hoc* test). (**E**) Representative images of Rat2 fibroblasts that were transfected with siCtrl or rat siSnap29 for 72 h and then treated for 2 h with 0.1% DMSO or 100 nM rapamycin (RAPA), with p150^Glued^–AP-2β PLA signals (red), immunofluorescently labeled tubulin (green), and DAPI-stained nuclei (blue). Scale bar = 10 μm. (**F**) Quantification of the number of p150^Glued^–AP-2β PLA puncta in cells that were treated as in E. The data are expressed as the mean number of PLA puncta per cell, normalized to the control variant (siCtrl + DMSO) ± SEM. *N* = 4 independent experiments. *n* = 172 cells (siCtrl + DMSO), 161 cells (siCtrl + RAPA), 172 cells (siSnap29 + DMSO), 179 cells (siSnap + RAPA). \**p* < 0.05, *ns* – nonsignificant (two-way ANOVA followed by Tukey’s multiple-comparison post hoc test). (**G**) Representative images of Rat2 fibroblasts that were treated with 0.1% DMSO for 2 h, 100 nM rapamycin (RAPA) for 2 h, 20 mM NH_4_Cl (NH_4_Cl) for 3 h, or pretreated with 20 mM NH_4_Cl for 1 h and treated with 100 nM rapamycin for 2 h (NH_4_Cl + RAPA) with p150^Glued^-AP-2β PLA signals (red), immunofluorescently labelled tubulin (green), and DAPI-stained nuclei (blue). Scale bar = 10 μm. (**H**) Quantification of the number of p150^Glued^–AP-2β PLA puncta in cells that were treated as in G. The data are expressed as the mean number of PLA puncta per cell, normalized to the control variant (DMSO) ± SEM. *N* = 3 independent experiments, *n* = 122 cells (DMSO), 109 cells (RAPA), 125 cells (NH_4_Cl), 114 cells (NH_4_Cl + RAPA). *\*p* < 0.05, *ns* – nonsignificant (one-way ANOVA followed by Bonferroni multiple-comparison *post hoc* test).

Because CQ can alkalize the lysosome environment [80] and because ABMA was not tested for it in Rat2 cells, we used lysotracker staining to investigate effects of rapamycin, CQ, ABMA, and their combination on lysosomal acidification in our experimental model. As a control, we treated cells for 2 h with the vATPase inhibitor Baf A1. Two hours of treatment with rapamycin significantly increased the intensity of lysotracker staining and the number of positive lysotracker structures (**Fig. S11**). In contrast, both ABMA and CQ alone and in the presence of rapamycin significantly decreased both parameters (**Fig. S11**). However, effects of CQ and ABMA were different. Chloroquine had a stronger effect that was comparable to treatment with Baf A1, while influence of ABMA was much milder (**Fig. S11**). Thus, we concluded that although autophagosome fusion with the endolysosomal pathway did not affect the p150^Glued^–AP-2β interaction, ABMA and CQ might affect it, leading to improper lysosome acidification or disruption of the endolysosomal pathway as previously shown [77, 78]. To test the hypothesis that the correct pH in the cell is necessary for the rapamycin-induced interaction of p150^Glued^ and AP-2β, we repeated the experiment, this time treating the cells with 20 mM NH_4_Cl to alkalize the cells before administering rapamycin and running PLA (**Fig. 6G, H**). As in the previous experiments, rapamycin caused an increase in p150^Glued^–AP-2β PLA. At the same time, administration of NH_4_Cl under basal conditions had no significant effect on PLA signal. However, incubation with NH_4_Cl counteracted the increase in p150^Glued^–AP-2β PLA in response to rapamycin administration, supporting the hypothesis that appropriate acidification of the endolysosomal pathway is critical for the interaction of p150^Glued^ and AP-2β.

### Rapamycin enhances the lysosomal p150^Glued^–AP-2β interaction and affects Lamp-1 mobility

Based on the above observations, we investigated where in Rat2 cells p150^Glued^ and AP-2β interact. We used the PLA that was adjusted for EM, revealing that the p150^Glued^–AP-2β PLA signal in rapamycin-treated Rat2 cells localized primarily to organelles that contained electron dense material that is characteristic for lysosomes (43% of cells with PLA-EM signal) and in double-membrane organelles, which we classified as autophagosomes (57% of cells with PLA-EM signal). In control, in DMSO-treated cells, the PLA-EM signal was also spotted but less frequently and rarely on lysosome-like structures (**Fig. 7A**; 14% *vs*. 43% of cells with PLA-EM signal). Because the PLA procedure did not allow the preservation of a high-quality ultrastructure, we additionally analyzed in rapamycin-treated Rat2 cells the co-occurrence of the p150^Glued^–AP-2β PLA signal with the LC3B or lysosomal marker Lamp-1, either endogenous or overexpressed as a GFP fusion, using AiryScan high-resolution light microscopy. For LC3B, some co-occurrence was detected but not frequently (**Fig. S12**). For Lamp-1, we noticed an apparent, although still partial, co-localization of the PLA signal with lysosomes (**Fig. 7B, C**). Thus, the combined observations from PLA-EM and Airyscan images indicated that lysosomes are the primary localization of the p150^Glued^–AP-2β interaction under rapamycin treatment, raising the issue of whether rapamycin affects the mobility of Lamp-1-positive compartments.

**Fig. 7.**
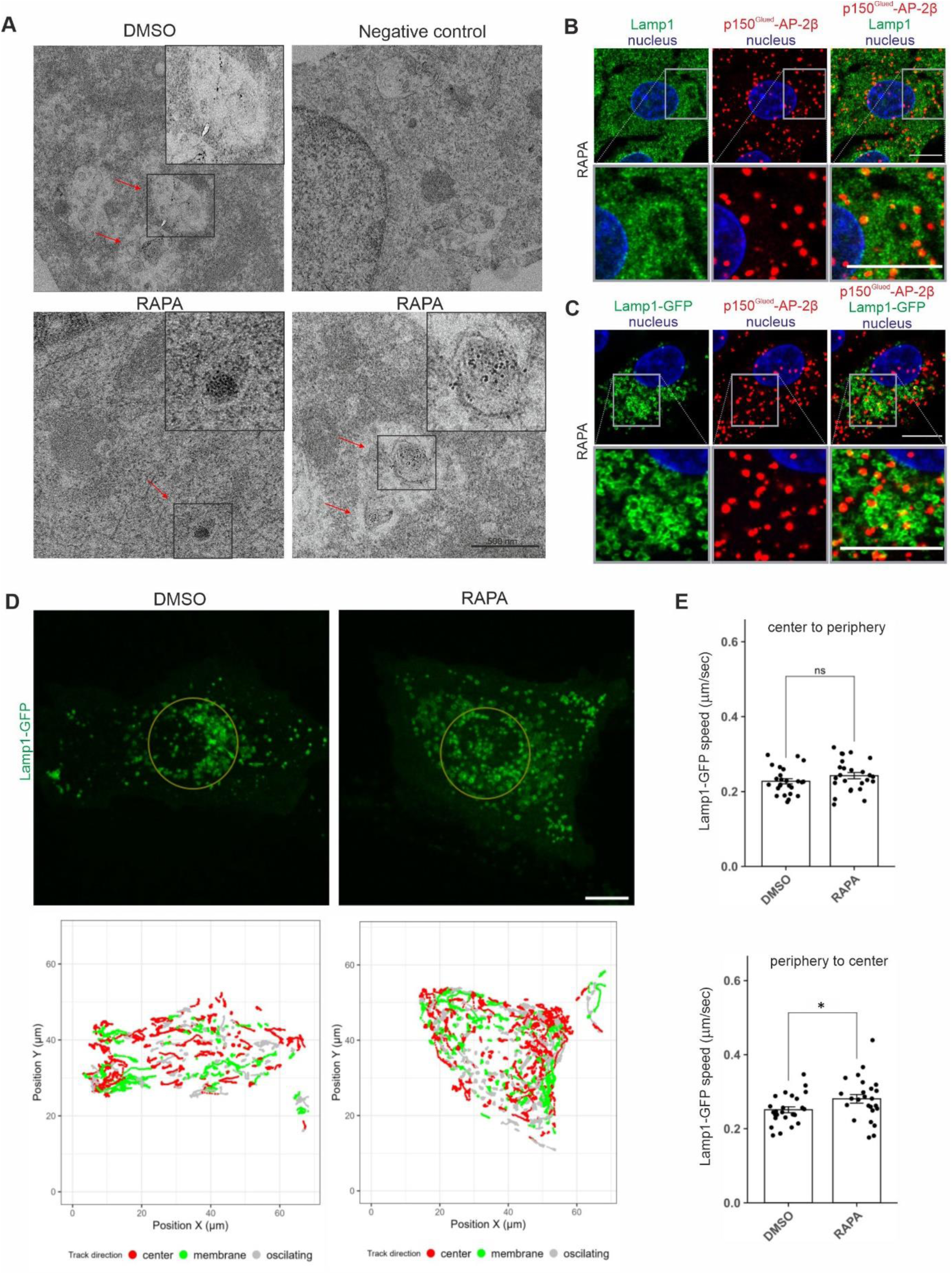
Rapamycin induces p150^Glued^-AP-2β interaction on lysosomes and regulates lysosomal mobility. (**A**) Representative electron microscopy images of Rat2 fibroblasts after 2 h treatment with 0.1% DMSO or 100 nM rapamycin (RAPA) and PLA analysis of p150^Glued^– AP-2β with and without (negative control) primary antibodies. Red arrows point to PLA signals that co-occurred with organelles that resembled lysosomes or autophagosomes. Black boxes indicate the regions shown in higher magnification. *n* = 30 cells per variant. *N* = 3 independent experiments. Scale bar = 500 nm. (**B**) Representative images of Rat2 cells treated with 100 nM rapamycin (RAPA) with p150^Glued^–AP-2β PLA signals (red) immunofluorescently labeled endogenous Lamp1 (green) and DAPI-stained nuclei (blue). Images were acquired using the AiryScan module. Scale bar = 10 μm. (Upper panel) Representative photograph of a single cell. (Lower panel) Close-up of a site with a high intensity of PLA-Lamp1 co-localization. (**C**) Representative images of Rat2 cells transfected with Lamp1-GFP (green) for 24 h and then treated with 100 nM rapamycin (RAPA) with p150^Glued^–AP-2β PLA signals (red), and DAPI-stained nuclei (blue). Images were acquired using the AiryScan module. Scale bar = 10 μm. (Top) Representative photograph of a single cell. (Bottom) Close-up of a site with a high intensity of PLA-Lamp1-GFP co-localization. (**D**) Representative images of cells that expressed Lamp1-GFP. The upper row shows the first frames from the time-lapse movies. The area inside the golden circle is considered the “center” compartment, and all movements outside this area are “peripheries.” The lower row shows trajectories (tracks) that were identified by the ImageJ “TrackMate” plugin that were longer than 6.8 mm (100 pixels). Trajectories were color-coded based on their directions, which were established using Pearson correlation coefficient (PCC) calculated by change in the distance from the cell center in time. If the distance was increasing with consecutive frames (PCC: 0.5 to 1) tracks were considered to move to the cell membrane. If distance was decreasing (PCC: -1 to -0.5), direction was described as moving to the center. Values in between were marked as oscillating. (**E**) Difference in speed of Lamp1-GFP vesicles’ movements between rapamycin-treated (RAPA) and control (DMSO) cells. The values are mean trajectories that were identified by the ImageJ “TrackMate” plugin that were at least 6.8 µm (100 pixels) long, divided according to their initial location (center or periphery) and direction (center or cell membrane) as indicated above the graphs. The single dot represents the mean value from one measured cell. *N* = 4 independent experiments. *n* = 25 cells for both RAPA and DMSO. \**p* < 0.05, *ns*, nonsignificant (Mann-Whitney test).

**Fig. 8.**
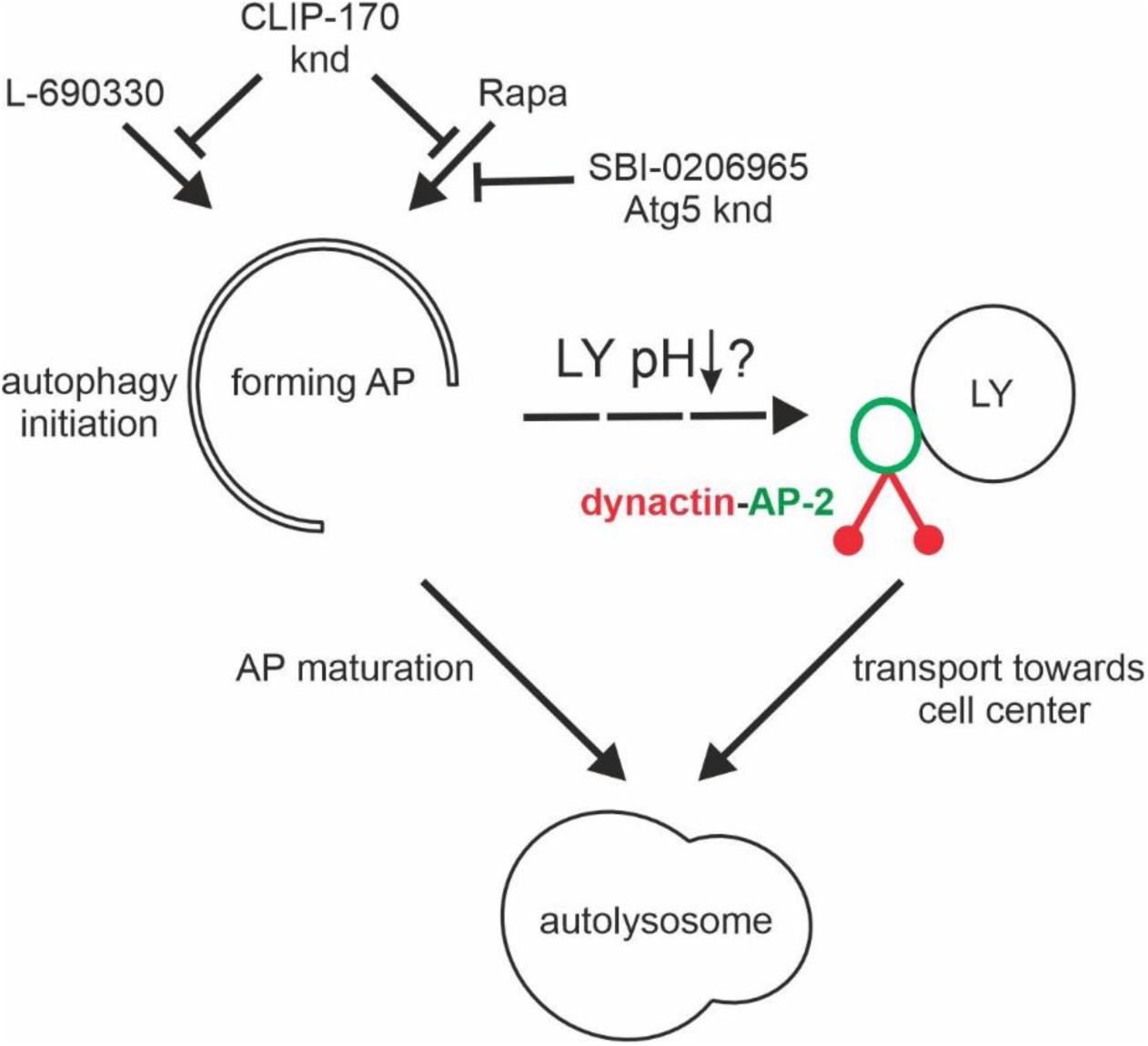
Postulated mechanism of autophagy initiation-induced recruitment of p150Glued and AP -2β to lysosomes. Administration of rapamycin or L-69030 initiates autophagosome formation (AP), which depends on the presence of CLIP -170. Simultaneously, autophagy inducers lead to a decrease in lysosomal pH (LY pH) and recruitment of the dynatin-AP2 complex to the lysosome (LY), presumably leading to intensification of LY retrograde transport to the perinuclear region and subsequent LY fusion with the mature AP. knd – knockdown, SBI-0206965 – Ulk inhibitor.

We found that rapamycin significantly increased lysosome acidification (**Fig. S11**), and lysosome acidification was previously related to their position in the cell. Therefore, we investigated whether the rapamycin-induced p150^Glued^–AP-2β interaction contributes to the mobility of lysosomes. We performed the live imaging of Rat2 cells that were transfected with Lamp-1-GFP and treated with 100 nM rapamycin for 2 h. Two distinct populations of Lamp-1-GFP objects were spotted. One aggregated near the cell center, and the other was more motile around the peripheries. When we focused on the latter one, we observed that rapamycin increased the speed of long-distance movements (> 6 µm) toward the cell center. Movement from the central Lamp-1-positive organelle pool toward the cell periphery was unaffected (**Fig. 7D, E**).

## Discussion

Recent work demonstrated that AP-2 is an adaptor protein for the dynein-dynactin transport of amphisomes along neuronal axons [22, 24]. To date, however, important questions about mechanistic details of the regulation of the dynactin–AP-2 interaction have not been answered. Here, we demonstrate that AP-2 and dynactin cooperate in neurons and non-neuronal cells under autophagy-permissive conditions, including, but not limited to, mTOR inactivation. Furthermore, we show that the co-occurrence of AP-2β with p150^Glued^ does not require binding the latter to microtubule plus ends or microtubule dynamics. However, this interaction requires the presence of CLIP-170, which contributes to autophagy initiation. Finally, we show that the autophagy-induced p150^Glued^–AP-2β interaction likely occurs on lysosomes, possibly increasing their mobility toward autophagosomes in the perinuclear area.

### Autophagy induction and a properly functioning endolysosomal pathway are essential for the p150^Glued^–AP-2β interaction

The initial finding that stimulated our research was the observation that rapamycin enhanced AP-2–dynactin complex formation in neurons. This discovery raised the issue of how mTORC1 inhibition stimulates this interaction. The present findings show that AP-2β or p150^Glued^ is not an mTORC1 substrate, instead supporting the scenario that mTORC1 inhibition promotes p150^Glued^–AP-2β complex formation via autophagy initiation (**Fig. 4**). Moreover, the pharmacological induction of autophagy that does not involve mTORC1 inhibition was sufficient to induce this interaction (**Fig. 5**), indicating that the direct trigger for the p150^Glued^– AP-2 interaction is autophagy. Furthermore, it explains why the coexistence of AP-2β and p150^Glued^ is readily visible in axons, whereas it is barely detectable in non-neuronal cells under basal conditions. In cultured neurons, axonal autophagy is relatively high and stable at a steady-state level. Autophagosomes are continuously formed at the axonal growth cone or at presynaptic sites and likely fuse with late endosomes, forming amphisomes, and are transported toward the cell soma [24, 28, 45, 82–84]. Thus, there is a continual need for AP-2–dynactin complexes in axons to transport amphisomes [24]. However, if axonal autophagy is constitutive and if autophagy is sufficient to drive the p150^Glued^–AP-2β interaction, then how does mTORC1 inhibition potentiate it? Although some studies show that mTORC1 inhibition does not increase autophagy in neurons [41, 42], other studies reported that it is nevertheless possible [43, 44]. Such observations show that this process can still be upregulated, despite its relatively high basal level. In contrast to neuronal axons, autophagy in many cultured non-neuronal cells is at a low level under basal conditions, and its induction requires additional stimuli (e.g., mTORC1 inactivation). Consequently, the degree of the p150^Glued^–AP-2β interaction in such cells is likely to be adjusted to the current level of autophagy in the cell. In the context of requiring the initiation of autophagy for the p150^Glued^–AP-2β interaction, it is also an interesting observation that this interaction requires the presence of CLIP-170. However, its important canonical functions (e.g., the regulation of microtubule dynamics) seem unnecessary. In contrast, we have shown that CLIP-170 is required for proper autophagy, but unclear is how CLIP-170 regulates this process. Thus, determining precisely how CLIP-170 regulates autophagy will require further studies.

Our previous experiments showed that AP-2β interacts with p150^Glued^ in neurons to transport amphisomes. Thus, we investigated whether unperturbed autophagosomal flux and the fusion of autophagosomes with organelles of the endolysosomal pathway are also required for this interaction in non-neuronal cells. Both inhibitors of these processes that we used (CQ and ABMA) significantly prevented the increase in the p150^Glued^–AP-2β PLA signal. This result may suggest that unperturbed autophagic flux is indeed crucial. Because both ABMA and CQ counteract the fusion of autophagosomes and amphisomes with lysosomes, this could suggest that this step is critical. Indeed, previous studies showed that AP-2 attaches to autolysosomes when autophagic lysosome reformation (ALR) is initiated. Autophagic lysosome reformation is a process that links autophagy, lysosomes, and AP-2 [85]. During ALR, protolysosomes emerge from autolysosomes and mature into lysosomes. Protolysosome formation requires clathrin and AP-2 [26]. Because ALR requires the merging of autophagosomes with the endolysosomal pathway, this would support the importance of this event for the p150^Glued^–AP-2β interaction. However, the lack of an effect of SNAP29 knockdown on the p150^Glued^–AP-2β interaction likely excludes this possibility. Thus, a question arises about how ABMA and CQ can block the p150^Glued^–AP-2β interaction. A common feature of these two compounds is their negative effect on the normal ultrastructure of the endolysosomal pathway [77, 78]. Our results clearly demonstrate that in Rat2 cells both ABMA and CQ lead to a significant decrease in lysotracker-positive organelles and a decrease in its intensity in the remaining ones. A similar effect of CQ on the pH of acidic organelles has been reported [80], although its absence has also been reported [78], suggesting that this effect may depend on the cell type. ABMA is a relatively recently described compound [77, 86], and its effect may also depend on the cell line and should always be tested experimentally. Thus, our results suggest that formation of the p150^Glued^–AP-2β complex requires proper function of the endocytic pathway and/or lysosome acidification. The latter possibility is further supported by the results of experiments with NH_4_Cl, which directly indicate that alkalinization of the cell prevents the increase in the p150^Glued^–AP-2β interaction upon rapamycin administration that initiates autophagy.

### Autophagy-induced p150^Glued^-AP-2β interaction occurs on lysosomes

Our results indicate that autophagy is a critical cellular process for the interaction of AP-2β with p150^Glued^. Nevertheless, our results did not identify autophagosomes as the primary p150^Glued^–AP-2β interaction site in non-neuronal cells. An important question is where the p150^Glued^–AP-2β interaction occurs upon autophagy induction. As mentioned above, the AP-2– dynactin complex in neurons is involved in the transport of signaling amphisomes in axons [24]. However, in non-neuronal cells, amphisomes are considered temporary structures [32, 87]. Moreover, ABMA, which potentiates their formation [77], did not enhance the p150^Glued^– AP-2β interaction. Finally, our PLA-EM and PLA-AiryScan confocal microcopy findings in Rat2 cells revealed a p150^Glued^–AP-2β PLA signal that often co-occurred with lysosomes. Notably, the presence of lysosomal AP-2 was previously reported and not only in the context of ALR, which corroborates our observation [88]. Thus, another issue emerges about the purpose of p150^Glued^–AP-2β complex formation on lysosomes.

Korolchuk et al. [89] showed that the dynein-dynactin-dependent transport of lysosomes toward the cell nucleus under cell starvation conditions is crucial for their pH regulation and fusion with autophagosomes. Hence, recruitment of the p150^Glued^–AP-2β complex to lysosomes at the onset of autophagy may be designed to ensure the proper fusion of these two organelles at the end of the process. A similar mechanism was previously reported for another microtubular transport lysosomal adaptor, ALG2 [51]. Under amino acid starvation or mTOR inhibition conditions, ALG2 and dynein are recruited to lysosomes in a Ca^2+^-dependent manner for their retrograde transport. Previous studies showed that RILP protein is responsible for lysosome transport by dynein-dynactin in response to changes in cholesterol levels [90]. This suggests that non-neuronal cells use several transport systems for one organelle, depending on the cellular conditions. This raises the question about what could trigger p150^Glued^–AP-2β complex binding to lysosomes upon mTOR inhibition. Based on our findings that CQ and ABMA decreased lysotracker fluorescence intensity, a tempting speculation is that changes in pH could serve as one such mechanism, but further research is needed to confirm this possibility.

In summary, our study provides new insights into the mechanisms that regulate formation of the p150^Glued^–AP-2β complex, which is essential for cargo transport along microtubules. Importantly, we showed that autophagy initiation is necessary and sufficient to trigger the formation of this complex. This finding exemplifies a basic mechanism that allows the coordination of various elements that are involved in a vital cellular process.

## Abbreviations

ABMA: 1-adamantyl(5-bromo-2-methoxybenzyl)amine
ANOVA: analysis of variance
ALR: autophagic lysosome reformation
Arp-1: actin-related protein 1
AP-2: adaptor protein 2
Baf A1: bafilomycin A1
BSA: bovine serum albumin
CHAPS: 3-([3-cholamidopropyl]dimethylammonio)-1-propanesulfonate
CHX: cycloheximide
CLIP-115: cytoplasmic linker protein 115
CLIP-170: cytoplasmic linker protein 170
CQ: chloroquine
DIV: day *in vitro*
DMEM: Dulbecco’s modified Eagle’s medium
DMSO: dimethylsulfoxide
EB3: end binding protein 3
EDTA: ethylenediamine tetraacetic acid
EGFP: enhanced green fluorescent protein
EGTA: ethylene glycol-bis(β-aminoethyl ether)-*N*,*N*,*N’*,*N’*-tetraacetic acid
EM: electron microscopy
FBS: fetal bovine serum
GST: glutathione-*S*-transferase
IP: immunoprecipitation
HRP: horseradish peroxidase
Lamp-1: lysosomal-associated membrane protein 1
LC3: microtubule-associated protein 1A/1B-light chain 3
mTORC1: mammalian/mechanistic target of rapamycin complex 1
PBS: phosphate-buffered saline
PCR: polymerase chain reaction
PFA: paraformaldehyde
PLA: proximity ligation assay
S6: ribosomal protein S6
RILP: Rab-interacting lysosomal protein
SDS-PAGE: sodium dodecyl sulfate-polyacrylamide gel electrophoresis
+TIP: plus end tracking protein
TrkB: tropomyosin receptor kinase B
vATPase: vacuolar (H+)-adenosine triphosphatase

## Acknowledgements

The authors thank Dr. Anna Akhmanova, Dr. Juan Bonifacino, Dr. Hong Cao, Dr. Iwona Ciechomska, Dr. Casper Hoogenraad, Dr. Volker Haucke, Dr. Mark McNiven, Dr. Lukasz Swiech, and Dr. Agata Zieba-Wicher for reagents, Dr. Rafaella de Pace, Dr. Carlos Guardia, and Dr. Juan Bonifacino for their support with establishing the protocol for the live imaging of neurons, Dr. Iwona Cymerman for help with designing cloning strategy, Dr. Katarzyna Poleszak for help with establishing protein production protocols, and Dr. Alexander Heberle for help with analyzing existing mTOR phosphoproteome datasets. We are also grateful to Dr. Iwona Ciechomska, Dr. Kathrin Thedieck, Dr. Viktor Korolchuk, Dr. Anne Spang for comments and discussions. We also thank Alina Zielinska and Marek Sarnacki from our laboratory and Tomasz Wegierski from the Microscopy and Flow Cytometry Core Facility at IIMCB for technical assistance and support, Angelika Jocek for laboratory management logistics, and Michael Arends for proofreading the manuscript.

## Authors Contributions

AT, KB, AB, JW, AM, and JJ designed the experiments. AT, KB, AB, JW, MMa, OT, AL, MC-K, TB, AAS, KO, MMl, SK, MB, and AM performed the experiments. AT, KB, AB, JW, MMa, OT, TB, MB, TR, AM, and JJ analyzed the data. AT, KB, AB, JW, AM, and JJ wrote the manuscript. All authors read and approved the manuscript.

## Data availability

All data generated or analyzed during this study are included in this published article and its supplementary materials. Raw data from all quantitatively analyzed experiments are available from the corresponding author upon reasonable request.

## Supplementary Information

### Supplementary Materials and Methods

#### Plasmids

β-actin-GFP-p150^Glued^ and β-actin-tdTomato-p150^Glued^ were obtained by polymerase chain reaction (PCR)-based subcloning of the p150^Glued^ coding sequence into SalI/NotI sites of β-actin-GFP and β-actin-tdTomato, respectively. The pAvi-tag-thrombin-HA-p150^Glued^N plasmid that encoded a fragment of p150^Glued^ (aa 1-490) was obtained by PCR using the following primers: 5’-GAAGAATTCATGGCCCAGAGCAAGAGGAC-3’ and 5’- GCGGGCGGCCGCTTAGGCGGCTTCCACTCGCTTCTG-3’, with pEGFPC2-Avi-tag- p150^Glued^ as a template. The resulting product was inserted into EcoRI/NotI restriction sites of pAvi-tag-thrombin-HA. pAvi-tag-thrombin-HA-p150^Glued^C plasmid that encoded a fragment of p150^Glued^ (aa 490-1278) was generated analogically using the following primers: 5’- GAAGAATTCATGGCAGGCGCCCGAGTAAGG-3’ and 5’- GCGGGCGGCCGCTTAGGAGATGAGACGACCGTGAAG-3’. The pAvi-tag-thrombin- HA-p150^Glued^C2 plasmid that encoded a fragment of p150^Glued^ (aa 1049-1278) was generated using 5’-GAAGAATTCATGGGCACTCCTGGGCAGGCTCCAGGCGC-3’ and 5’- GCGGGCGGCCGCTTAGGAGATGAGACGACCGTGAAG-3’ primers, with pAvi-tag-thrombin-HA-p150^Glued^C as a template. The resulting product was inserted into EcoRI/NotI restriction sites of pAvi-tag-thrombin-HA. The GST-p150^Glued^C plasmid that encoded a fragment of p150^Glued^ (aa 490-1278) with a GST tag was generated by PCR from the HA-Avi- tag-p150^Glued^C plasmid using 5’-GAAGAATTCATGGCAGGCGCCCGAGTAAGG-3’ and 5’-GCGGGCGGCCGCTTAGGAGATGAGACGACCGTGAAG-3’ primers. The product was inserted in pGEX-4T1 into EcoRI/NotI restriction sites. The GST-p150^Glued^C2 plasmid that encoded a fragment of p150^Glued^ (aa 1049-1278) with a GST tag was obtained by PCR analogically using 5’-GAAGAATTCATGGGCACTCCTGGGCAGGCTCCAGGCGC-3’ and 5’-GCGGGCGGCCGCTTAGGAGATGAGACGACCGTGAAG-3’ primers.

#### Antibodies

**Table S1.** Primary antibodies used for the study.

| Primary antibody | Manufacturer, catalog no. | Application, dilution |
| --- | --- | --- |
| mouse anti-p150<sup>Glued</sup> | BD Bioscience, 610474 | PLA 1:100<br>WB 1:1000<br>IF 1:200<br>IP 2 µg/reaction |
| rabbit anti-p150<sup>Glued</sup> | Cell Signaling Technology, 69399 | WB 1:1000 |
| rabbit anti-AP-2β | Thermo Fisher, PA1-1066 | PLA 1:100<br>IP 2 µg/reaction<br>IF 1:250 |
| rabbit anti-AP-2β | Santa Cruz Biotechnology, sc-10762 | WB 1:500 |
| mouse anti-AP-2β | Santa Cruz Biotechnology, sc-74423 | WB 1:500 |
| goat anti-AP-2β | Santa Cruz Biotechnology, sc-6425 | IF 1:200<br>IP 10 µg/reaction |
| mouse anti-AP-2α | BD Bioscience, 610502 | WB 1:500 |
| mouse anti-Arp1 | Santa Cruz Biotechnology, sc-376010 | WB 1:250 |
| mouse anti-p62 | Santa Cruz Biotechnology, sc-55603 | WB 1:250 |
| mouse anti-DIC1/2 | Santa Cruz Biotechnology, sc-13524 | WB 1:250 |
| Rabbit anti-phospho-S6 Ser235/236 | Cell Signaling Technology, 4858 | WB 1:1000<br>IF 1:200 |
| mouse anti-S6 | Cell Signaling Technology, 2217 | WB 1:1000 |
| mouse anti-CLIP-170 | Santa Cruz Biotechnology, 28325 | WB 1:500<br>IF 1:500 |
| rabbit anti-GFP | MBP, 598 | WB 1:1000 |
| mouse anti-tubulin | Sigma-Aldrich, T5168 | WB 1:5000<br>IF 1:500 |
| rabbit anti-tubulin | Abcam, ab4074 | WB 1:3000 |
| rabbit anti-LC3B | Sigma-Aldrich, L7543 | WB 1:2500<br>IF 1:200 |
| rabbit anti-LC3B | Cell Signaling Technology, 2775 | IF 1:250 |
| mouse anti-Lamp1 | DSHB, H4A3 | IF 1:100 |
| mouse anti-beclin-1/BECN1 | Santa Cruz Biotechnology, sc-48341 | WB 1:250 |
| rabbit anti-phospho-beclin-1 Ser30 | Cell Signaling Technology, 54101 | WB 1:500 |
| rabbit anti-p62/SQSTM1 | Cell Signaling Technology, 5114 | WB 1:500<br>IF 1:250 |
| mouse anti-Atg5 | Cell Signaling Technology, 12994 | WB: 1:500 |
| mouse anti-actin | Sigma-Aldrich, A4700 | WB 1:1000 |
| mouse anti-GAPDH | Sigma-Aldrich, MAB374 | WB 1:500 |
| rabbit IgG | Cell Signaling Technology, 3900 | IP 2 µg/reaction |
| goat IgG | Sigma-Aldrich, I5256-10MG | IP 10 µg/reaction |
| rabbit anti-neurofascin | Antibodies Incorporated, 75-172 | Live imaging 1:100 |
| mouse anti-his-tag | Cell Signaling Technology, 2366 | WB 1:1000 |
| rat anti-GST-tag | Chromotek, 6g9-100 | WB 1:1000 |
| rabbit anti-HA-tag | Cell Signaling Technology, 3724 | WB 1:1000 |

PLA, proximity ligation assay; WB, Western blot; IF, immunofluorescence; IP, immunoprecipitation.

#### Cell line cultures and transfection

Rat2 and HEK293T cells were purchased from the American Type Culture Collection and grown in Dulbecco’s modified Eagle’s medium (DMEM) with 4500 mg/ml of D-glucose that contained 10% fetal bovine serum (FBS) and 1% penicillin-streptomycin (all from Sigma-Aldrich) at 37°C in a 5% CO_2_ atmosphere. For the proximity ligation assay (PLA) and PLA-electron microscopy (EM) experiments, Rat2 cells were grown on glass coverslips or Thermanox plastic coverslips (Thermo Fisher; catalog no. 174950), respectively, coated with 0.2% gelatin for 1 h at 37°C. For live experiments, Rat2 cells were seeded on glass coverslips that were coated with poly-L-lysine (50 μg/ml in H_2_O for 1 h). For plasmid DNA transfection, HEK293T cells at 70% confluency were transfected using polyethylenimine PEI 25K (Polysciences, catalog no. 23966) according to the manufacturer’s protocols. After transfection, HEK293T cells were grown in DMEM that was supplemented with 5% FBS for 48 h. Rat2 cells were transfected with plasmid DNA using electroporation. For each transfection, a total of 10^6^ cells were suspended in Opti-MEM medium (Thermo Fisher, catalog no. 31985-047), mixed with 10 μg of DNA, and added to 2 mm gap cuvettes (Nepagene, catalog no. EC-002S). Cells were electroporated using a NEPA21 electroporator (Nepagene) with poring pulses (6×, 150 V, 2.5 ms length, 50 ms interval, 10% decay rate), followed by transfer pulses (5×, 20 V, 50 ms length, 50 ms interval, five pulses, 40% decay rate) with “±” polarity set for both. After electroporation, the cells were grown in DMEM with 10% FBS without antibiotics. The medium was changed the next day for a medium that contained 1% penicillin-streptomycin. The siRNA transfection of HEK293T and Rat2 cells was performed on trypsinized cells using Lipofectamine RNAiMAX Transfection Reagent (Thermo Fisher, catalog no. 13778150) according to the manufacturer’s protocol. For the combined siRNA transfection/DNA electroporation experiments, cells after siRNA transfection were seeded on 10 cm plates and allowed to attach. After 24 h, cells were harvested by trypsinization and electroporated as described earlier. After electroporation, each variant was seeded on glass coverslips that were coated with poly-L-lysine (50 μg/ml in H_2_O for 1 h).

#### Primary neuron preparation and transfection

Primary hippocampal cultures were prepared from embryonic day 18 rat brains as described previously [1]. The rats that were used to obtain neurons for further experiments were sacrificed according to protocols that complied with European Community Council Directive 2010/63/EU. Transfections of hippocampal neurons were performed on day *in vitro* 5 (DIV5) with Lipofectamine2000 (Thermo Fisher, catalog no. 11668019) as described previously [1], except that the incubation time with the transfection mixture was reduced to 2 h. DNA (2 µg) and 1.5 µl of Lipofectamine2000 were used per well of a 12-well dish.

#### Proximity ligation assay

For the PLA, cells were fixed for 5 min with ice-cold 100% methanol and 2 mM ethylene glycol-bis(β-aminoethyl ether)-*N*,*N*,*N’*,*N’*-tetraacetic acid (EGTA) at -20°C, followed by 10 min with 4% paraformaldehyde (PFA)/4% sucrose in phosphate-buffer (pH 7.4). Fixed cells were washed three times with phosphate-buffered saline (PBS) and incubated with mouse anti-p150^Glued^ and rabbit anti-AP-2β antibodies that were diluted in PBS that contained 1% donkey serum and 0.2% Triton-X100 at 4°C overnight. The next day, the cells were washed twice for 5 min with PBS with 1% donkey serum and 0.2% Triton-X100, washed once for 1 min in antibody diluent that was provided in the manufacturer’s PLA kit (Sigma-Aldrich, catalog no. DUO92101), and then incubated for 60 min at 37°C with relevant secondary antibodies that were conjugated to oligonucleotide PLUS or MINUS (Sigma-Aldrich; catalog no. DUO92002 and DUO92004, respectively) and diluted in antibody diluent. The coverslips were then washed twice for 5 min with buffer A (Sigma-Aldrich, catalog no. DUO82046). Ligation and amplification were performed according to the manufacturer’s protocol using Duolink In Situ Detection Reagents Red (Sigma-Aldrich, catalog no. DUO92008). The cells were then counterstained with mouse anti-tubulin antibody that was diluted in PBS that contained 1% donkey serum and 0.2% Triton-X100 for 30 min at room temperature and washed three times with PBS and incubated with anti-mouse Alexa-488-conjugated secondary antibody that was diluted in PBS that contained 1% donkey serum and 0.2% Triton-X100 for 30 min at room temperature to visualize cell shape. Finally, the cells were washed three times with PBS and mounted with DuoLink *in situ* mounting medium that contained DAPI (Sigma-Aldrich, catalog no. DUO82040). For each experiment, negative controls that lacked one of the primary antibodies were used.

#### Immunofluorescence

For analyses of the co-localization of p150^Glued^ and AP-2β proteins, Rat2 cells were fixed with ice-cold 100% methanol with 2 mM EGTA for 5 min at -20°C and then with 4% PFA/4% sucrose in phosphate-buffer (pH 7.4) for 10 min at room temperature and washed three times with PBS. After fixation, the cells were blocked for 1 h in blocking buffer (5% donkey serum, 0.3% Triton X-1001 in PBS). Next, the cells were incubated overnight with primary antibodies in antibody dilution buffer (1% BSA, 0.3% Triton X-100 in PBS) at 4°C and then washed three times with PBS at room temperature. Afterward, specimens were incubated with Alexa 488- and Alexa 594-conjugated secondary antibodies for 1 h at room temperature, followed by three washes with PBS. Coverslips were mounted with Prolong Gold with DAPI (Thermo Fisher, catalog no. P36941). For the immunofluorescent staining of ribosomal protein S6 phosphorylated at Ser235/236 (P-S6), neurons were fixed with 4% PFA/4% sucrose in phosphate-buffer (pH 7.4) for 10 min at room temperature and washed three times with PBS. After fixation, the cells were blocked for 1 h in blocking buffer (5% donkey serum, 0.3% Triton X-1001 in PBS). Next, the cells were incubated overnight with primary antibody in antibody dilution buffer I (1% BSA, 0.3% Triton X-100 in PBS) at 4°C and then washed three times with PBS at room temperature. Alexa 488-conjugated secondary antibody with Alexa 568-conjugated phalloidin in antibody dilution buffer was next added for 1 h at room temperature. Afterward, the cells were washed three times with PBS, and coverslips were mounted with Prolong Gold (Thermo Fisher, catalog no. P36934). For the immunofluorescent detection of CLIP-170, the cells were fixed for 5 min with ice-cold 100% methanol and 2 mM EGTA at - 20°C, followed by 10 min with 4% PFA/4% sucrose in phosphate-buffer (pH 7.4) at room temperature, and then washed three times with PBS. Next, the cells were washed twice with antibody dilution buffer II (1% donkey serum, 0.2% Triton X-100 in PBS) and incubated overnight with the primary antibody in antibody dilution buffer II at 4°C. After incubation with primary antibodies, the cells were washed three times with PBS at room temperature and incubated with Alexa 488-conjugated secondary antibody in antibody dilution buffer II for 1 h at room temperature. Afterward, the cells were washed three times with PBS, and nuclei were stained with Hoechst 33258 (1 µg/ml; Thermo Fisher Scientific) for 5 min. The cells were then washed two more times with PBS, and coverslips were mounted with Prolong Gold. To analyze LC3B immunofluorescence, the cells were fixed according to the CLIP-170 staining protocol. After fixation, the cells were blocked for 15 min at room temperature in antibody dilution buffer III (2% donkey serum, 0.2% Triton X-100 in PBS) and incubated for 1 h with the primary antibody in antibody dilution buffer III at room temperature. The cells were then washed three times with antibody dilution buffer III at room temperature and incubated with Alexa 488-conjugated secondary antibody in antibody dilution buffer III for 1 h at room temperature. The next steps were identical to the description of CLIP-170 immunofluorescence.

#### Fixed cell image acquisition and analysis

Microscopic images of fluorescently labeled *in vitro* cultured cells, with the exception of quantitative co-localization experiments, were acquired using a Zeiss LSM800 confocal microscope (40× or 63× Plan Apo oil immersion objective, NA=1.4) as Z-stacks at 0.2-0.5 μm intervals at 1024 × 1024 pixel resolution. To obtain higher image resolution and conduct studies of protein co-localization, an AiryScan detector was used with the 63× oil objective. To enable comparisons between images, the microscope settings were kept constant for all scans. The Z-stacks, with the exception of images acquired in AiryScan mode, were converted to single images using a maximum intensity projection. For quantification of the number of PLA puncta in Rat2 cells, the cell shape was indicated as a region of interest for each cell, and the threshold was set manually and uniformly for all images in each experiment to extract specific signals from background. The number of PLA puncta per cell was counted in ImageJ software using the “Analyze Particles” macro. The average number of PLA puncta per cell was then calculated for each condition, and the results were normalized by subtracting the average number of PLA puncta per cell from the negative control. The obtained average number of PLA puncta per cell was further normalized by dividing the obtained values from a particular experiment by the average number of PLA puncta per cell from the control from at least three repetitions. For the analysis of P-S6 levels in neurons, the mean fluorescent intensity of P-S6 was measured in the neuronal soma using ImageJ software.

For quantitative co-localization analysis, images of fluorescently labeled cells were registered using a Leica SP8 confocal microscope with a 63× oil immersion Plan Apo objective lens (NA=1.4). The microscope system was equipped with acousto-optical beam splitter and multialkali single-channel PMTs and hybrid detectors (HyD). The fluorescence of DAPI was excited with a 405 nm light (5 mW diode laser) and detected in the 410-470 nm range. The fluorescence of Alexa 488 and Alexa 568 was excited with a supercontinuum (white light) laser (15 mW) at their absorption maxima and detected at 490-550 nm and 575-630 nm ranges, respectively. The images (optical sections) were registered using PMT (gain 600, DAPI) or HyD (gain 200, Alexa) working in the integration mode at 16-bit precision. The pixel size was 90 nm in the x-y plane and 350 nm along the optical axis. The pixel dwell time was 2.5 µs (4× averaging), and the confocal pinhole was set to 1 Airy unit (at 530 nm). The cells were imaged at room temperature. Stacks of optical sections (Alexa 488 and Alexa 568) were subjected to blind deconvolution (15 iterations) using Huyghens v. 3.7 (Scientific Volume Imaging, Hilversum, The Netherlands). Processing was initialized with the nominal PSF of the objective and performed at an SNR set to 15. The background was estimated as a minimum of average intensity in a 25 × 25 region. Following deconvolution, the nuclei (DAPI) were segmented using Otsu thresholding (two classes). Overlapping nuclei were separated by Euclidean distance transform, followed by watershedding. The Alexa 488 and Alexa 568 images were summed, processed with a median filter (7 × 7 × 3 size) and dilated (11 × 11 × 3 neighborhood). The cells were segmented using Otsu thresholding, and single cell masks were constructed with iterative dilation of the nuclear masks. The dilation steps were ordered by the intensity of processed cell images. Volumes that corresponded to nuclei were then excluded from the cell masks. Coefficients of intensity correlation (Pearson and Spearman) between Alexa 488 and Alexa 568 images were calculated on a cell-by-cell basis using the respective masks. The operation was repeated using the Alexa 568 images that were shifted by ±8 voxels in the x and y directions and ±2 voxels in the z direction. The respective coefficients were calculated using the union of two respective cellular masks and corresponded to the random association of Alexa 488 and Alexa 568 fluorescence in a cell. The raw correlation coefficients were then divided by their random counterparts (on a cell-by-cell basis) to create standardized Pearson and Spearman values.

#### Opera Phenix High content imaging of fixed cells

For the high-throughput analysis of effects of different drugs on lysosomal acidity, Rat2 cells were seeded on CellCarrier-96 well Black glass bottom plates (Perkin Elmer) that were coated with 0.2% gelatin at a density of 7 × 10^3^ cells per well 1 day before treatment (see above). After 80 min of drug exposure, LysoTracker Red DND-99 (500 nM, Invitrogen, catalog no. L7528) was added to the cells for the last 40 min. Subsequently, the cells were fixed with 4% PFA for 10 min and subjected to three 10-min washes with PBS. High-content screening microscopy was conducted using the Opera Phenix system (PerkinElmer) that was equipped with a 40× 1.1 NA water immersion objective. Image acquisition and subsequent analysis were performed using Harmony 4.9 software (PerkinElmer). Statistical analyses were performed using the R and RStudio software packages (Cran). To ensure consistent and reliable data analysis, cells were initially filtered based on morphological criteria to obtain uniform cell populations. The identification of lysotracker-positive compartments was achieved using the “Find spots” algorithm that is integrated within the Harmony software. Additionally, the total signal intensity of Lysotracker staining was quantified within both the cell cytoplasm and the previously defined compartment regions.

#### Protein production in bacteria and pull-down experiment

Recombinant proteins were produced in the *E. coli* BL21 strain. Individual clones that were transformed with plasmids that encoded proteins of interest were picked from the plates and inoculated to 5 ml of LB with appropriate antibiotic. After overnight culture, at 37°C with shaking, bacteria were refreshed with new medium in a 1:50 ratio and further cultured until reaching an optical density at 600 nm (OD_600_) of 0.6-0.8. Protein production was induced using 1 mM isopropyl-β-D-1-thiogalactopyranoside (Carl Roth, catalog no. CN08.3). The His_6_-AP-2β appendage domain, GST-Eps15, and free GST-tag were produced at 37°C for 2 h, and GST-p150^Glued^ fragments were produced at 21°C overnight. For protein purification, cultures were centrifuged at 4,500 × *g* for 10 min, and the pellet was resuspended in lysis buffer (50 mM Tris [pH, 8.0], 150 mM NaCl, 0.1% Triton X-100, and protease inhibitors). For all subsequent stages, lysates were kept on ice. After resuspension, the cells were lysed by sonication in a Sonics VCX130 PB sonicator (Vibra-Cell) in two 30-s sessions with a 70% amplitude. After sonication, lysates were centrifuged at 13,000 × *g* for 5 min to remove the insoluble fraction. For the pull-down experiment, the bait protein was then added to Glutathione-Sepharose 4B resin (Merck, catalog no. GE17-0756-01) at a ratio of 30 μl beads for each 10 ml of original bacterial culture and incubated for 1 h with end-to-end rotation. After incubation, the resin was washed three times with lysis buffer, and the prey protein lysate was added. After another 1 h of incubation, the resin was again washed three times with lysis buffer and prepared for Western blot by dissolving in 1× Laemmli buffer and incubation in a heat block at 94°C for 15 min.

#### Whole-cell lysate preparation

For the Western blot analysis of proteins in whole-cell lysates, Rat-2 cells were lysed in lysis buffer (20 mM Tris [pH 7.5], 150 mM NaCl, 2 mM ethylenediamine tetraacetic acid [EDTA], 0.5% Triton X-100, 0.5% NP-40, 2 mM MgCl_2_, and 10% glycerol supplemented with protease and phosphatase inhibitors). The total protein concentration in whole-cell lysates was measured using the Pierce BCA Protein Assay Kit (Thermo Fisher, catalog no. 23225) according to the manufacturer’s instructions. Afterward, 4× Laemmli sample buffer was added to the lysates, followed by boiling for 5 min at 95°C.

#### Immunoprecipitation

For the immunoprecipitation (IP) of endogenous proteins from HEK293T cells, the cells were lysed for 15 min on ice in lysis buffer (20 mM HEPES [pH 7.5], 120 mM KCl, and 0.3% 3-[(3-cholamidopropyl)dimethylammonio]-1-propanesulfonate [CHAPS] supplemented with protease and phosphatase inhibitors, in addition to 100 nM rapamycin for variants in which cells were treated with rapamycin) and spun. Next, 2 µg of anti-AP-2β or anti-p150^Glued^ primary antibody or IgG (as a negative control) was added to the supernatant and incubated overnight at 4°C while rotating. The next day, a mixture of lysate and antibodies was added to 30 µl of Dynabeads Protein G (Thermo Fisher, catalog no. 10004D) and incubated for 4 h at 4°C while rotating. Beads were then washed four times in wash buffer (20 mM HEPES [pH 7.5], 120 mM KCl, and 0.1% CHAPS), eluted in 2× Laemmli sample buffer, incubated for 5 min in 95°C and analyzed by sodium dodecyl sulfate-polyacrylamide gel electrophoresis (SDS-PAGE), followed by Western blot. The IP of AP-2β from rat brain extracts was performed as described previously [2]. For the IP of heterologous proteins, HEK293T cells were transfected with pEGFPC1-Ap2b1, GFP-β-actin-p150^Glued^, or pEGFPC1 (as a negative control). Forty-eight hours after transfection, the cells were lysed for 15 min on ice in lysis buffer (10 mM Tris [pH 7.5], 150 mM NaCl, 0.5 mM EDTA, and 0.15 % CHAPS supplemented with protease and phosphatase inhibitors) and spun using a benchtop centrifuge at maximum speed for 15 min at 4°C. The lysates were then incubated with GFP-Trap Agarose (20 µl per variant; Chromotek, catalog no. gta-10) for 2 h at room temperature, followed by four washes with wash buffer (10 mM Tris [pH 7.5], 150 mM NaCl, and 0.5 mM EDTA). The immunoprecipitated proteins were next used for the mTOR kinase assay.

#### Avi-tag pull down of biotinylated proteins and AP-2β-ear binding

The His_6_-AP-2β appendage domain was produced as described in the *Protein production in bacteria* section. For protein purification, cultures were centrifuged at 4,500 × *g* for 10 min, and the pellet was resuspended in lysis buffer (50 mM NaH_2_PO_4_/Na_2_HPO_4_ [pH 8], 300 mM NaCl, 10 mM imidazole, 10% glycerol, 10 mM β-mercaptoethanol, 0.1% Triton X-100, and protease inhibitors). For all subsequent stages, lysates were kept on ice. After resuspension, the cells were lysed by sonication in a Sonics VCX130 PB sonicator (Vibra-Cell) in two 25-s sessions with a 70% amplitude. After sonication, the lysates were centrifuged at 13,000 × *g* for 5 min to remove the insoluble fraction. The column with 5 ml agarose - Ni - NTA used for affinity chromatography of recombinant protein was equilibrated with 20 ml of lysis buffer and a flow rate of approximately 1 ml/min. The cell lysate that was obtained from 2 L of culture was then added to the column at a flow rate of approximately 0.5 ml/min. After the lysate passed through the column, the agarose resin was washed successively with 50 ml of appropriate buffers: lysis buffer, lysis buffer supplemented with 2 M NaCl, and lysis buffer supplemented with 20 mM imidazole. The protein was then eluted with 25 ml of elution buffer (50 mM NaH_2_PO_4_/Na_2_HPO_4_ [pH 8], 2 M NaCl, 250 mM imidazole, 10% glycerol, 10 mM β-mercaptoethanol 0.1% Triton X-100). One milliliter fractions were collected and verified for the presence of purified protein. Those that contained the highest amount of the His_6_-AP-2β ear domain were pooled and used for the binding experiments. M-280 streptavidin Dynabeads (Thermo Fisher, catalog no. 11205D) were incubated for 1 h with blocking buffer (20 mM HEPES [pH 7.5], 120 mM KCl, 0.5 mg/ml BSA, and 20% glycerol). The beads were washed with lysis buffer (20 mM HEPES [pH 7.5], 120 mM KCl, and 0.3% CHAPS supplemented with protease and phosphatase inhibitors). Forty-eight hours after transfection with BirA and plasmids that encoded Avi-tagged proteins, HEK293T cells were lysed for 15 min on ice in lysis buffer and spun using a benchtop centrifuge at maximum speed (15 min, 4°C). The supernatant was added to the previously prepared M-280 streptavidin Dynabeads and incubated together for 1 h at 4°C while rotating. The beads were then washed on ice four times in wash buffer (20 mM HEPES [pH 7.5], 500 mM KCl, and 0.1% CHAPS) and incubated with 0.2 mg/ml His_6_-AP-2β ear domain purified from bacteria in binding buffer (20 mM HEPES [pH 7.5], 300 mM KCl, and 0.1% CHAPS) for 2 h at 4°C while rotating. The beads were then washed four times with binding buffer, eluted in 2× Laemmli sample buffer, incubated for 5 min in 95°C and analyzed by SDS-PAGE, followed by Western blot.

#### Western blot

Protein samples were analyzed by SDS-PAGE and immunoblotting according to standard laboratory protocols. Primary antibodies that were used for Western blot are listed in Table 1. Proteins of interest were detected using HRP- or IRdye-conjugated secondary antibodies in a chemo-luminescence reaction or with the Odyssey LiCor Biosciences system, respectively. The densitometry analysis of the amount of LC3B was performed using Image Studio Lite (LiCor Biosciences) and a previously described method [3]. Specifically, the signal intensities for LC3B I and LC3B II were first measured. Based on these values, normalization was performed. First, for all variants in a single replicate, the sum of intensities for a given protein was calculated. Second, individual protein band intensities were divided by this sum. Such normalization was performed for LC3B I and LC3B II. Finally, after normalizing the values, the ratio between LC3B II and LC3B I was calculated by dividing the normalized values of LC3B II by LC3B I.

#### Kinase assays

An *in vitro* phosphorylation assay was performed for 20 min at 30°C in kinase assay reaction buffer (25 mM HEPES [pH 7.5], 50 mM KCl, and 10 mM MgCl_2_) in the presence of 2.5 μCi [γ-^33^P]ATP (Hartmann Analytic) and 2.5 mM “cold” ATP using a recombinant, active mTOR kinase fragment (1362-2549 aa; Millipore, catalog no. 14-770) and proteins that were immunoprecipitated from HEK293T cells or commercial mTOR substrate (Merck Millipore, catalog no. 12-645). The reaction was stopped by adding 4× Laemmli sample buffer and boiling for 5 min at 95°C. Next, the samples were separated by SDS-PAGE. The gel was stained with Coomassie Brilliant Blue and dried. The radioactive signal was detected and analyzed using a Typhoon Trio+ Phosphorimager (GE Healthcare).

#### RNA Isolation for Quantitative Real-Time PCR

RNA was isolated from Rat2 cells using the RNeasy Mini Kit (Qiagen) and reverse transcribed with the High-Capacity cDNA Reverse Transcription Kit (Thermo Fisher Scientific). Quantitative real-time polymerase chain reaction (qRT-PCR) was performed using FAM fluorophore TaqMan probes with TaqMan Gene Expression Master Mix buffer (Applied Biosystems). TaqMan probes used for the experiments were the following: tubulin (Rn01532518_g1) and Snap29 (Rn00587482_m1; Thermo Fisher). Data were acquired with the LightCycler 96 System (Roche) and analyzed using the 2^(−ΔΔCt)^ method for relative quantification.

## Supplementary Figures

**Fig. S1.**
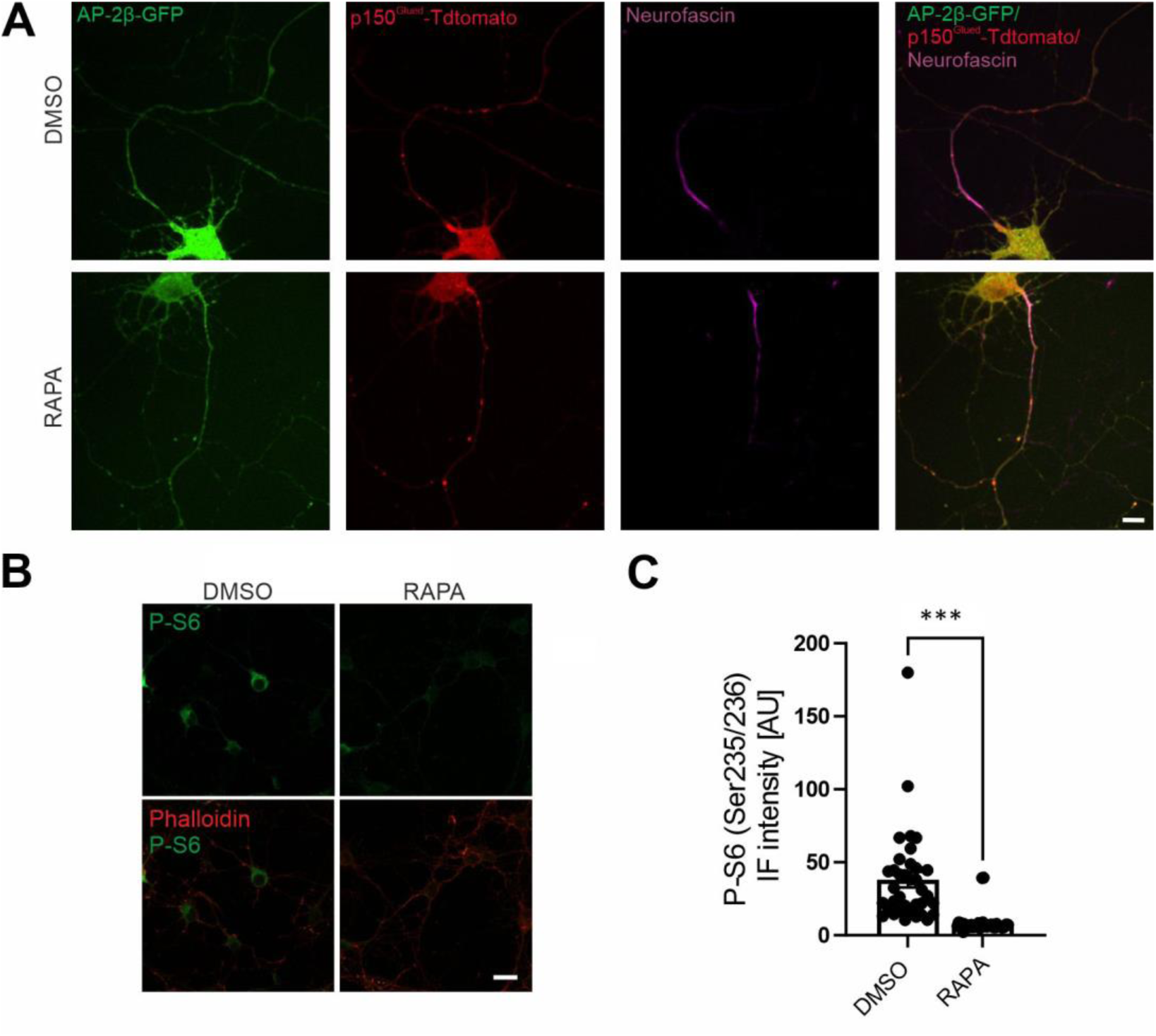
mTORC1 inhibition in neurons increases p150^Glued^–AP-2β interaction in axons. (**A**) Snapshots of DIV7 neurons that were treated as indicated and expressed p150^Glued^- Tdtomato (red) and AP-2β-GFP (green) and stained for the axon initial segment with CF640R-conjugated neurofascin antibody (magenta). Scale bar = 10 μm. (**B**) Representative pictures of DIV7 neurons that were treated as indicated and immunofluorescently labeled for endogenous P-S6 (Ser235/236) (green) and stained with Alexa Fluor 568 phalloidin. Scale bar = 20 µm. (**C**) Quantification of P-S6 (Ser235/236) immunofluorescence level in cell bodies of neurons that were treated as indicated. The data are expressed as the mean fluorescence intensity ± SEM. *N* = 2 independent experiments. *n* = 36 cells (DMSO), 39 cells (RAPA). *\*\*\*p* < 0.001 (Mann-Whitney test).

**Fig. S2.**
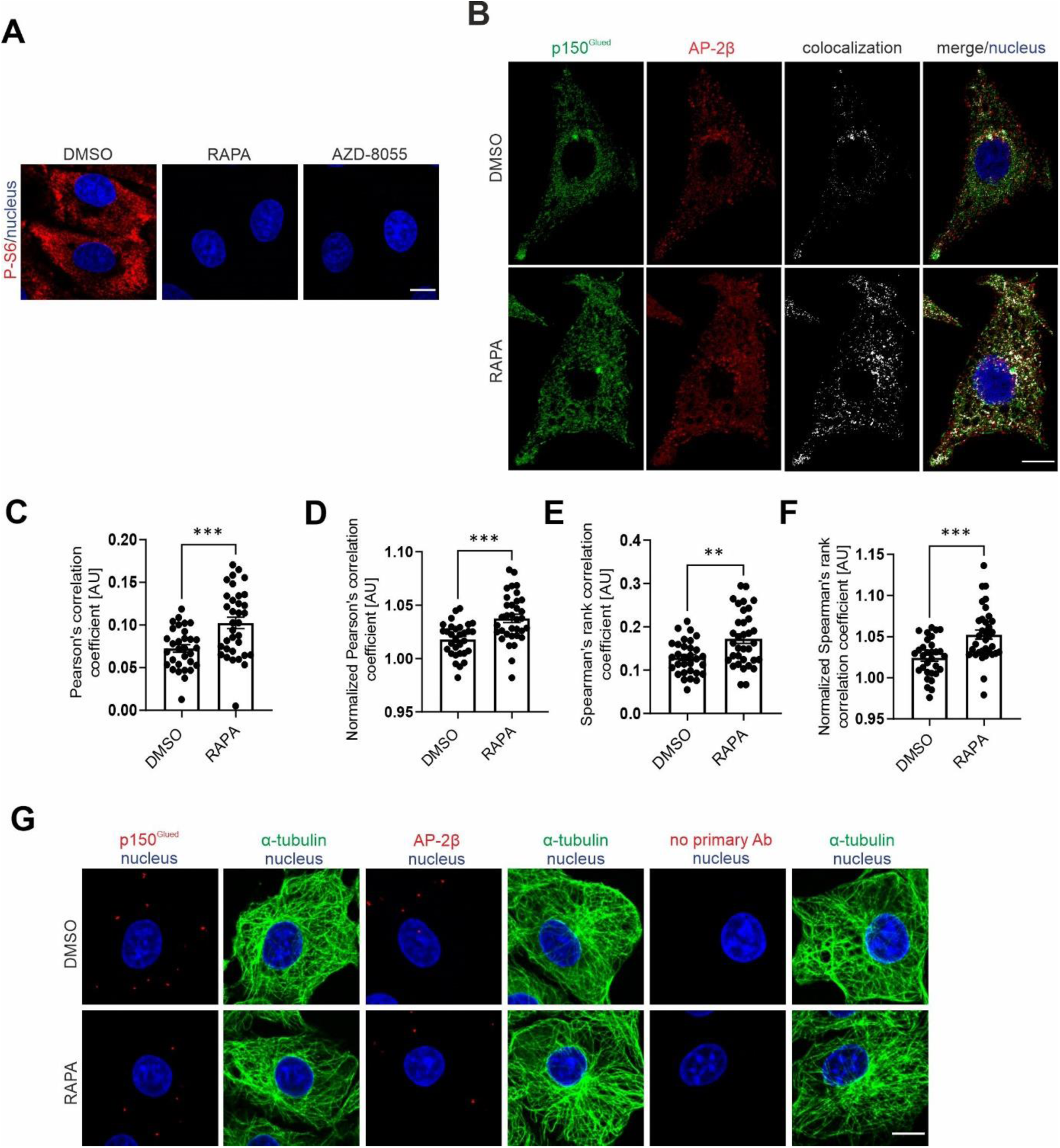
mTORC1 inhibition increases p150^Glued^–AP-2β interaction in non-neuronal cells. (**A**) Representative images of Rat2 cells that were treated for 2 h with 0.1% DMSO, 100 nM rapamycin (RAPA), or 100 nM AZD-8055 with immunofluorescently labeled endogenous P-S6 (Ser235/236) (red) and nuclei that were stained with Hoechst 33258 (blue). Scale bar = 10 µm. (**B**) Representative images of Rat2 cells that were treated for 2 h with 0.1% DMSO or 100 nM rapamycin (RAPA) with immunofluorescently labeled endogenous p150^Glued^ (green) and AP-2β (red) and additionally stained with DAPI (blue). Scale bar = 10 µm. The white channel indicates p150^Glued^ and AP-2β immunofluorescent signal co-localization. (**C**) Analysis of Pearson’s correlation coefficients of endogenous AP-2β/p150^Glued^ co-localization without normalization. The data are expressed as the mean coefficient of co-localization of two proteins ± SEM. *N* = 2 independent experiments. *n* = 31 cells (DMSO), 34 cells (RAPA). *\*\*\*p* < 0.001 (Student’s *t*-test). (**D**) Analysis of Pearson’s correlation coefficients of AP-2β/p150^Glued^ with normalization to the appropriate value that resulted from the random distribution of fluorescence in the cell. The data are expressed as the mean coefficient of co-localization of two proteins ± SEM. *N* = 2 independent experiments. *n* = 31 cells (DMSO), 34 cells (RAPA). *\*\*\*p* < 0.001 (Student’s *t*-test). (**E**) Analysis of Spearman’s rank correlation coefficient of AP-2β/p150^Glued^ co-localization without normalization. The data are expressed as the mean coefficient of co-localization of two proteins ± SEM. *N* = 2 independent experiments. *n* = 31 cells (DMSO), 34 cells (RAPA). *\*\*p* < 0.005 (Student’s *t*-test). (**F**) Analysis of Spearman’s rank correlation coefficient of AP-2β/p150^Glued^ co-localization with normalization to the appropriate value that resulted from the random distribution of fluorescence in the cell. The data are expressed as the mean coefficient of co-localization of two proteins ± SEM. *N* = 2 independent experiments. *n* = 31 cells (DMSO), 34 cells (RAPA). *\*\*\*p* < 0.001 (Student’s *t*-test). (**G**) Representative images of Rat2 fibroblasts with negative controls for the PLA with one primary antibody present (p150^Glued^ or AP-2β) or without primary antibodies (Ab) (red), with immunofluorescently labeled tubulin (green) and DAPI-stained nuclei (blue). Cells were treated for 2 h with 0.1% DMSO or 100 nM rapamycin (RAPA). Scale bar =10 µm.

**Fig. S3.**
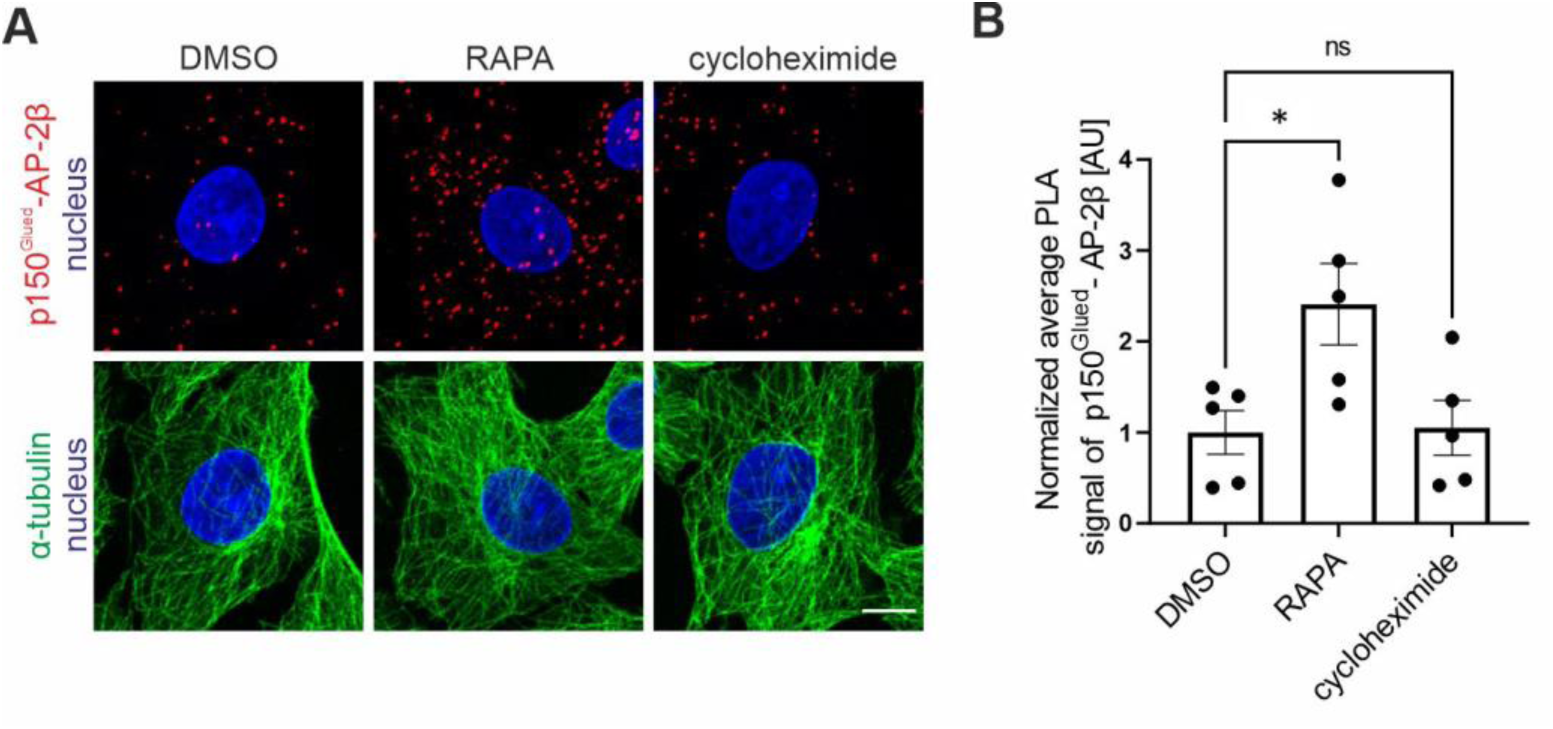
Protein synthesis inhibition does not enhance p150^Glued^–AP-2β interaction. (**A**) Representative images of Rat2 fibroblasts treated for 2 h with 0.1% DMSO, 100 nM rapamycin (RAPA), or 35 µM cycloheximide with p150^Glued^–AP-2β PLA signals (red), immunofluorescently labeled tubulin (green) and DAPI-stained nuclei (blue). Scale bar = 10 μm. (**B**) Quantification of the number of p150^Glued^–AP-2β PLA puncta in cells that were treated as in A. The data are expressed as the mean number of PLA puncta per cell, normalized to the control variant (DMSO) ± SEM. *N* = 5 independent experiments. *n* = 173 cells (DMSO), 210 cells (RAPA), 172 cells (cycloheximide). \**p* < 0.05, *ns* – nonsignificant (one-way ANOVA followed by Bonferroni *post hoc* test).

**Fig. S4.**
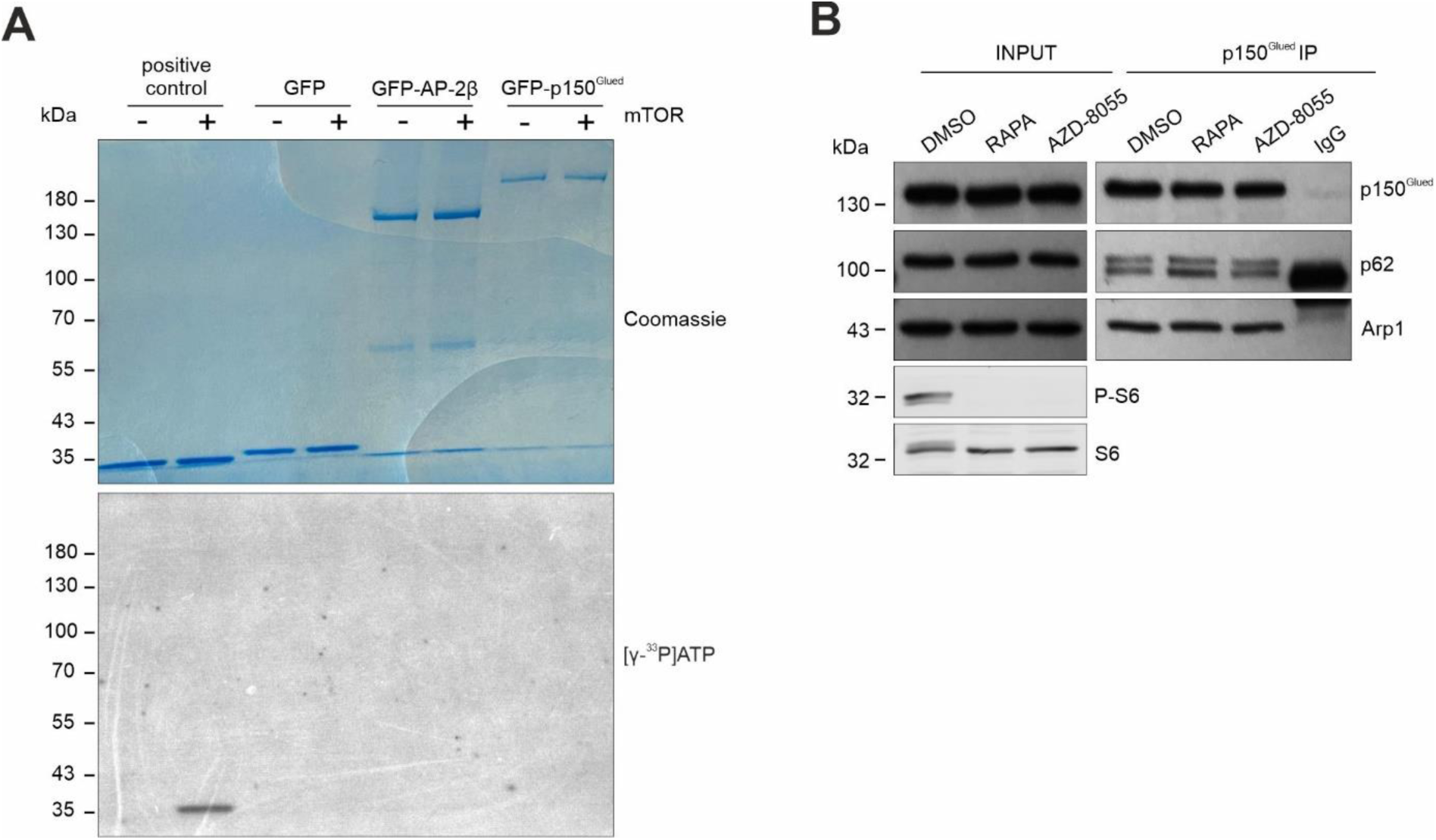
mTOR does not phosphorylate AP-2β or p150^Glued^ and does not affect p150^Glued^ binding to Arp1 or p62. (**A**) Results of kinase assay using recombinant active mTOR fragment and GFP-AP-2β, GFP-p150^Glued^, or GFP (negative control) immunoprecipitated from HEK293T cells as substrates. Commercial mTOR substrate served as a positive control. The upper panel shows the Coomassie staining of SDS-PAGE gels with the analyzed proteins. The lower panel shows the radioactive signal level of [γ-33P] ATP that was incorporated into the analyzed proteins. Shown is a representative example from *N* = 2 independent experiments. (**B**) Western blot analysis of endogenous p150^Glued^, Arp1, p62, S6, and P-S6 (Ser235/236) levels in HEK293T cells and co-immunoprecipitation of p150^Glued^, Arp1, and p62 from HEK293T cells that were treated for 2 h with 0.1% DMSO, 100 nM rapamycin (RAPA), or 100 nM AZD80550. Shown is a representative example from *N* = 2 independent experiments.

**Fig. S5.**
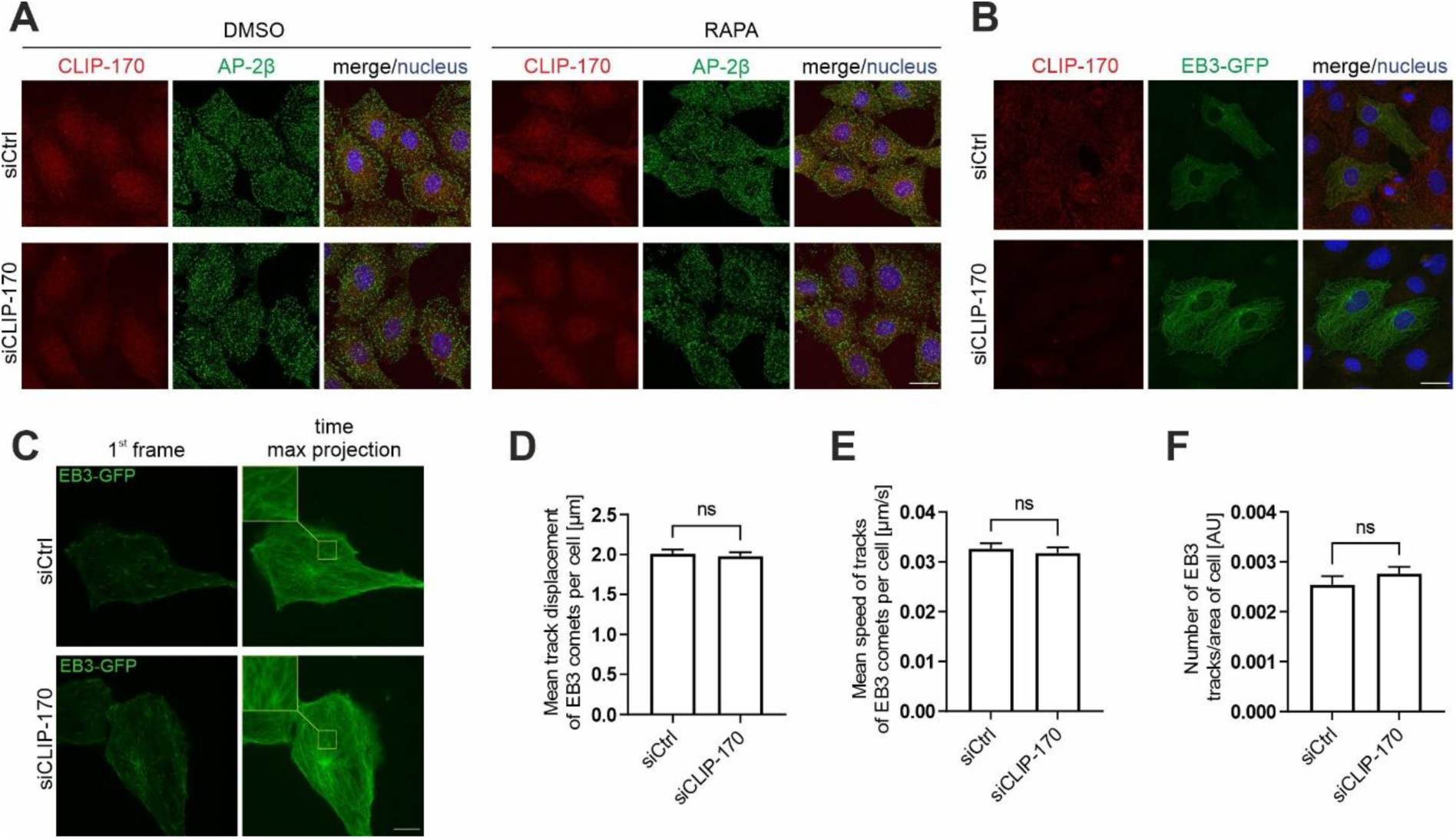
CLIP-170 knockdown does not influence AP-2β distribution or dynamic microtubules. (**A**) Representative images of fluorescently labeled endogenous CLIP-170 (red), AP-2β (green), and nuclei stained with Hoechst 33258 (blue) in Rat2 cells that were transfected with siCtrl or rat siCLIP-170 for 72 h, followed by treatment with 0.1% DMSO or 100 nM rapamycin for 2 h. Scale bar = 20 µm. (**B**) Representative images of fluorescently labeled endogenous CLIP-170 (green) and nuclei stained with Hoechst 33258 (blue) in Rat2 cells that were transfected with siCtrl or rat siCLIP-170 and 24 h later electroporated with a plasmid that encoded EB3-GFP. Cells were fixed 48 h after electroporation. Scale bars = 20 µm. (**C**) Representative images from time-lapse movies of microtubule dynamics in Rat2 cells that were transfected with siCtrl or siCLIP-170 and 24 h later electroporated with a plasmid that encoded EB3-GFP. The experiment was performed 48 h after electroporation. Images in the left column show the first frame from the time-lapse movies. Images in the right column show maximum intensity projections from Z-stacks of all 600 frames. Scale bar = 10 µm. (**D**) Quantification of the mean track run lengths of EB3-GFP comets per cell. The data are expressed as the mean ± SEM. *N* = 3 independent experiments. *n* = 26 cells (siCtrl), 26 cells (siCLIP-170). *ns*, nonsignificant (Student’s *t*-test). (**E**) Quantification of the mean speed of EB3-GFP comet. The data are expressed as the mean ± SEM. *N* = 3 independent experiments. *n* = 26 cells (siCtrl), 26 cells (siCLIP-170). *ns*, nonsignificant (Mann-Whitney test). (**F**) Quantification of the number of all microtubule tracks per cell divided by cell area for EB3-GFP. The data are expressed as the mean ± SEM. *N* = 3 independent experiments. *n* = 26 cells (siCtrl), *n* = 26 cells (siCLIP-170). *ns*, nonsignificant (Student’s *t*-test).

**Fig. S6.**
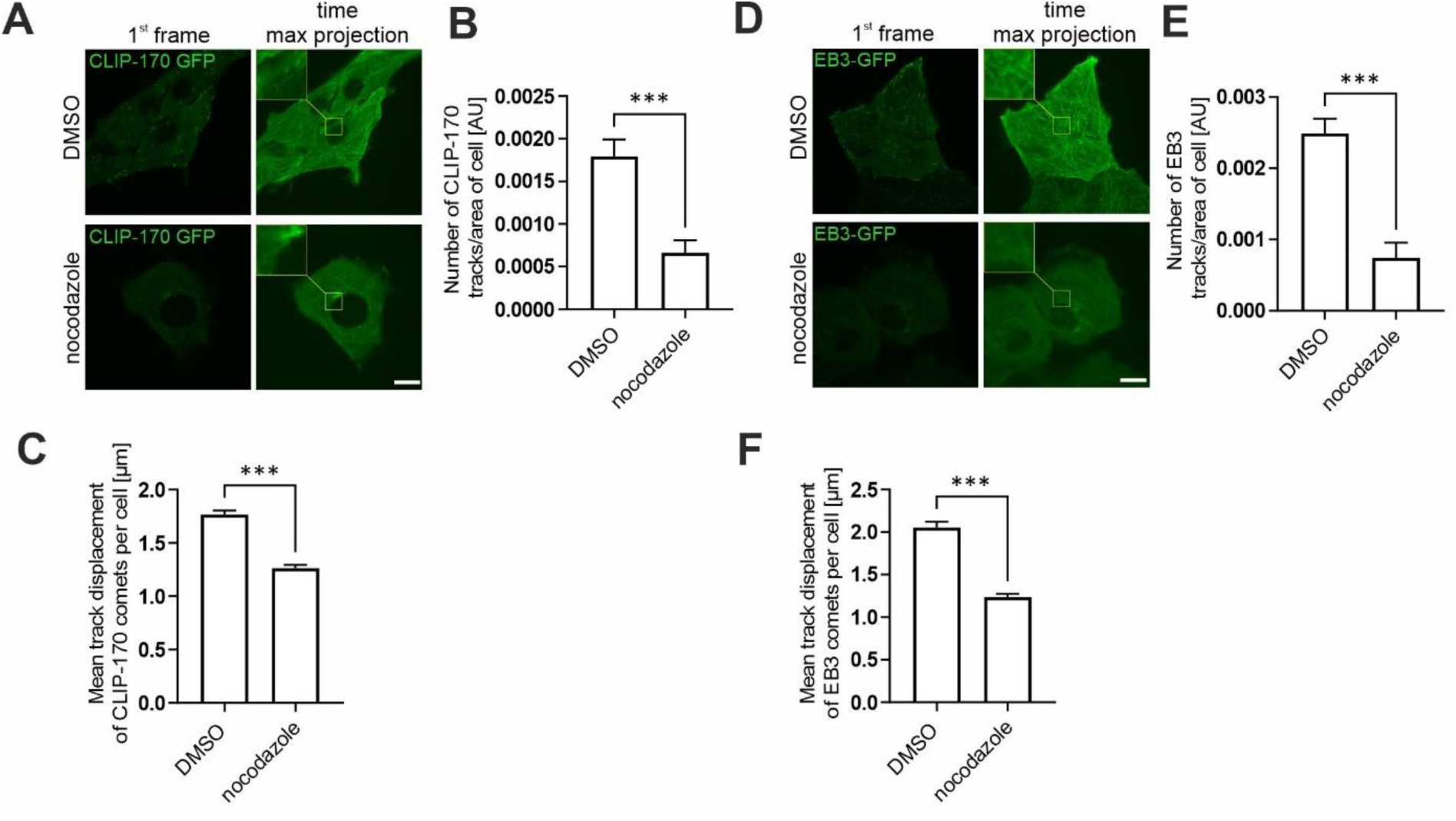
Low doses of nocodazole affect microtubule dynamics. (**A**) Representative images from time-lapse movies of microtubule dynamics in Rat2 cells, measured by microtubule plus-end tracking behavior of CLIP-170-GFP after 1 h treatment with 0.1% DMSO or 100 nM nocodazole. Images in the left column show the first frame from the time-lapse movies. Images in the right column show maximum intensity projections from Z-stacks of all 600 frames. Scale bar = 10 µm. (**B**) Quantification of the number of all microtubule tracks per cell divided by cell area for CLIP-170-GFP. The data are expressed as the mean ± SEM. *N* = 3 independent experiments. *n* = 31 cells (DMSO), 25 cells (nocodazole). *\*\*\*p* < 0.001 (Mann-Whitney test). (**C**) Quantification of the mean track run length of CLIP-170-GFP comets per cell. The data are expressed as the mean ± SEM. *N* = 3 independent experiments. *n* = 31 cells (DMSO), 25 cells (nocodazole). *\*\*\*p* < 0.001 (Mann-Whitney test). (**D**) Representative images from time-lapse movies of microtubule dynamics in Rat2 cells, measured by microtubule plus-end tracking behavior of EB3-GFP after 1 h treatment with 0.1% DMSO or 100 nM nocodazole. Images in the left column show the first frame from the time-lapse movies. Images in the right column show maximum intensity projections from Z-stacks of all 600 frames. Scale bar = 10 µm. (**E**) Quantification of the number of all microtubule tracks per cell divided by cell area for EB3-GFP. The data are expressed as the mean ± SEM. *N* = 3 independent experiments. *n* = 23 cells (DMSO), 24 cells (nocodazole). *\*\*\*p* < 0.001 (Mann-Whitney test). (**F**) Quantification of the mean track run lengths of EB3-GFP comets per cell. The data are expressed as the mean ± SEM. *N* = 3 independent experiments. *n* = 23 cells (DMSO), 24 cells (nocodazole). *\*\*\*p* < 0.001 (Mann-Whitney test).

**Fig. S7.**
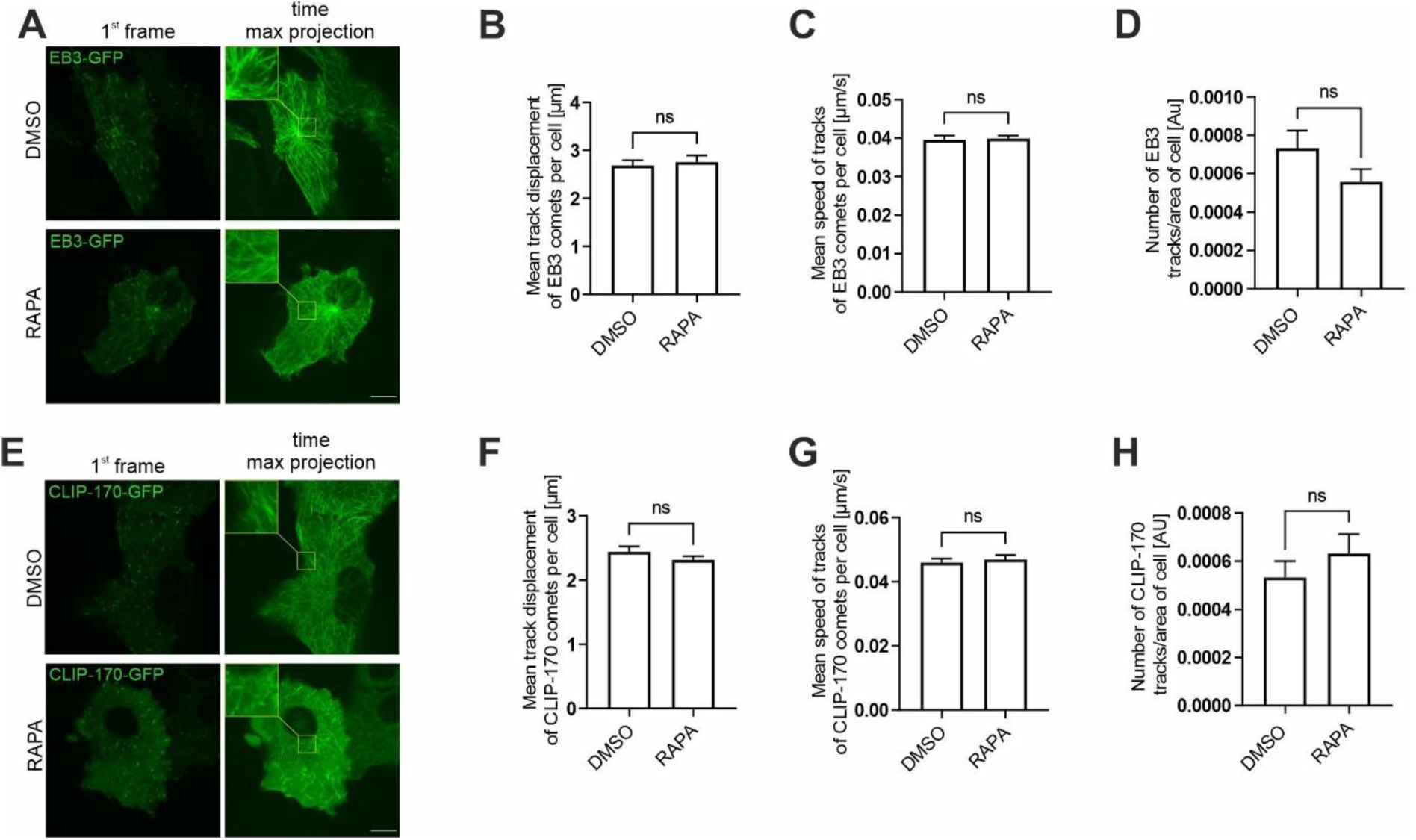
Rapamycin does not affect microtubule dynamics in Rat2 cells. (**A**) Representative images from time-lapse movies of microtubule dynamics in Rat2 cells, measured by microtubule plus-end tracking behavior of EB3-GFP after 2 h treatment with 0.1% DMSO or 100 nM rapamycin (RAPA). Images in the left column show the first frame from the time-lapse movies. Images in the right column show maximum intensity projections from Z-stacks of all 600 frames. Scale bar = 10 µm. (**B**) Quantification of the mean track run lengths of EB3-GFP comets per cell. The data are expressed as the mean ± SEM. *N* = 3 independent experiments. *N* = 27 cells (DMSO), 27 cells (RAPA). *ns*, nonsignificant (Mann-Whitney test). (**C**) Quantification of the mean speed of EB3-GFP comet. The data are expressed as the mean ± SEM. *N* = 3 independent experiments. *n* = 27 cells (DMSO), 27 cells (RAPA). *ns*, nonsignificant (Mann-Whitney test). (**D**) Quantification of the number of all microtubule tracks per cell divided by cell area for EB3-GFP. The data are expressed as the mean ± SEM. *N* = 3 independent experiments. *n* = 27 cells (DMSO), 27 cells (RAPA). *ns*, nonsignificant (Mann-Whitney test). (**E**) Representative images from time-lapse movies of microtubule dynamics in Rat2 cells, measured by microtubule plus-end tracking behavior of CLIP-170-GFP after 2 h treatment with 0.1% DMSO or 100 nM rapamycin (RAPA). Images in the left column show the first frame from the time-lapse movies. Images in the right column show maximum intensity projections from Z-stacks of all 600 frames. Scale bar = 10 µm. (**F**) Quantification of the mean track run lengths of CLIP-170-GFP comets per cell. The data are expressed as the mean ± SEM. *N* = 3 independent experiments. *n* = 35 cells (DMSO), 34 cells (RAPA). *ns*, nonsignificant (Mann-Whitney test). (**G**) Quantification of the mean speed of CLIP-170-GFP comet. The data are expressed as the mean ± SEM. *N* = 3 independent experiments. *n* = 35 cells (DMSO), 34 cells (RAPA). *ns*, nonsignificant (Mann-Whitney test). (**H**) Quantification of the number of all microtubule tracks per cell divided by cell area for CLIP-170-GFP. The data are expressed as the mean ± SEM. *N* = 3 independent experiments. *n* = 3*5* cells (DMSO), 34 cells (RAPA). *ns*, nonsignificant (Mann-Whitney test).

**Fig. S8.**
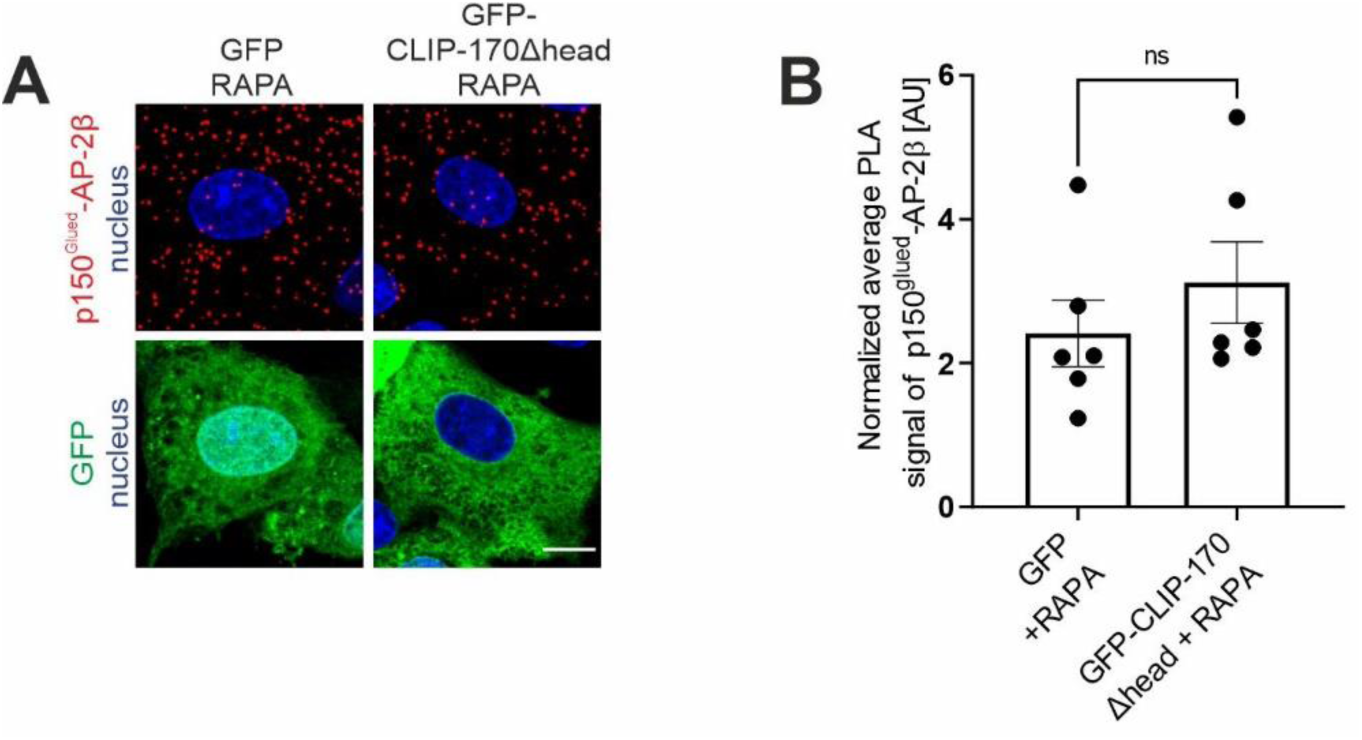
Overexpression of dominant-negative mutant of CLIP-170 does not impact p150^Glued^–AP-2β interaction. (**A**) Representative images of Rat2 fibroblasts that were transfected with pEGFPC1 or pEGFPC1-CLIP-170Δhead (green) and treated for 2 h with 100 nM rapamycin (RAPA) with PLA p150^Glued^–AP-2β signals (red) and DAPI-stained nuclei (blue). Scale bar = 10 μm. (**B**) Quantification of the number of p150^Glued^–AP-2β PLA puncta in cells that were treated as in A. The data are expressed as the mean number of PLA puncta per cell, normalized to the control variant (GFP + DMSO; not shown) ± SEM. *N* = 6 independent experiments. *n* = 94 cells (GFP + RAPA), 114 cells (GFP-CLIP-170Δhead + RAPA). *ns*, nonsignificant (Mann-Whitney test).

**Fig. S9.**
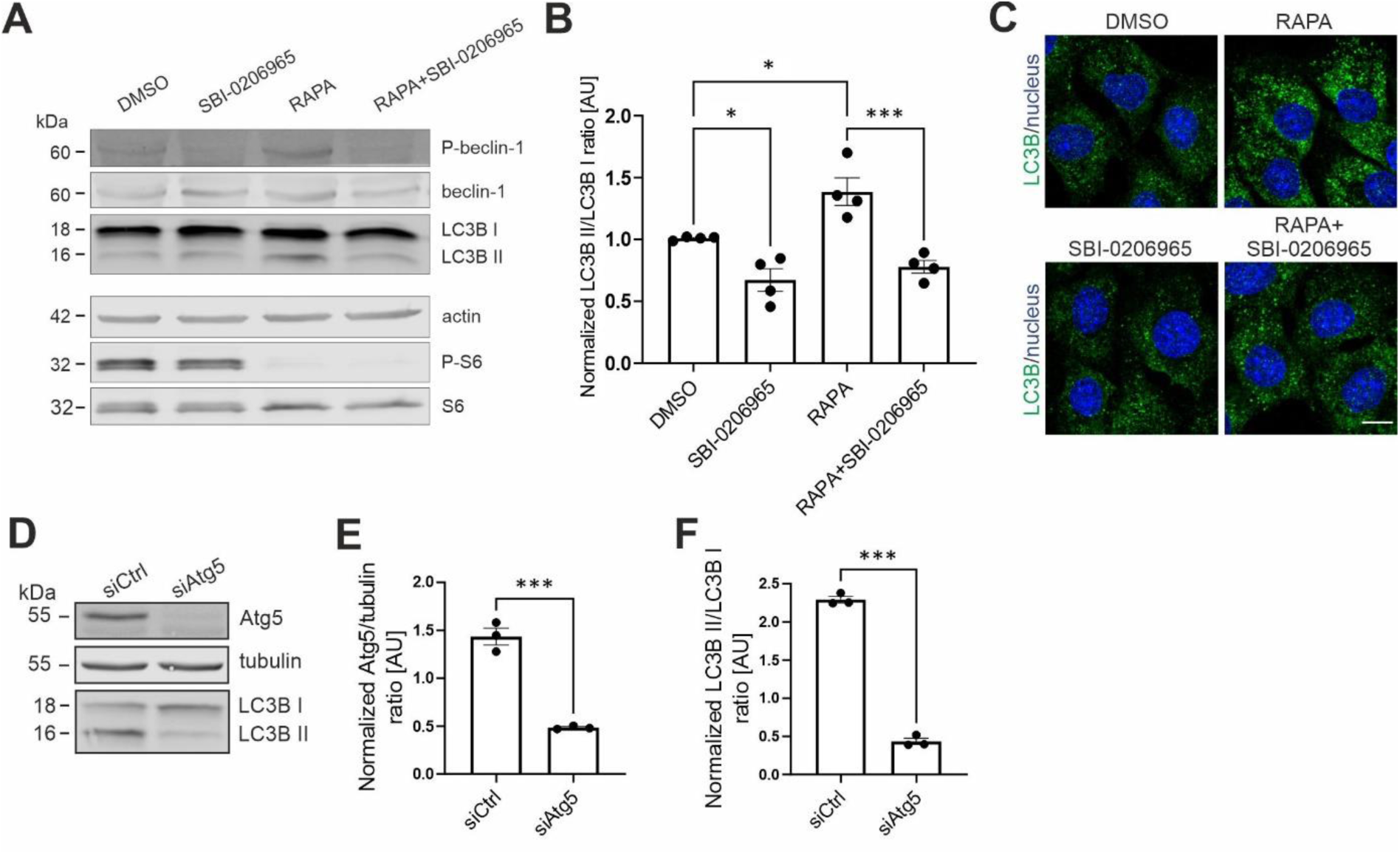
Confirmation of effects of rapamycin, SBI-0206965, and Atg5 knockdown on autophagy. (**A**) Western blot analysis of endogenous P-beclin-1 (Ser30), beclin-1, LC3B I, LC3B II, actin, P-S6 (Ser235/236), and S6 levels in Rat2 fibroblasts that were treated with 0.1% DMSO for 2 h, 100 nM rapamycin (RAPA) for 2 h, 25 μM SBI-0206965 for 2 h 30 min, or 25 μM SBI-0206965 for 30 min and 100 nM rapamycin for 2 h (RAPA + SBI-0206965). (**B**) Densitometry analysis of normalized LC3B II/LCB I ratio in Rat2 cells that were treated as in A. The data are presented as mean of the normalized ratio of LC3B II to LC3B I levels ± SEM. *N* = 4 independent experiments. *\*p <* 0.05, *\*\*\*p* < 0.001 (one-way ANOVA followed by Bonferroni multiple-comparison *post hoc* test). (**C**) Representative images of Rat2 cells that were treated with 0.1% DMSO for 2 h, 100 nM rapamycin (RAPA) for 2 h, 25 μM SBI-0206965 for 2 h 30 min, or 25 μM SBI-0206965 for 30 min and 100 nM rapamycin for 2 h (RAPA + SBI-0206965), with immunofluorescently labeled endogenous LC3B (green) and nuclei that were stained with Hoechst 33258 (blue). Scale bar = 10 µm. (**D**) Western blot analysis of endogenous Atg5, LC3B I, LC3B II, and tubulin in Rat2 cells that were transfected with siCtrl or rat siAtg5 for 72 h. (**E**) Densitometry analysis of normalized Atg5 in Rat2 cells that were treated as in D. The data are presented as mean of the normalized ratio of Atg5 to tubulin levels ± SEM. *N* = 3 independent experiments. *\*\*\*p* < 0.001 (Student’s *t*-test). (**F**) Densitometry analysis of normalized LC3B II/LCB I ratio in Rat2 cells that were treated as in D. The data are presented as mean of the normalized ratio of LC3B II to LC3B I levels ± SEM. *N* = 3 independent experiments. *\*\*\*p* < 0.001 (Student’s *t*-test).

**Fig. S10.**
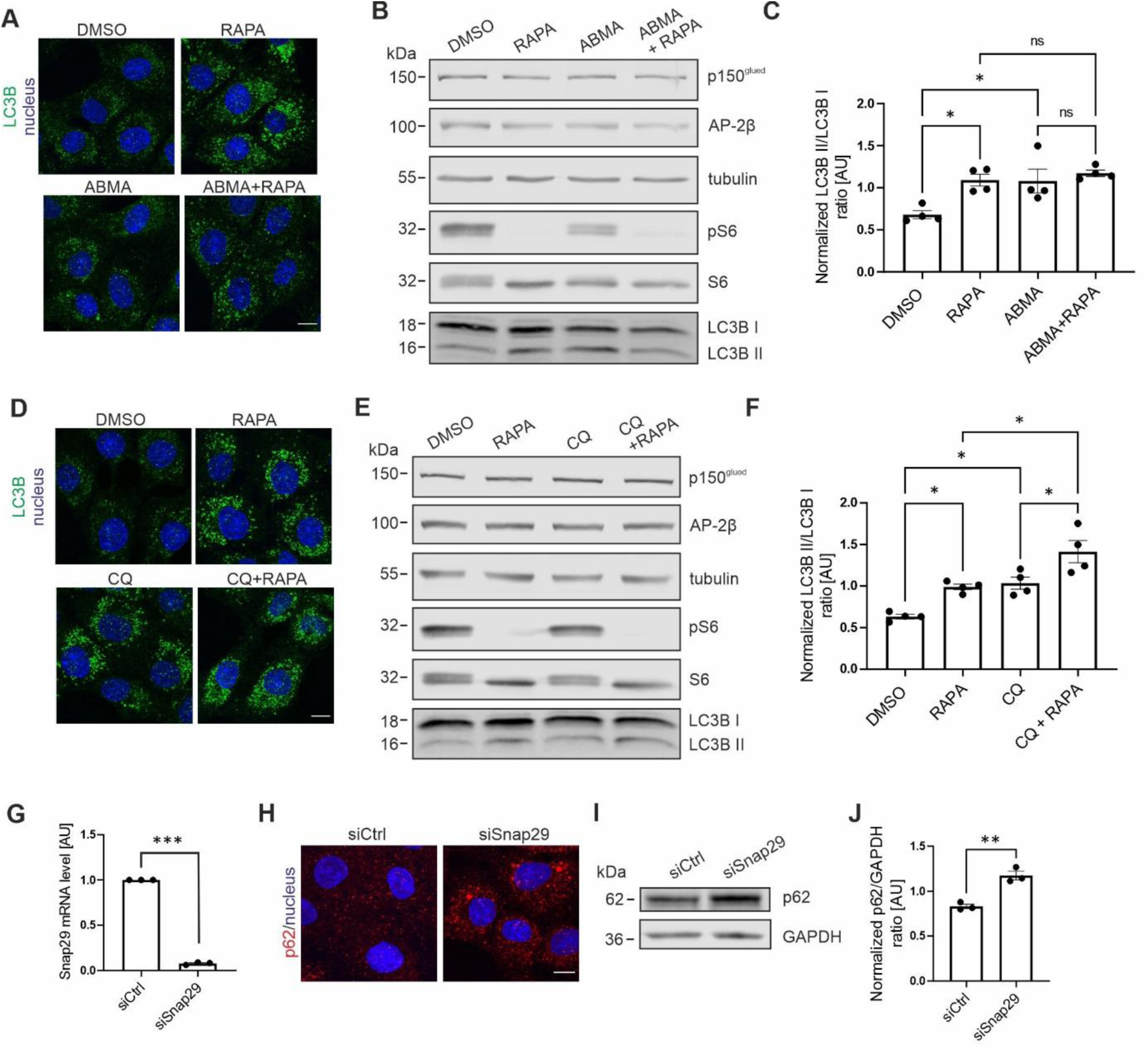
Confirmation of effects of ABMA, CQ, and SNAP29 knockdown on autolysosomal pathway. (**A**) Representative images of Rat2 cells that were treated for 2 h with 0.1% DMSO, 100 nM rapamycin (RAPA), 60 μM ABMA (ABMA), or 60 μM ABMA and 100 nM rapamycin (ABMA+RAPA), with immunofluorescently labeled endogenous LC3B (green) and nuclei that were stained with Hoechst 33258 (blue). Scale bar = 10 µm. (**B**) Western blot analysis of endogenous proteins levels as indicated in Rat2 cells that were treated as in A. (**C**) Densitometry analysis of normalized LC3B II/LCB I ratio in Rat2 cells that were treated as in A. The data are presented as mean of the normalized ratio of LC3B II to LC3B I levels ± SEM. *N* = 4 independent experiments. *\*p <* 0.05, *ns* – nonsignificant (one-way ANOVA followed by Bonferroni *post hoc* test). (**D**) Representative images of Rat2 cells that were treated for 2 h with 0.1% DMSO, 100 nM rapamycin (RAPA), 50 μM chloroquine (CQ), or 50 μM chloroquine and 100 nM rapamycin (CQ+RAPA), with immunofluorescently labeled endogenous LC3B (green) and nuclei that were stained with Hoechst 33258 (blue). Scale bar = 10 µm. (**E**) Western blot analysis of endogenous protein levels as indicated in Rat2 cells that were treated as in D. (**F**) Densitometry analysis of normalized LC3B II/LCB I ratio in Rat2 cells that were treated as in D. The data are presented as mean of the normalized ratio of LC3B II to LC3B I levels ± SEM. *N* = 4 independent experiments. *\*p <* 0.05 (one-way ANOVA followed by Bonferroni *post hoc* test). (**G**) Results of qRT-PCR analysis of Snap29 mRNA levels relative to tubulin mRNA in Rat2 cells that were transfected with control siRNA (siCtrl) or siRNA against Snap29 (siSnap29) for 72 h. The results are expressed as means, normalized to control ± SEM. *N* = 3 independent experiments. \*\*\**p* < 0.001 (one-sample *t*-test). (**H**) Representative images of Rat2 cells that were treated as in G, with immunofluorescently labeled endogenous p62/SQSTM1 (red) and nuclei that were stained with Hoechst 33258 (blue). Scale bar = 10 µm. (**I**) Western blot analysis of endogenous protein levels as indicated in Rat2 cells that were treated as in G. (**J**) Densitometry analysis of normalized p62/SQSTM1 in Rat2 cells that were treated as in G. The data are presented as mean of the normalized ratio of p62 to tubulin levels ± SEM. *N* = 3 independent experiments. *\*\*p* < 0.01 (Student’s *t*-test).

**Fig. S11.**
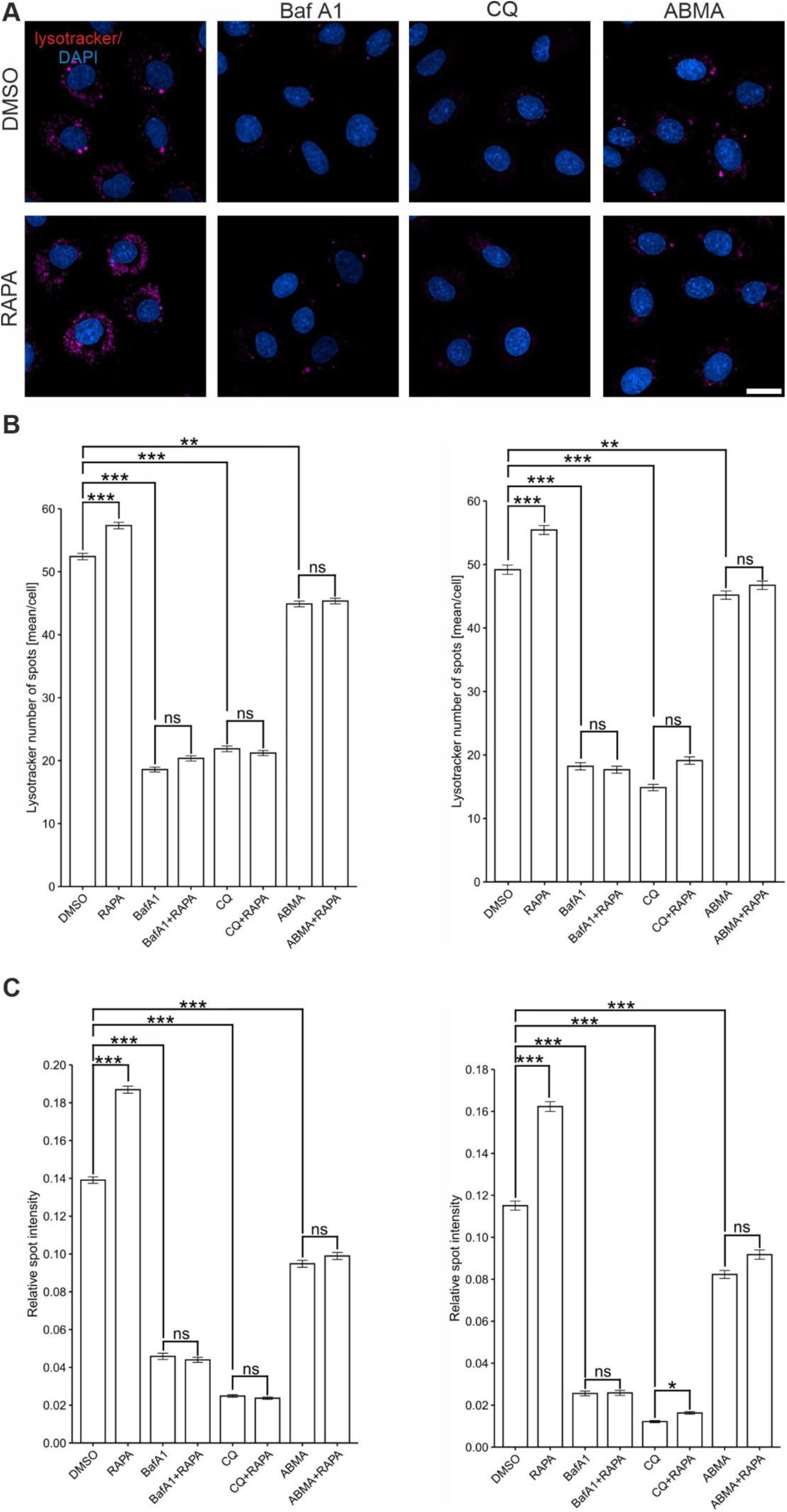
(**A**) Representative images of Rat2 cells that were treated for 2 h as indicated, with immunofluorescently labeled Lysotracker (magenta) and nuclei that were stained with Hoechst 33258 (blue). Scale bar = 20 µm. (**B, C**) Results of two independent repetitions of high-throughput microscopy analyses of Lysotracker mean number of spots per cell and relative spot intensity in cells that were treated with 0.1% DMSO, 100 nM rapamycin (RAPA), 100 nM bafilomycin A1 (BafA1), 50 μM chloroquine (CQ), 60 μM ABMA (ABMA), or a combination of 100 nM rapamycin with 100 nM bafilomycin A1 (BafA1 + RAPA), 50 μM chloroquine (CQ + RAPA), or 60 μM ABMA (ABMA + RAPA). The results are expressed as the mean ± SEM. *N* = 2 independent experiments. *n* = 913 and 689 cells analyzed in each group for repetition 1 and 2. \**p* < 0.05, \*\**p* < 0.01, **8*p* < 0.001, *ns* – nonsignificant (Kruskal-Wallis test followed by Dunn’s *post hoc* test).

**Fig. S12.**
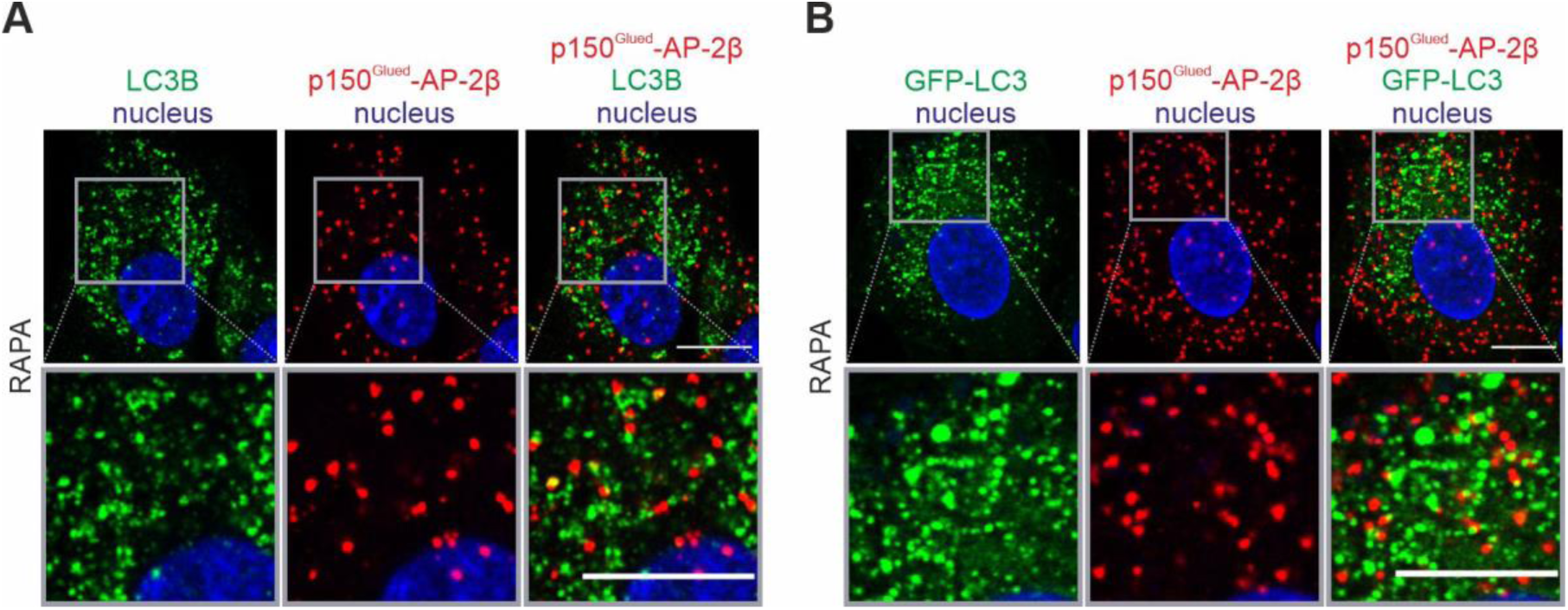
p150^Glued^–AP-2β PLA signal co-localizes poorly with LC3. (**A**) Representative images of Rat2 cells treated with 100 nM rapamycin (RAPA) with PLA p150^Glued^–AP-2β signals (red) immunofluorescently labeled for endogenous LC3B (green) and nuclei stained with DAPI (blue). Images were acquired using the AiryScan module. Scale = 10 μm. (Upper panel) Representative photograph of a single cell. (Lower panel) Close-up of a site with PLA and LC3B signals. (**B**) Representative images of Rat2 cells transfected with GFP-LC3 for 24 h and then treated with 100 nM rapamycin (RAPA) with PLA p150^Glued^–AP-2β signals (red), overproduced GFP-LC3 (green), and nuclei stained with DAPI (blue) and. Images were acquired using the AiryScan module. Scale = 10 μm. (Top) Representative photograph of a single cell. (Bottom) Close-up of a site with PLA and GFP-LC3B signals.

**Fig. S13.**
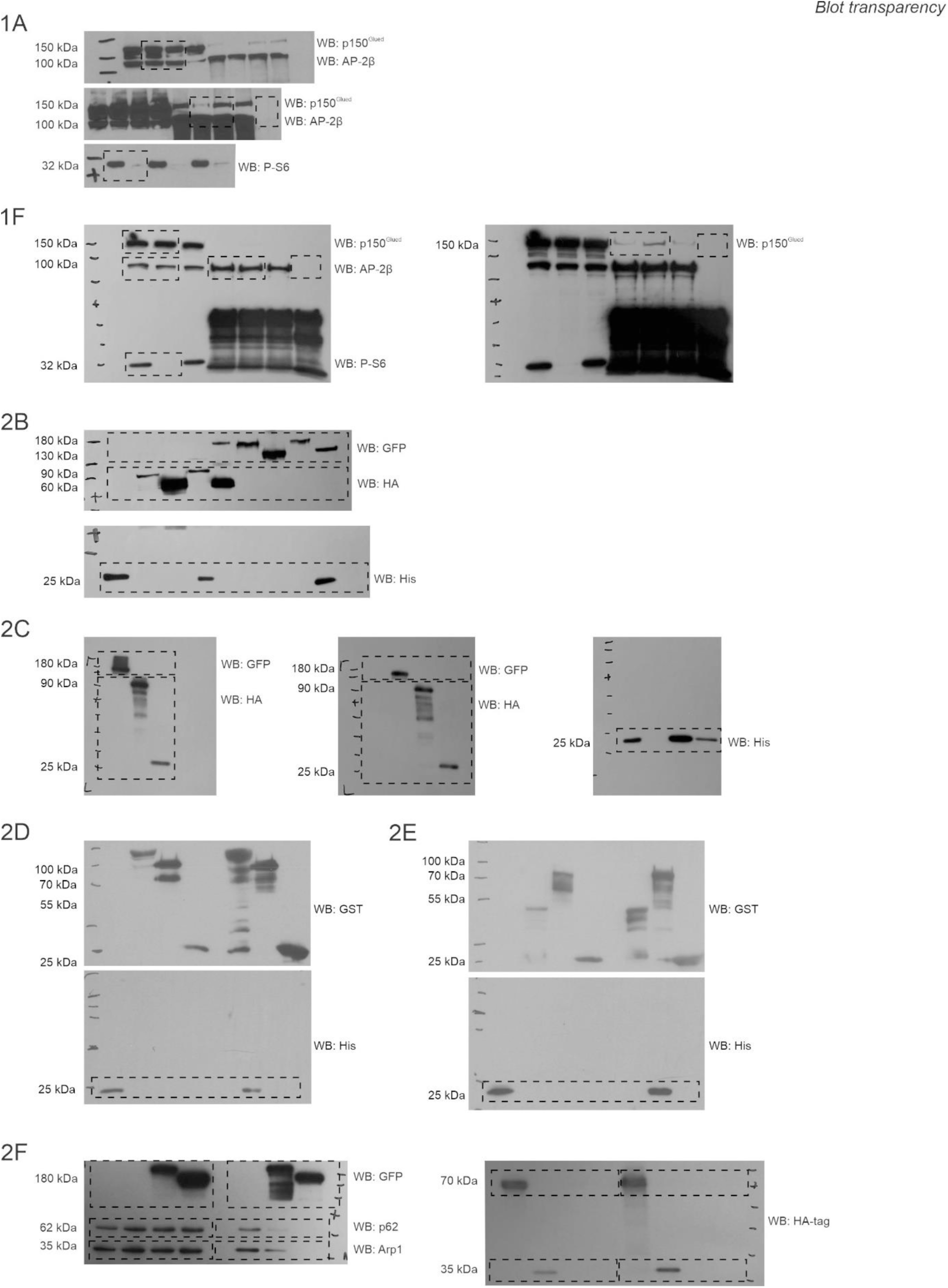
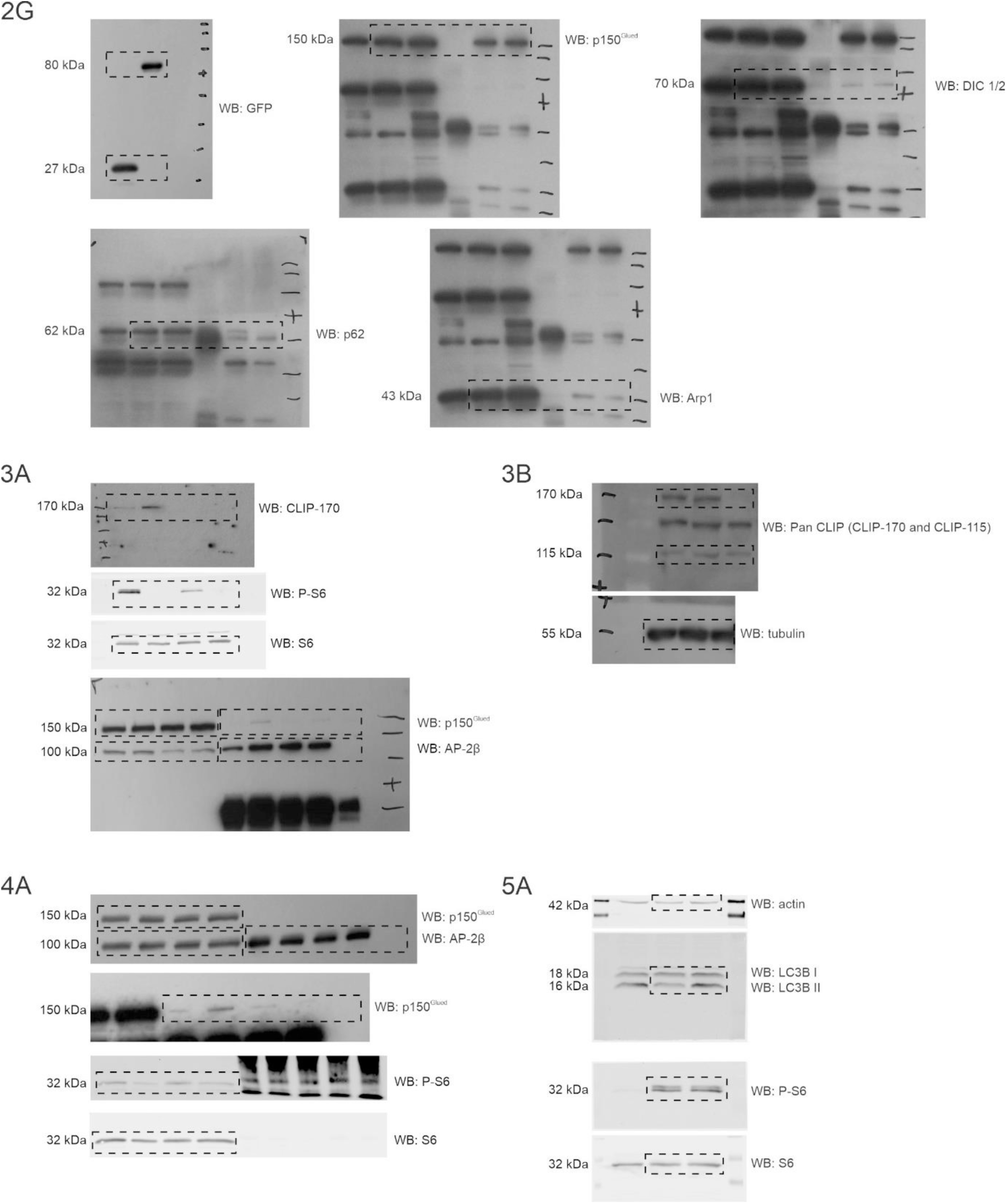
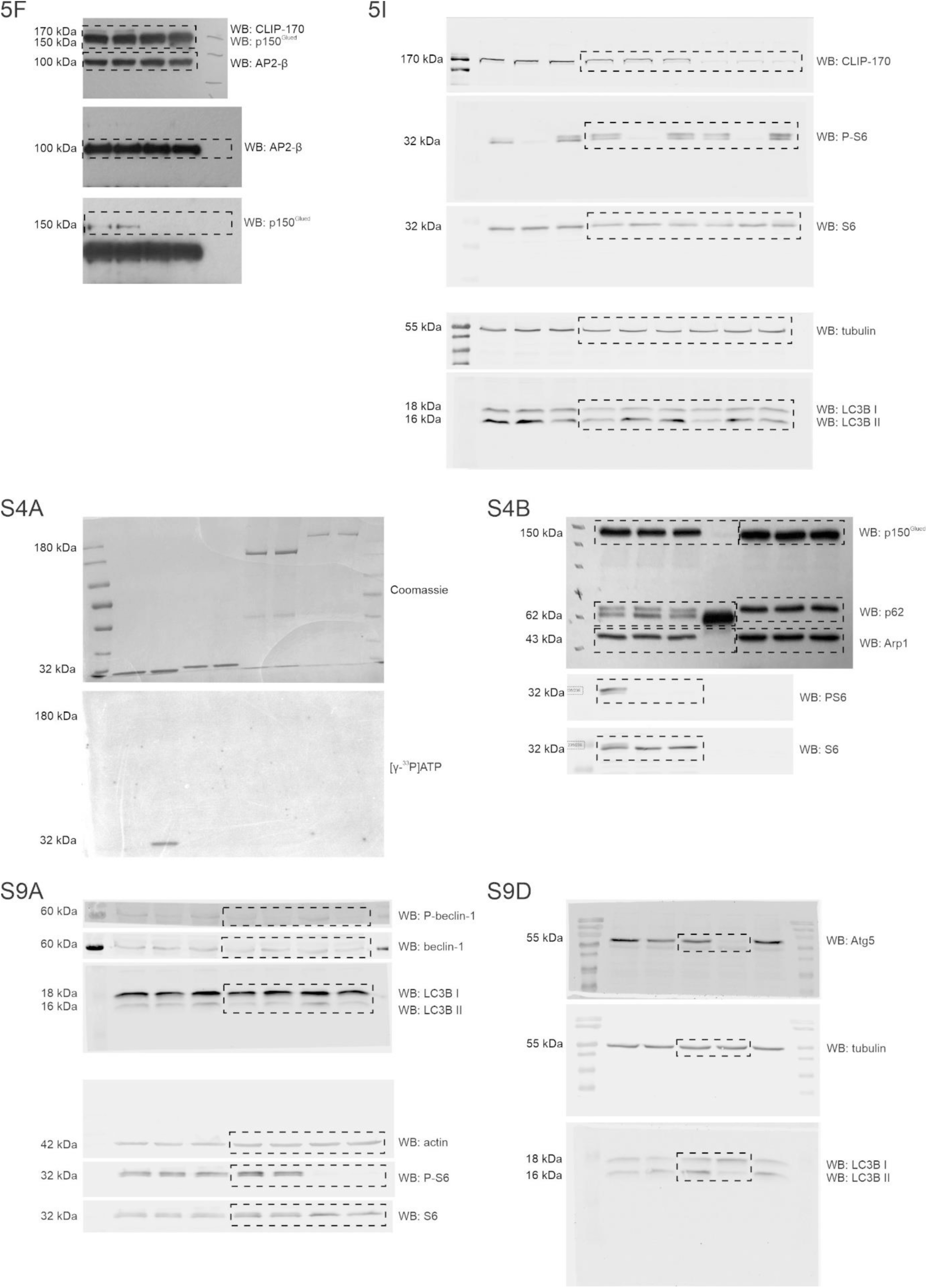
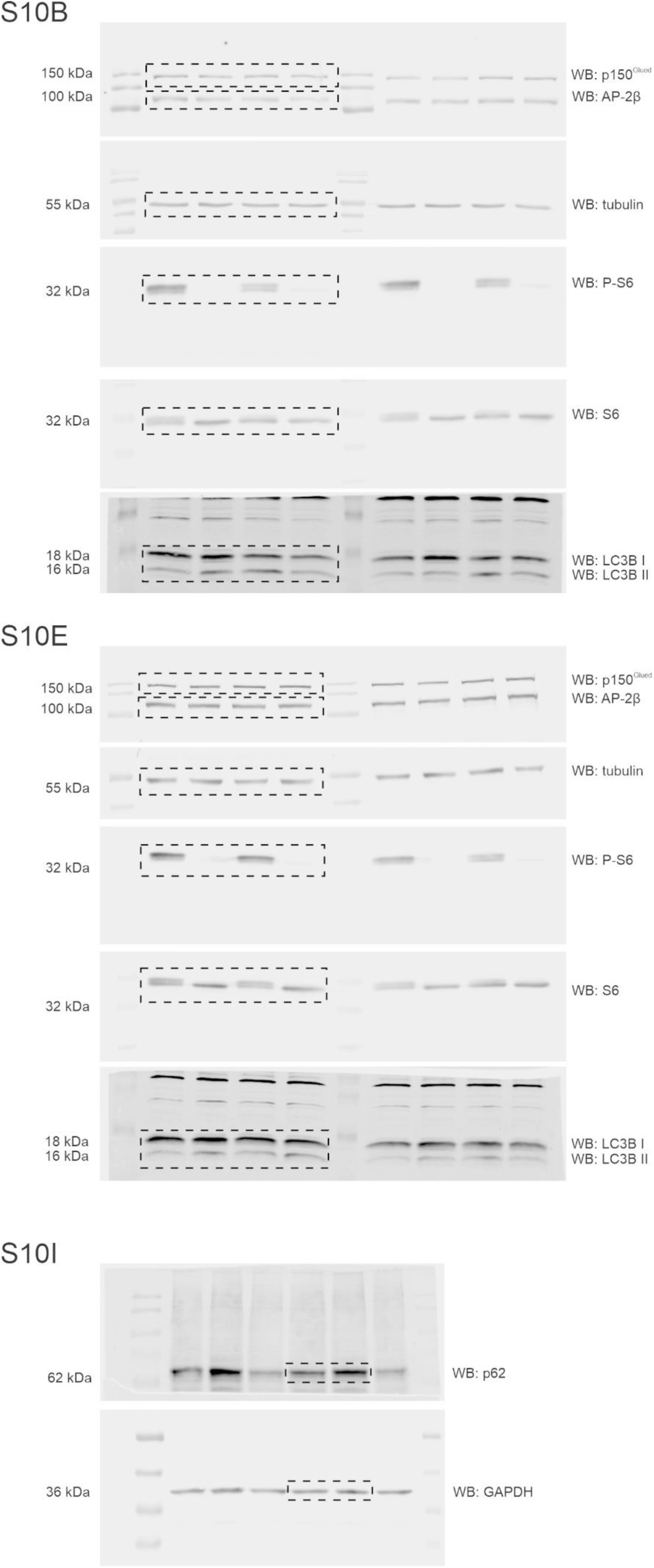
Blot transparency. Full-size blots that correspond to cropped images that are presented in the manuscript.

## Movies

**Movie 1.**
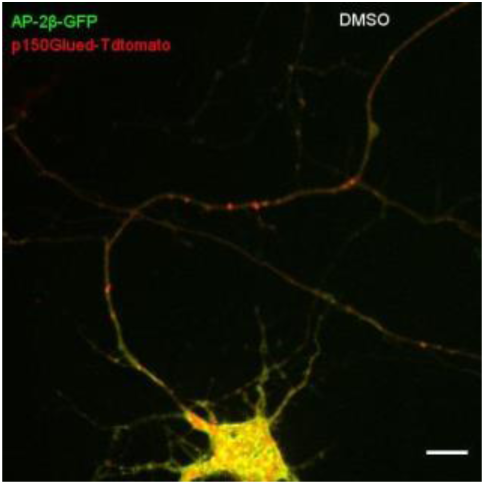
Hippocampal neuron that was transfected with plasmids that encoded p150^Glued^-Tdtomato (red) and AP-2β-GFP (green) and treated for 2 h with 0.1% DMSO. Speed = 12× real time. Scale bar = 10 µm.

**Movie 2.**
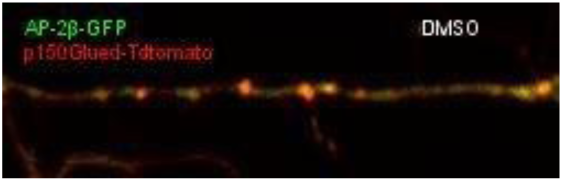
Straightened fragment of axon of neuron from Movie 1. Speed = 12× real time. Scale bar = 10 µm.

**Movie 3.**
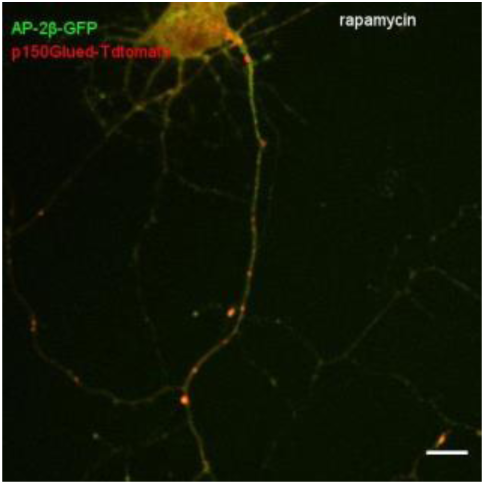
Hippocampal neuron that was transfected with plasmids that encoded p150^Glued^-Tdtomato and AP-2β-GFP and treated for 2 h with 100 nM rapamycin. Speed = 12× real time. Scale bar = 10 µm.

**Movie 4.**
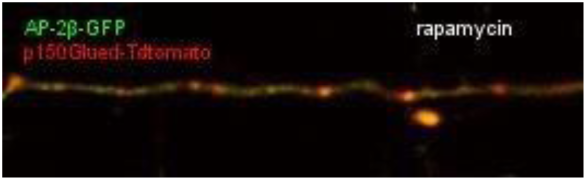
Straightened fragment of axon of neuron from Movie 3. Speed = 12× real time. Scale bar = 10 µm.

**Movie 5.**
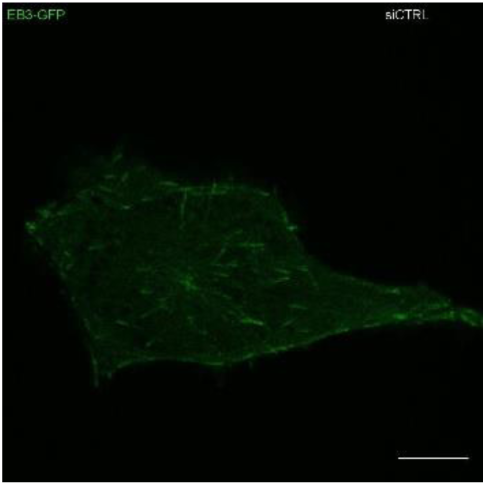
Rat2 cells that were transfected with siCtrl and 24 h later electroporated with a plasmid that encoded EB3-GFP. Speed = 10× real time. Scale bar = 10 µm.

**Movie 6.**
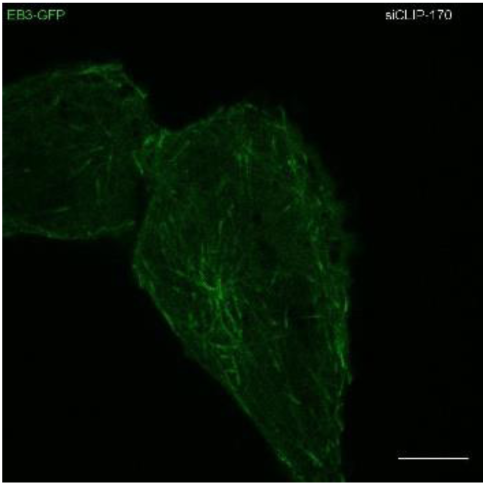
Rat2 cells that were transfected with siCLIP-170 and 24 h later electroporated with a plasmid that encoded EB3-GFP. Speed = 10× real time. Scale bar = 10 µm.

**Movie 7.**
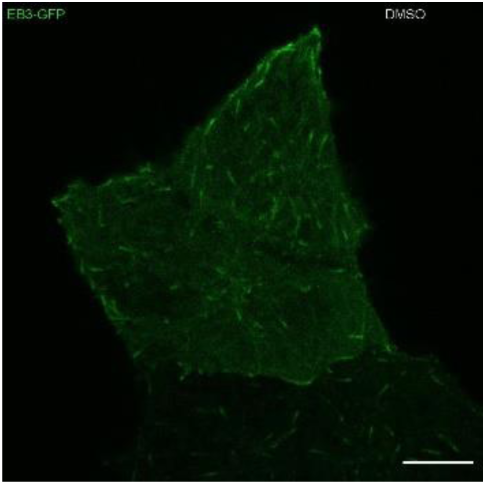
Rat2 cells that were electroporated with a plasmid that encoded EB3-GFP and treated with 0.1% DMSO for 1 h. Speed = 10× real time. Scale bar = 10 µm.

**Movie 8.**
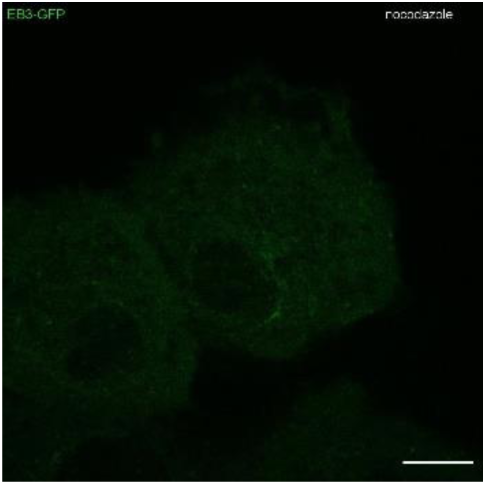
Rat2 cells that were electroporated with a plasmid that encoded EB3-GFP and treated with 100 nM nocodazole for 1 h. Speed = 10× real time. Scale bar = 10 µm.

**Movie 9.**
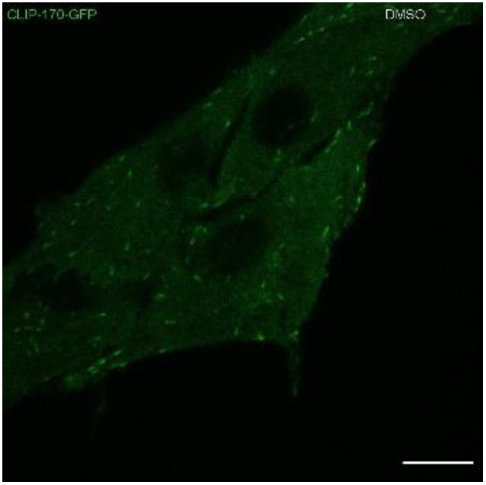
Rat2 cells that were electroporated with a plasmid that encoded CLIP-170-GFP and treated with 0.1% DMSO for 1 h. Speed = 10× real time. Scale bar = 10 µm.

**Movie 10.**
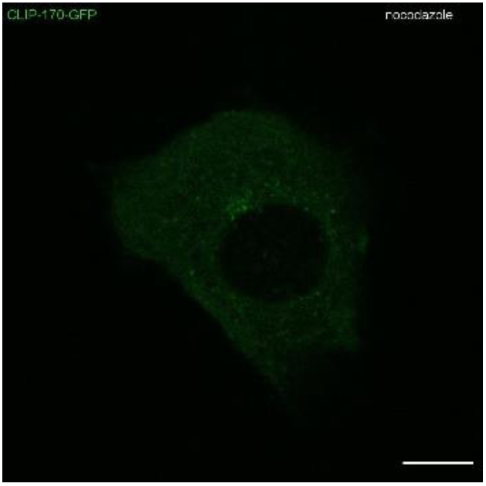
Rat2 cells that were electroporated with a plasmid that encoded CLIP-170-GFP and treated with 100 nM nocodazole for 1 h. Speed = 10× real time. Scale bar = 10 µm.

**Movie 11.**
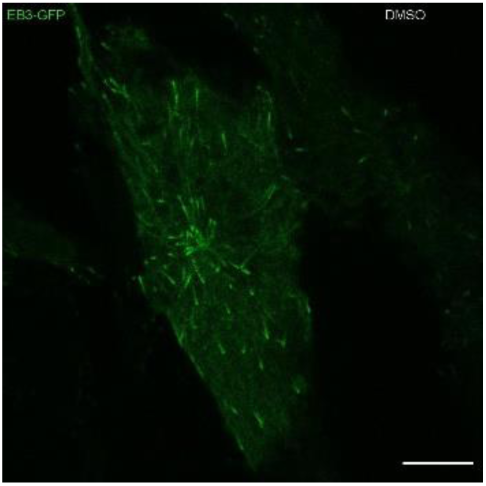
Rat2 cells that were electroporated with a plasmid that encoded EB3-GFP and treated with 0.1% DMSO for 2 h. Speed = 10× real time. Scale bar = 10 µm.

**Movie 12.**
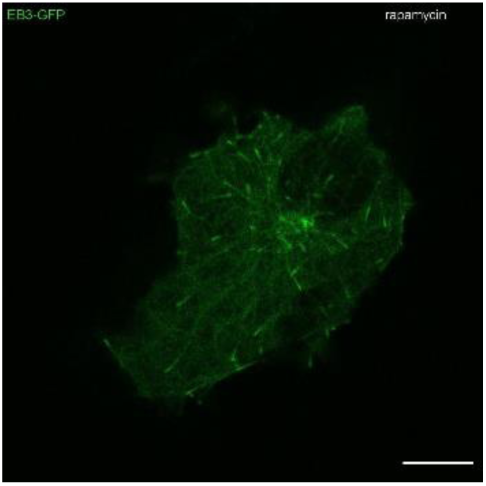
Rat2 cells that were electroporated with a plasmid that encoded EB3-GFP and treated with 100 nM rapamycin for 2 h. Speed = 10× real time. Scale bar = 10 µm.

**Movie 13.**
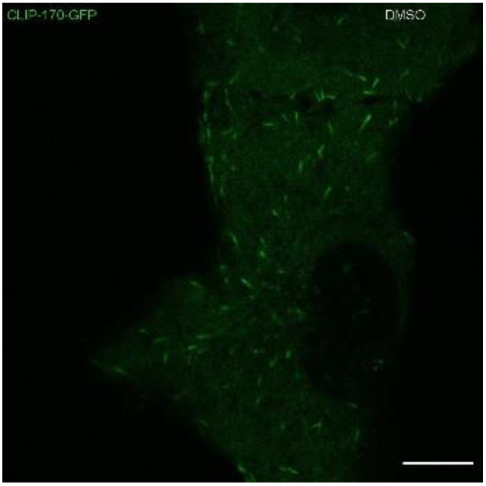
Rat2 cells that were electroporated with a plasmid that encoded CLIP-170-GFP and treated with 0.1% DMSO for 2 h. Speed = 10× real time. Scale bar = 10 µm.

**Movie 14.**
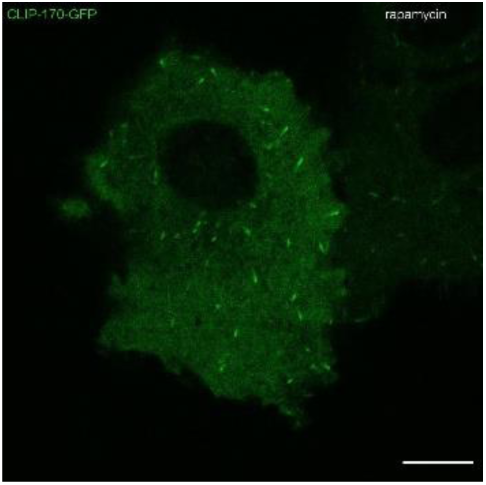
Rat2 cells that were electroporated with a plasmid that encoded CLIP-170-GFP and treated with 100 nM rapamycin for 2 h. Speed = 10× real time. Scale bar = 10 µm.

**Movie 15.**
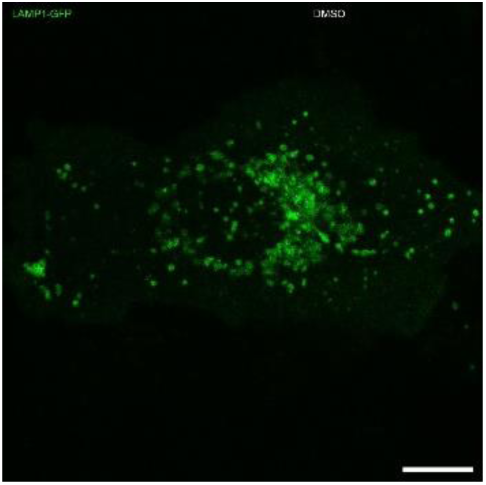
Rat2 cells that were electroporated with a plasmid that encoded LAMP1-GFP and treated with 100nM rapamycin for 2 h. Speed = 10x real time. Scale bar = 10 µm.

**Movie 16.**
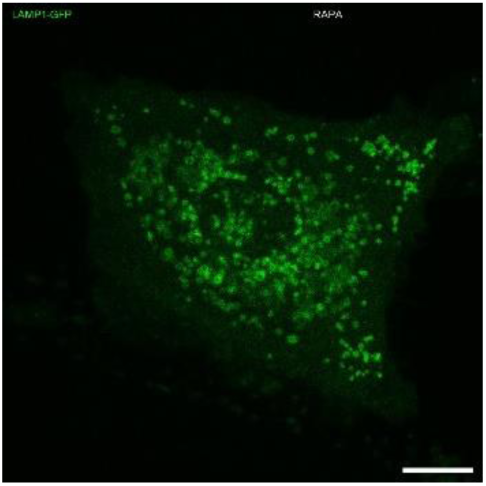
Rat2 cells that were electroporated with a plasmid that encoded LAMP1-GFP and treated with 0.1% DMSO for 2 h. Speed = 10x real time. Scale bar = 10 µm.

